# Evaluating camera trap methods for monitoring population trends in ungulates: insights from simulation

**DOI:** 10.1101/2025.03.20.644315

**Authors:** Clément Calenge, Sonia Saïd, Jules Chiffard, Mathieu Garel, Maryline Pellerin

**Affiliations:** Office Français de la Biodiversité – Direction Surveillance, Évaluation, Données – Unité Données et Appui Méthodologique, Saint Benoist, BP 20. 78612 Le Perray en Yvelines, France.; Office Français de la Biodiversité – Direction Recherche et Appui Scientifique - Service Conservation et gestion des espèces à enjeux, Birieux, France.; Office Français de la Biodiversité – Direction Recherche et Appui Scientifique - Service SantéAgri, Villeneuve de Rivière, France.; Office Français de la Biodiversité – Direction Recherche et Appui Scientifique - Service Anthropisation et fonctionnement des écosystèmes terrestres, Gières, France.; Office Français de la Biodiversité – Direction Recherche et Appui Scientifique - Service Conservation et gestion durable des espèces exploitées, Chateauvillain, France.

**Keywords:** random encounter models, instantaneous sampling, simulations, animal movement, habitat selection, sampling design

## Abstract

Camera traps have been widely used in the last decade to monitor abundance of unmarked animal populations. Most estimation methods rely either on the number of times animals pass through the detection zones, like random encounter models (REM) or on the number of capture occasions in a time-lapse program when animals were seen on the pictures, like the instantaneous sampling approach (IS). Yet, the ability of these two popular method classes to both reliably detect population trends and estimate population size has rarely been evaluated. We filled this gap by simulating a setup of either 100 or 25 camera traps randomly distributed on a 2600-ha area (respectively ≈ 4 and 1 trap/km^2^), along with the movements of a fictional population of 300 roe deer (*Capreolus capreolus*). Simulations were informed by field data on habitat, habitat selection and activity patterns of GPS-monitored roe deer. Under idealized conditions (e.g., perfect knowledge of day range and visibility), both IS and REM provided unbiased population estimates, though uncertainty remained substantial (CV from 15% to 30% with 4 and 1 trap/km^2^ respectively). However, our results show that neglecting imperfect detectability leads to severe biases in absolute density estimation. Moreover, despite idealized conditions and large sampling efforts, a simulated 20% population decline over 5 years went undetected by both approaches in 65-75% of simulations at high trap density and 80% at low trap density. Testing other sampling strategies to improve sensitivity either led to an unchanged population size estimation precision (stratified sampling) or to biased estimated trends (sampling only in high-quality habitats). Simulating animals with a 10 times larger home-range, led to miss the decline less frequently (5% – 40% at high trap density, 33% – 67% at low trap density). These results suggest that the key metric for camera trap use is the average number of different traps visited per animal, which in turn depends on trap density, home-range size and space use heterogeneity. We provide a R package allowing the reader to reproduce these simulations, and carry out their own.

## Introduction

Camera traps are used to reach many goals in wildlife studies, including estimating occupancy, studying animal behaviour, and investigating species interactions, in particular predator-prey interactions (see the review by Burton et al., 2015). Because in some species individual identification is possible based on phenotypic patterns, estimating the size of populations with capture-recapture methods has also been among the first use of camera traps, especially in elusive carnivores (Karanth, 1995). More recently, a large amount of literature has been devoted to the use of camera traps to estimate the size of the population of non individually identifiable animals (Rowcliffe et al., 2008; Nakashima et al., 2017; Howe et al., 2017; Moeller et al., 2018). The basic idea of these methods is that the number of animals detected by the traps can be used to infer the number of animals present in the study area.

Two main families of methods have been proposed to achieve this aim: those based on animal-trap encounters, which quantify the number of animals passages through the detection zones of trap motion sensors, and those based on animal-trap associations, which count animals at capture occasions within time-lapse programs (Campos-Candela et al., 2018). The random encounter model (REM, Rowcliffe et al., 2008) is a prominent example of the first family, while camera trap distance sampling (Howe et al., 2017), instantaneous sampling (IS, Moeller et al., 2018) or the similar “association model” of Campos-Candela et al. (2018) exemplify the second.

These estimation methods are often seen as a grail, in that they allow the estimation of population abundance, a key population parameter (Williams et al., 2002) without the intensive effort of capture-recapture. This is particularly appealing to many practitioners (Gilbert et al., 2020), who often rely on less burdensome index-based methods that only allow for trend estimation (Morellet et al., 2007; Garel et al., 2010). Consequently, some even propose replacing their established monitoring methods (whether capture or index-based) with camera trap-based approaches, driven by the perception that they are easier to implement and will facilitate both abundance and trend estimation (pers. obs.). This might be a problem, as the properties of these new methods are not yet fully understood, and numerous factors must be considered when using them (Calenge, 2026).

Thus, one major issue when estimating a population size is the imperfect detectability of the animals in the detection zone (Gilbert et al., 2020). Most camera trap studies rely on the use of motion sensors to trigger the traps when an animal enters the detection zone. However, such sensors are less sensitive when the animals are small (Glen et al., 2013) or far from the trap (Howe et al., 2017). The camera traps characteristics (brand, model, wear, etc.) may also have an effect on the detection probability of the animals in the detection zone (Palencia et al., 2021). Obstacles and environmental conditions may further limit its effective surface area (Palencia et al., 2021; Moeller et al., 2023). Numerous authors have insisted on the need to account for this imperfect detection in population size estimation (e.g., Howe et al., 2017; Moeller et al., 2023). However, most practitioners continue to ignore this issue, leading to potential biases in their estimates (Burton et al., 2015). Nevertheless, population trend estimates may still be unbiased, unless habitat structure changes over time (e.g. vegetation growth or human disturbance, Guillera-Arroita, 2017).

An issue tied to the broader problem of imperfect detection is the existence of periods during which animals are inactive and less detectable by motion sensors. Such periods do not necessarily affect estimation accuracy if the camera traps are randomly distributed over the study area, and if the resting areas have the same probability to be monitored as other areas (i.e., no refuges where the animals are undetectable, Rowcliffe et al., 2014). However, even when these assumptions are satisfied, the decreased detectability of inactive animals by motion sensors in the trap detection zone can lead to density underestimation (Corlatti et al., 2020). Numerous methods have been proposed to account for such activity patterns in population size estimation (e.g. Nakashima et al., 2017), though a common approach is to focus the monitoring on periods when it is reasonable to suppose that all animals are active (Howe et al., 2017).

Given these potential sources of bias, the empirical validation of camera trap-based methods for estimating population size is crucial, yet remains limited. So far, most comparative studies have employed reference density values estimated using established methods such as conventional distance sampling (Rovero and Marshall, 2009; Caravaggi et al., 2016) or capture-recapture (Twining et al., 2022). However, these comparisons, carried out over a few years, often involve only a limited number of imprecise estimates from both camera-trap estimation methods and “reference” methods, often leading to conclusions that no difference can be detected between the two families of methods, which does not constitute proof of estimate equivalence. On the other hand, simulations offer a valuable alternative, enabling numerous replications of a specific camera trap study setup under fully known and user-controlled demographic and spatial conditions (Santini et al., 2022). However, so far, few studies have explored the conditions under which camera trap-based estimates may be biased or robust.

In this paper, we assess the feasibility of using camera traps to monitor roe deer (*Capreolus capreolus*) abundance over time through simulation. We examine how the number of camera traps, habitat characteristics, and animal behaviour affect the frequency of detections and the reliability of the resulting population estimates. Specifically, we explore how the differential use of habitats with varying detection properties, combined with the animals’ activity rhythms, affects the detection rates. To do so, we simulated realistic movements of animals, incorporating habitat selection and activity patterns (resting, foraging and exploring) informed by the GPS-monitoring in a roe deer population. We simulated a decreasing trend in population size across years, and a correlated increase in home-range size (Kjellander et al., 2004). We assessed the estimation precision and the ability to detect the simulated declining trend of two of the most widely classes of models for estimating population size: one based on individual encounters (random encounter model, Rowcliffe et al., 2008) and the other on associations (instantaneous sampling, Moeller et al., 2018). We also assessed the effect of placing all camera traps in the most selected habitat type, as well as the effect of stratified sampling, on the precision of the trend estimates – two commonly used procedures when setting up camera traps in the field. Finally, we explored the possibility to expand these results to more mobile species with larger home ranges (e.g., larger cervids or carnivores).

## Material and Methods

### Simulating realistic roe deer movements and their detection by camera traps

We first simulated a population of moving animals in the Chizé reserve, France (46.106^◦^ N, 0.424^◦^ W), during five years. We chose this enclosed 2614 ha forested study area because its roe deer population has been monitored for 50 years (Pellerin et al., 2017), and space use by the roe deer was extensively studied (Saïd et al., 2005; Pellerin et al., 2008, 2010; Gaudry et al., 2018). The population monitoring carried out in Chizé provides us with a wealth of data on demographic parameters (survival, reproductive rates, Gaillard et al., 1993, 2013). Using this well-characterized study area allowed us to calibrate our simulations with high biological and environmental realism, ensuring that movement patterns, detectability, and demographic trends are grounded in real-world data rather than generic assumption. This approach was initially designed to assess the feasibility of a practical monitoring program in this specific area, but the insights gained are applicable to similar forest ecosystems.

The northern part of the study area (1385 ha) is mainly composed of an oak stand (*Quercus* spp.) characterized by a high quality forage source, whereas the southern part (1229 ha) is mainly composed of a beech stand (*Fagus sylvatica*) with poor resource quality (Pettorelli et al., 2001). Consequently, roe deer density is higher in the northern part of the area, a spatial heterogeneity that we incorporated into our subsequent simulations.

We simulated roe deer movements within the study area using a framework that integrates three key behavioral components: sedentarity, velocity autocorrelation, and multi-state dynamics. While the Ornstein-Uhlenbeck process is a standard method for simulating attraction toward the home-range centre (Dunn and Gipson, 1977), this basic model is limited to a single attraction point and results in a simple circular or elliptic home-range geometry. To represent more realistic home ranges with multiple core areas, we simulated animals switching between several processes of the Ornstein-Uhlenbeck family (Blackwell, 1997). Specifically, we employed the Ornstein-Uhlenbeck with foraging (OUf) model (Fleming et al., 2014), which also accounts for velocity autocorrelation – where the animal’s direction at time *t* depends on its state at *t* − 1. By varying parameters such as attraction point locations, speed autocorrelation, and relocation autocorrelation in these processes, we represented diverse behaviours (e.g., foraging and inter-patch movements). Such multi-state models are commonly used to simulate animal trajectories (e.g., Breed et al., 2017; Michelot et al., 2019; Santini et al., 2022). Finally, periods of immobility were included into the trajectories to represent resting behavior.

Sets of attraction points for these OUf processes were simulated using the following approach: we first used the kernel method to estimate 25 utilization distributions using GPS data from 15 roe deer monitored in Chizé in March from 2003 to 2008 (some roe deer were monitored during multiple years; see Pellerin et al., 2008, for more details on the monitoring of these animals). We then identified the local modes of these distributions. This resulted in 25 sets of points (between 1 and 17 points per set), which we used to simulate the presence of attraction points for our simulated animals. To simulate the movement of an animal, we randomly selected one of these sets and randomly placed it on our study area, giving its centroid a probability equal to 0.64 to be located in the northern part to reflect the higher habitat quality in this region, as reported by Pettorelli et al. (2001). We made sure that all the simulated movements remained within the study area, to guarantee animal presence throughout the study and avoid edge effects on our results. When an animal survived across multiple years (see below), we used the same centroid for this animal during all years, thereby simulating its sedentarity.

We then simulated our movement model using this located set of attraction points. Analysis of the GPS data showed that we could roughly describe animal movements as randomly switching between three behaviours: resting (immobility), patch-level movements (movement concentrated within a very small portion of the home range, generally corresponding to foraging) and between-patch movements (larger movements within the home range). Our movement model therefore simulated three behaviour types: (i) resting (the animal does not move), (ii) Patch-level movements characterized by OUf processes with small variances and centred on one attraction point (see below); (iii) Between-patch movements characterized by OUf processes with larger variances, also centred on one attraction point. Appendix S1 gives more details on the precise parameterization of these OUf processes. We used the GPS data to derive rough estimates of the daily frequency and timing of these movements types (unpublished results), which were used to parameterize the movements in our simulations.

Fig. 1 gives a schematic description of the approach used to simulate the random switching between behaviours. We simulated the same activity cycle for all animals and all days of the study period. We simulated the animal movement during each day between the 1st and 31st of March, as follows. For each day (starting at 17:00 and ending at 16:59 next day), we first simulated the presence of a main resting period with a probability equal to 0.85, its starting time *m* being randomly drawn between 8:00 and 15:00. The duration (in hours) of this resting period was equal to 15-*m* + *e*, with *e* a residual randomly drawn from an Gaussian distribution N (0, 1), and truncated so that this resting period never ends after 16:59. We then randomly placed patch-level movements during the night between 17:00 and 08:00. We simulated the start time of these patch-level movements by simulating a Poisson process with constant intensity ensuring the presence of 0.66 patches per night on average, with a duration randomly drawn between 0h and 5h. Finally, we placed resting patches in the remaining periods (except between 6:00 and 8:00, when all deer are considered active). We simulated the start time of these resting patches by simulating a Poisson process with constant intensity ensuring the presence of 0.33 patches per night on average, with a duration also randomly drawn between 0h and 5h. Between-patch movements were simulated in the remaining periods.

**Figure 1.**
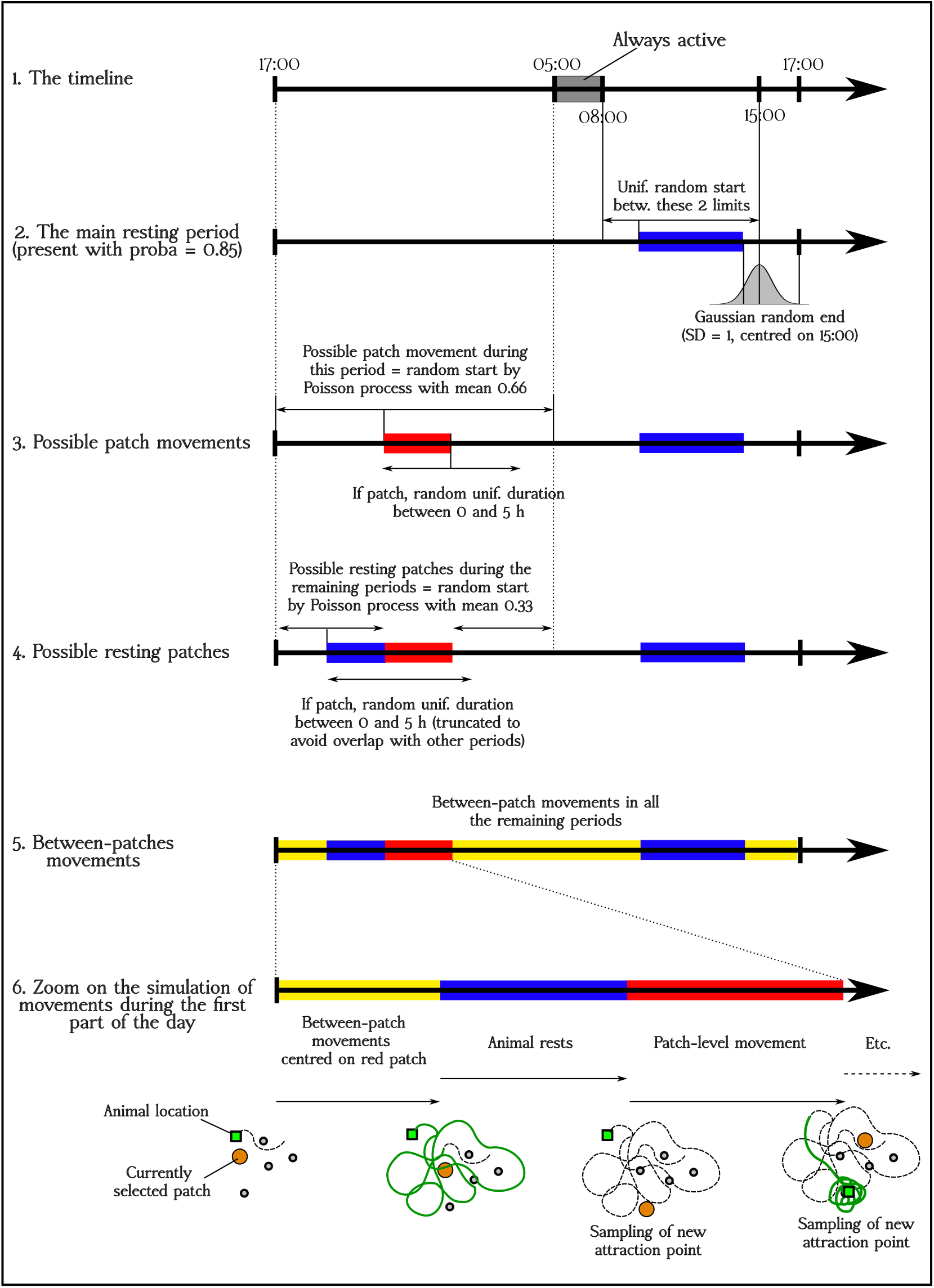
Simulation of the movement of a roe deer during a typical day, starting at 17:00 and ending at 16:59 the following day. The sequence 1 – 5 describes the simulation of the activity cycle. The point 6 illustrates the simulation of the movement process itself, zooming on the beginning of the day.

When the animal switched from one movement type to another (e.g., from between-patches to patch-level, from patch-level to between-patch, or from resting to between-patch), a new attraction point was randomly drawn from the simulated set. When the animal initiated a resting period, it stopped moving immediately and remained stationary until the period ended. An example of simulated movements is given in Fig. 2A.

**Figure 2.**
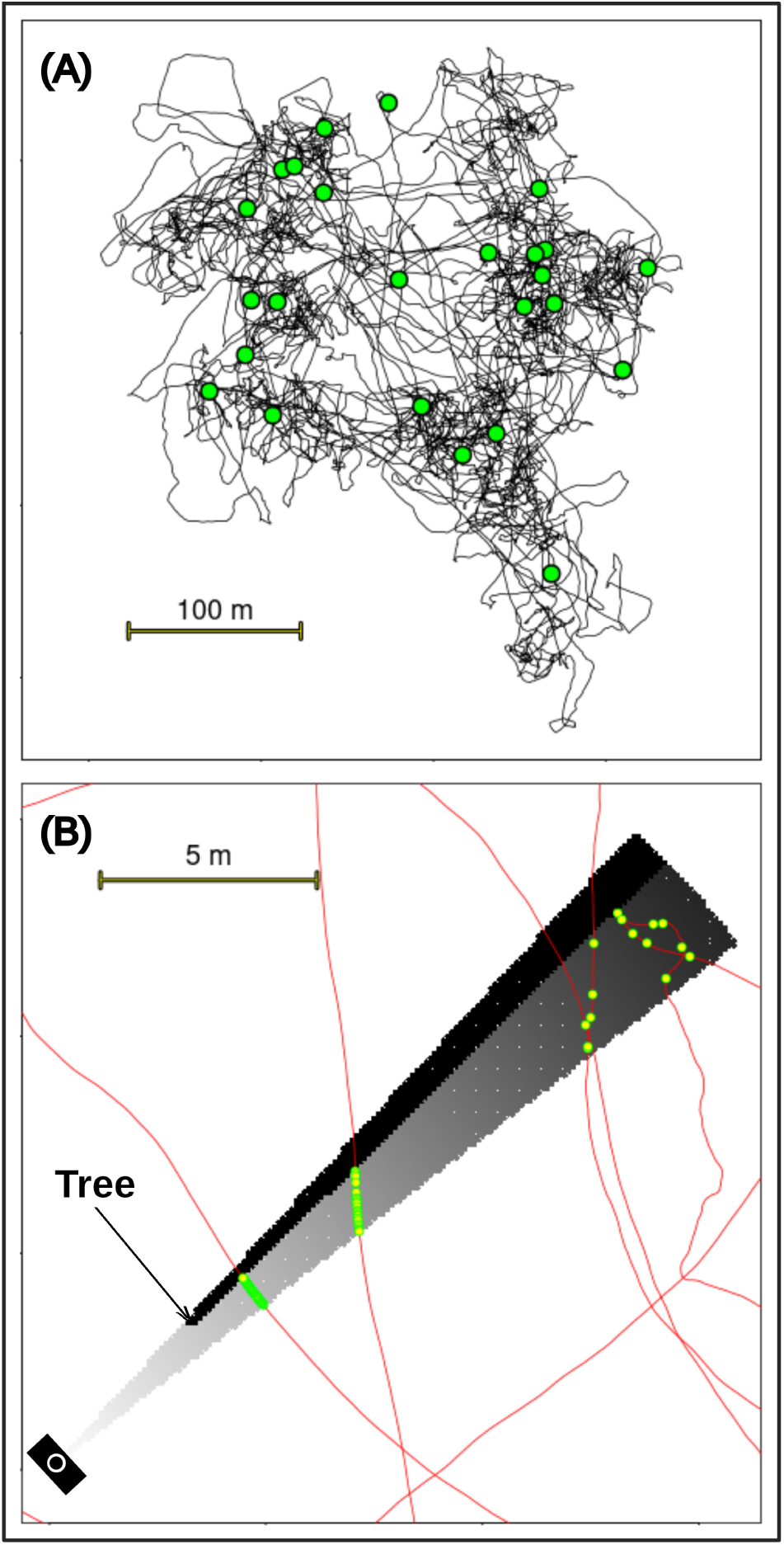
Simulation process used : (A) simulation of the movements of a roe deer using a model combining multiple Ornstein-Uhlenbeck process, including resting patches (green points) within the movement (when the animals stay at the same place without moving at all, (B) simulation of the detection process by a camera trap. The detection zone is modelled as a circular sector with a radius of 20 m and an angle equal to 10 degrees. Detection probability decreases with increasing distance from the trap, represented by varying shades of grey (darker = lower detection probability, and black = no detection at all). We also simulate the presence of trees that may limit the visibility within the detection zone. The movements of the animal simulated on panel (A) are displayed in red on this figure. The camera traps can capture the animals every second. The points correspond to the detections of the animals (more numerous close to the trap).

To satisfy the assumptions underlying the estimation methods, we randomly placed camera traps on the study area (see next sections), and simulated for each trap a motion sensor (i.e., camera traps triggered by the presence of animals, taking at most one picture per second). We defined a detection zone for each trap consisting of a 20 metres circular sector with an angle of 10 degrees. We chose this angle based on Rowcliffe et al. (2008), but the results of the simulations were not affected by this choice (see section S5.5 of Appendix S1 for a set of simulations with an angle of 42 degrees). When the movement of animals crossed the detection zone, we simulated imperfect detection by the sensors (Fig. 2B). We simulated a detection probability decreasing with the distance to the camera trap, and depending on the habitat type where the trap was located. Three habitat types are present in our area: (i) Coppice (76% of the study area), (ii) Open habitat (6% of the study area), (iii) Regeneration (18% of the study area). In Coppice and Open habitat types, we chose the parameters of the detection probability to ensure that the detection probability was equal to 0.99 at 5 metres from the trap and to 0.01 at 18 metres from the trap (calibrated visually from the results of Howe et al., 2017). We simulated the presence of trees obscuring the view in the Coppice habitat type (see Appendix S1 for more details on the distribution of tree diameters). In the Regeneration habitat type, we did not simulate explicitly the presence of such obstacles, but we subjectively chose parameters ensuring a detection probability of 0.99 at 0.5 metres and 0.01 at 3 metres. Finally, we supposed that an animal resting in the detection zone could not be detected by the sensors. More details on the parameters of the detection process are given in Appendix S1.

### Three types of simulations

#### Roe deer population, no habitat selection

We simulated a simple population dynamics process. We used the data collected on the Chizé study area for the last 45 years as a broad reference for key demographic parameters of a roe deer population (Gaillard et al., 1992, 1993, 1997, and subsequent updates fitted every year to field data). While not directly adopting published estimates, we fine-tuned these parameters to ensure a population decrease over time. Thus, we established an initial population size of 300 roe deer with a balanced sex ratio. Each year, 90% of females were assumed to reproduce, with an average of 1.7 fawns per reproducing female. To facilitate this population decline, we simulated a small survival probability for both the fawns (set at 0.3) and adults (set at 0.7). Though these probabilities were smaller than the values reported by Gaillard et al. (1993) in this area, such probabilities are not unrealistic and could be supposed to result from a high hunting pressure. This indeed resulted in a gradual population decline from 300 animals in Year 1 to 239 animals in Year 5 (see Appendix S1 for exact numbers of animals simulated every year).

We simulated an influence of the decreasing population density on animals’ home-range by gradually increasing mean home-range size over time. Kjellander et al. (2004) indeed observed this negative correlation between home-range size and population density in two populations. In reality this relationship can be complex (see discussion), but our aim here was to assess the effect of the opposite effects of the density decline and increasing home-range size on the number of detections by the traps. More precisely, we gradually increased the variance of the OUf process controlling for the between-patch movements. We also varied the list of sets of attraction points used to define the patches, considering only the 19 “small” sets with less than 10 attraction points during the first year, and progressively adding an increasingly larger sample of the 6 “large” sets with more than 10 attraction points in year 2 to 5. This led to a progressive increase of the home-range size, from an average of 13.1 ha during year 1 (SD = 4.4 ha) to 25.8 ha during year 5 (SD = 12.9 ha). Note that this increase in home-range size also affected the average movement speed of the animals, which may be important given that REM estimators require the knowledge of average speed.

We simulated the movements of all animals of the population during the month of March for every year of the 5-year period. This month was chosen because it coincides with common roe deer monitoring practices and a period of relative stability: it occurs after reproduction (when animals are no longer territorial) and the hunting season, yet before budburst and leaf unfolding, thereby maximizing detection probability. To simulate camera trap monitoring, we randomly placed either 100 or 25 camera traps over the study area, corresponding to a density of approximately 4 traps/km^2^ (i.e. ≈ one trap per roe-deer home range in average) and 1 trap/km^2^ respectively. We then used the REM and IS to estimate the population size from the data collected by these traps (see below). For each density of camera traps (4 or 1/km^2^), we carried out 1000 simulations of a camera trap monitoring.

#### Roe deer population, selection of paths neighborhood

The previous simulations did not account for habitat selection by the target species. In practice, researchers often use their knowledge of habitat use by the species to enhance their monitoring design. This can involve stratification, where pre-specified proportions of camera traps are placed in the different habitats, with overall density calculated as a weighted average of the habitat densities (Rowcliffe et al., 2008). Alternatively, some studies abandon their aim to obtain unbiased population size estimates to improve the precision of unbiased trends estimates. To achieve this, they maximize detection numbers by deploying camera traps only in highly selected habitats or in locations with high detection probabilities (e.g., Open habitat), assuming a constant bias in size estimates. It is a widely held assumption among practicioners that spatial randomness can be sacrificed to maximize encounter rates, as long as the primary goal is trend detection rather than absolute density estimation (pers. obs.). We assessed these two strategies with a new set of simulations, by simulating a strong habitat selection by the roe deer. We limited these simulations to a trap density of 4 traps/km^2^ (100 traps).

We simulated a strong selection of the human paths and roads by the roe deer in our study area. This specific selection of linear features by the roe deer is not expected in reality from a biological point of view, as such features are generally found at the home-range periphery (Seigle-Ferrand et al., 2021). However, this artificial scenario allows us to illustrate the ideal situation where the camera traps are placed in habitats with both high target species abundance and maximum detection probability. In addition, this approach allows us to generalize the results to other species using intensively linear structures, such as large predators (Dickie et al., 2019).

We defined “paths neighborhood” as the 20-metres buffer around each human path or road present in our study area. We simulated a selection of this habitat type by the roe deer: we first selected the location of the home-range centroid in the northern part with a probability of 0.64, as before. However, for this set of simulations, we ensured that: (i) the centroid was located in the paths neighborhood, (ii) it was located at more than 600 m from the border of the study area (to make sure that all the movements were located within the area). Between 3 and 10 attraction points were randomly selected at distances ranging from 100m to 300m from this centroid, making sure that all these points were located within the path neighborhood. We then carried out again the simulations described in the previous section, using these randomly generated attraction points instead of a randomly selected set of attraction points as before. As for the previous set of simulations, we varied the parameter *σ*^2^ of Ornstein-Uhlenbeck processes to simulate an increase in home-range size across years (see Appendix S1).

We then simulated two types of camera trap monitoring, corresponding to the two strategies described previously. In the first simulation type, we assessed the impact of placing all camera traps within highly selected habitat, specifically by randomly distributing them in the paths neighborhood.

In the second type of simulations, we evaluated the impact of stratified sampling on the precision of population size estimates. We simulated various stratified sampling designs, allocating 5% to 95% of the 100 traps to path neighborhoods in 5% increments. In practice, we did not simulate a single study with X% of traps in the path neighborhoods. Instead, we simulated two separate studies: one with all 100 traps placed in path neighborhoods and another with all 100 traps placed outside of path neighborhoods. To simulate stratified sampling, we combined the detection data from these two scenarios by randomly selecting the detections from X% of the traps from the first study (all traps in path neighborhoods) and the detections from 100-X% of the traps from the second study (all traps outside path neighborhoods). This allowed us to assess the impact of different proportions of traps in path neighborhoods on population size estimates without directly simulating a stratified sampling design within the population itself (which was less computer intensive). Although the animals simulated in the two scenarios (100% and 0% in paths neighborhood) are not the same, we showed in Appendix S1 that the results obtained with this approach do not differ from an approach where we simulated an actual stratified monitoring of the population where all 100 traps study the same animals. Note that we assessed stratification for the estimation of population size during one year only (i.e., no assessment for population trend estimation).

We simulated each strategy type 500 times.

#### Large-range roe deer population, no habitat selection

Finally, we aimed to assess the effect of the species home-range size on the results. To do so, we considered an animal with a home-range size similar to that of the red deer (*Cervus elaphus*). The simulated animal remains the same as before (e.g. same activity rhythm), with the only difference being a significantly larger home range, so that we will refer to this animal as the “large-range roe deer” in the following. We used the same movement algorithm as for the roe deer, with the following differences in parameterization: (i) we increased strongly the value of the variance of the between-patch and patch-level movement, (ii) the attraction points are randomly drawn in a rectangular box of 4 kilometres wide centred on the home range centroid. The simulated home-ranges covered 469 ha on average (interquartile range: 383 to 548 ha). We did not simulate a home-range size varying with population decline over the 5 years. Our primary objective here was to assess the effect of movement scale (i.e., large-range species) rather than the impact of temporal shifts in space use, which was addressed in our earlier scenarios. Nevertheless, additional simulations where the average home-range size increased from 405 ha in Year 1 to 724 ha in Year 5 led to the same results as those obtained using a constant home-range size (see Appendix S1, section S5.4.3). Consequently, we focus hereafter only on the constant home-range scenarios.

As in the previous scenario, we simulated a population size of 300 large-range roe deer. This corresponds to a high but not unrealistic density when compared to species with similar home-range sizes, such as the red deer (Borowik and Jwdrzejewska, 2018). We used published red deer estimates as a broad reference to derive realistic demographic parameters for this large-range roe deer (Bonenfant et al., 2002; Borowik and Jwdrzejewska, 2018). These parameters were fine-tuned ensure a declining population while allowing for comparison with the standard roe deer results. Specifically, we assumed a sex-ratio of 0.5, an adult annual survival of 0.76, and a reproduction equal to 0.5 fawns per female. The simulated number of animals was equal to 240 during the last year, corresponding to a 20% decrease in 5 years.

We then placed randomly either 4 traps/km^2^ or 1 trap/km^2^ over the study area. For each number of traps, we simulated 1000 times the camera trap monitoring.

### Population size estimates

For each simulation of each simulation set, we estimated the population size using two methods, REM and IS. To use the REM population size estimator, we need to consider the total number *Q* of animal-trap encounters. The estimator is:

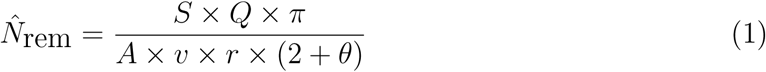

With *N̂* the population size estimate, *S* the study surface area, *A* the the cumulated activation duration of the traps, *v* the mean travel speed of the animals, and *r, θ* the depth and angle of the detection zone. Note that one of the biggest difficulties with the REM is that the mean travel speed of the animals is to be estimated, which can be an important source of imprecision of the population size estimation. In our simulations, we have considered that this mean speed is known. Although this will never be the case in practice, our aim was not to devise means to estimate travel speed.

To use the IS population size estimator, we needed to reshape our data. We discretized our study period (the month of March of every year during five years) into one-second intervals. We define “capture occasions” as *J* discrete moments marking the start of these intervals. For each capture occasion and each animal-camera trap pair, we determined whether a detection occurred (i.e., if the simulated motion sensor of the camera had detected the animal at that time). For each trap *i* and each capture occasion *j*, we define a new variable *n_ij_* corresponding to the number of animal-trap associations (number of animals present in the detection zone at that time and detected by the trap). Let *s_i_* be the surface area of the detection zone of the camera trap *i*. The IS population size estimator is:

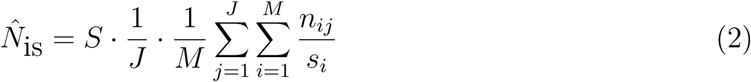

where *M* is the number of traps placed in the area.

The two estimation methods were applied each with three modalities: (i) we used the above estimators without accounting for the limited detectability of animals due to imperfect sensors, lack of visibility or activity rhythm (hereafter, these estimators are simply called REM and IS), (ii) we accounted for the limited visibility in the detection zones by modifying these estimators (hereafter called REM_d and IS_d, see below), (ii) we accounted for both this limited visibility and the reduced detectability during resting periods (hereafter called REM_da and IS_da). For the REM estimator, the mean travel speed was computed exactly from the simulated movements, using all data for the REM and REM_d, and using only data collected during active periods for the REM_da.

To account for the imperfect visibility in the detection zone in the REM estimator, we replaced *Q* in equation 1 by *Q/p̅*, where *p̅* is the proportion of all encounters that were detected by the traps. This average detection probability of encounters was estimated from the simulated data, where we knew the exact times when animals crossed trap detection zones. To account for this imperfect detectability of animals in the detection zone in the IS estimator, we replaced *s_i_* by *s_i_* × *d̅* in equation 2 where *d̅* is the mean detection probability of associations. This average detection probability of associations was also estimated from simulated data, where we knew the exact detection probability for all traps (as in fig. 2(B)). Finally, to account for the limited detectability of animals in the detection zone during resting, we changed our study period to remove all the data collected between 8:00 and 17:00, when the main resting period occurs.

For each estimator and each estimation modality, we estimated the population size for each year of the 5 years period, as well as the change rate: CR = (*N*_5_ − *N*_1_)*/N*_1_. We also calculated a mean estimated trend over the 5-year period:

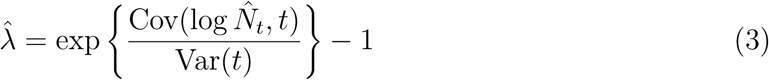

where *t* is the year and *N̂_t_* the estimated population size during year *t*. This parameter *λ̂* estimates the proportion of the population disappearing in one year. We refer to Gerrodette (1987) and Harris (1986) for the theoretical framework underlying this approach; the corresponding equations used in our simulations are detailed in Appendix S1, section S4.2.2).

When the simulations implied a stratified sample, we calculated one population size estimate for each strata and summed the two estimates. We used the bootstrap to estimate the variances and 90% confidence intervals on population size, change rate and trend estimates, by resampling for each estimation 1000 times with replacement the camera traps and recomputing the estimates to obtain a distribution of 1000 bootstrap estimates. When the simulations implied stratified samples, we bootstrapped the two strata separately.

In total, all these simulations required ≈ 25 days of calculation on a Dell T5610 workstation with a processor Intel(R) Xeon(R) CPU E54-2650 (2.6GHz). We provide a companion R package named `simCTChize`, available at https://github.com/ClementCalenge/simCTChize, containing all the code and data used for these simulations (also available on Zenodo: https://doi.org/10.5281/zenodo.16989360). Appendix S1 corresponds to the package PDF vignette, and describes how the user can install this package and easily reproduce the calculations carried out in this paper. This vignette also includes numerous additional details on the simulations that we omit in this paper to keep it as concise as possible (specific values of the parameters used for the movement processes or the detection functions of the camera traps, density of trees simulated in the detection zone, etc.). Finally, this package also contains a tutorial allowing the user to design their own simulations if they want to design a monitoring for another species.

## Results

### Simulating roe deer without habitat selection

We present the results of the simulations in Tables 1 (4 traps/km^2^) and 2 (1 trap/km^2^). In both cases, it is clear that the two estimators strongly underestimate the population size when the imperfect visibility in the detection zone is not accounted for. The IS estimator showed a far greater bias, underestimating by approximately 80% versus about 35% for the REM. Accounting for this imperfect visibility leads to an unbiased estimate of the population size with the REM, but IS still results in a biased estimate (≈ 20%). Accounting for the limited detectability of the animals in the detection zone when they rest results in an unbiased estimate for both IS and REM. The bootstrap allowed us to correctly estimate the standard error of the sampling distribution for all estimates. Across the 1000 simulations, the trends and change rate were, on average, correctly estimated for all modalities and methods, although considerable variability existed between the simulations.

With 4 traps/km^2^ on the study area, the population size estimation was rather imprecise (with a coefficient of variation of about 15% to 20%, depending on the method). A population size decrease was correctly identified in more than 80% (up to 91%) of the simulations with all methods; however, this decrease was significant only in 1/4 to 1/3 of the simulations, depending on the estimation method.

With 1 trap/km^2^ on the study area, the population size estimation was even more imprecise (with a coefficient of variation of about 30% to 40%). A population size decrease was estimated correctly in more than 2/3 (up to 74%) of the cases, but this decrease was significant in only 17% of the simulations).

### Simulating roe deer with habitat selection

When we simulated habitat selection and placed all the camera traps in the most selected habitat type, the population decrease was overestimated (Table 3). Indeed, the estimated population decline was approximately 8% per year, leading to a 28% decrease in year 5 (i.e., 1 − (1 − 0.08)^4^ = 0.28), instead of the simulated 20% decline (corresponding to a yearly decline of 5.5%).

Using a stratified sample allowed an unbiased population size estimation. The highest precision of the estimation was obtained when half of the traps were placed in the paths neighborhood and half out of this habitat (Fig. 3). However, this stratified sampling did not improve noticeably over a simple random sampling (the smallest coefficients of variation on this graph were similar to those obtained in Table 1).

**Figure 3.**
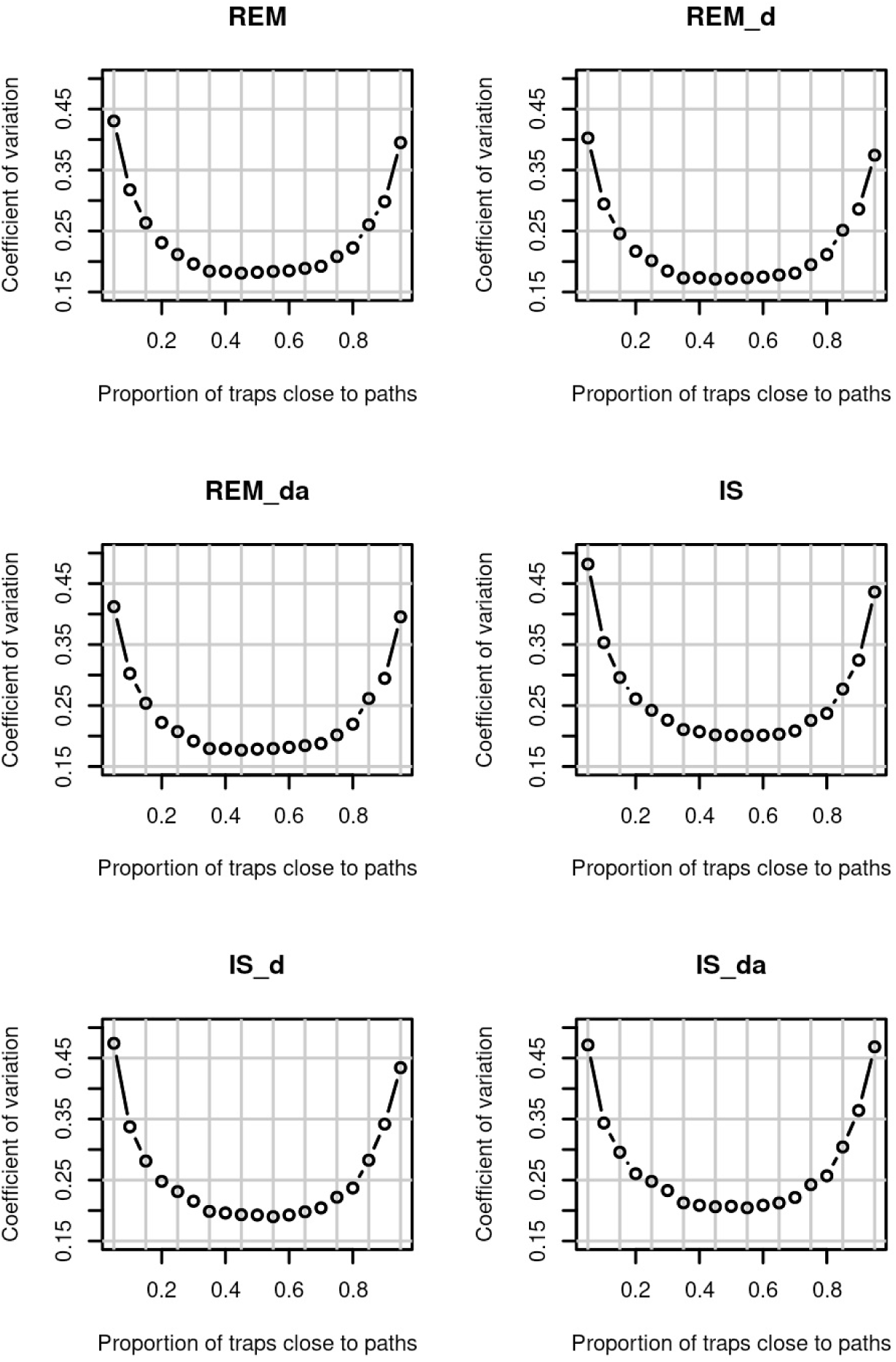
Precision of the estimation of population size in a monitoring of a roe deer population with 300 individuals with 4 camera traps/km^2^, using stratified sampling. The animals display a strong preference for the paths neighborhood on our study area (which represents 38% of the study area), and we vary the proportion of traps in this habitat type. We use two methods to estimate population size, the random encounter model (REM) and the instantaneous sampling (IS). For each method, we estimate the population size: (i) without accounting for imperfect visibility in the detection zones or the limited detectability of the animals in the detection zones during their resting periods (REM and IS), (ii) accounting for imperfect detectability of animals in the detection zones but not for these resting periods (REM_d, IS_d), (iii) accounting for both imperfect detectability and these resting periods (REM_da and IS_da)

**Table 1:** Results of 1000 simulations of a roe deer population monitored for 5 years using 4 camera traps/km^2^.

| Method | Param | TrueValue | MeanEst | SE | SEEst | SE.SEEst | Pcov | PDec | PsigD |
| --- | --- | --- | --- | --- | --- | --- | --- | --- | --- |
| REM | N1 | 300 | 237.31 | 37.875 | 38.687 | 6.215 | 0.511 |  |  |
| REM | N5 | 239 | 181.767 | 33.792 | 31.553 | 6.114 | 0.444 |  |  |
| REM | $\lambda$ | -0.0547 | -0.063 | 0.049 | 0.048 | 0.007 | 0.869 | 0.906 | 0.364 |
| REM | CR | -0.2033 | -0.22 | 0.165 | 0.165 | 0.043 | 0.859 | 0.906 | 0.378 |
| REM_d | N1 | 300 | 303.86 | 45.68 | 49.51 | 7.853 | 0.923 |  |  |
| REM_d | N5 | 239 | 232.533 | 40.045 | 40.352 | 7.572 | 0.882 |  |  |
| REM_d | $\lambda$ | -0.0547 | -0.063 | 0.049 | 0.048 | 0.007 | 0.867 | 0.906 | 0.367 |
| REM_d | CR | -0.2033 | -0.22 | 0.165 | 0.169 | 0.042 | 0.9 | 0.906 | 0.327 |
| REM_da | N1 | 300 | 303.224 | 47.154 | 50.959 | 8.427 | 0.928 |  |  |
| REM_da | N5 | 239 | 232.706 | 41.018 | 41.146 | 7.83 | 0.882 |  |  |
| REM_da | $\lambda$ | -0.0547 | -0.062 | 0.051 | 0.05 | 0.007 | 0.867 | 0.889 | 0.344 |
| REM_da | CR | -0.2033 | -0.217 | 0.172 | 0.176 | 0.045 | 0.904 | 0.891 | 0.311 |
| IS | N1 | 300 | 54.903 | 9.955 | 10.084 | 1.897 | 0 |  |  |
| IS | N5 | 239 | 42.983 | 8.975 | 8.3 | 1.844 | 0 |  |  |
| IS | $\lambda$ | -0.0547 | -0.058 | 0.055 | 0.055 | 0.008 | 0.88 | 0.843 | 0.289 |
| IS | CR | -0.2033 | -0.198 | 0.195 | 0.198 | 0.057 | 0.87 | 0.848 | 0.279 |
| IS_d | N1 | 300 | 262.512 | 47.812 | 46.688 | 11.689 | 0.737 |  |  |
| IS_d | N5 | 239 | 205.374 | 40.169 | 37.611 | 9.195 | 0.682 |  |  |
| IS_d | $\lambda$ | -0.0547 | -0.057 | 0.056 | 0.054 | 0.008 | 0.884 | 0.854 | 0.279 |
| IS_d | CR | -0.2033 | -0.197 | 0.196 | 0.197 | 0.056 | 0.882 | 0.841 | 0.276 |
| IS_da | N1 | 300 | 298.869 | 57.784 | 56.794 | 15.407 | 0.881 |  |  |
| IS_da | N5 | 239 | 233.97 | 49.233 | 45.015 | 12.228 | 0.849 |  |  |
| IS_da | $\lambda$ | -0.0547 | -0.057 | 0.059 | 0.057 | 0.009 | 0.885 | 0.833 | 0.272 |
| IS_da | CR | -0.2033 | -0.193 | 0.211 | 0.214 | 0.067 | 0.891 | 0.834 | 0.257 |

Abbreviation: REM, random encounter model; IS, instantaneous sampling; \_d, accounting for imperfect visibility; \_da, accounting for imperfect visibility and resting periods; N1 and N5, population size during years 1 and 5 respectively; $\lambda$ , trend over five years; CR = $[N5-N1]/N1$ ; TrueValue, simulated value of a parameter; MeanEst, mean of estimated values; SE, standard deviation of estimated values; SEEst, mean of the bootstrap estimates of SE; SE.SEEst, standard deviation of the bootstrap estimates of SE; Pcov, estimated coverage probability of the 90% confidence interval; Pdec, proportion of simulations for which estimated $\lambda$ or CR is negative; PsigD, proportion of simulations for which this decrease is significant at the 10% level.

**Table 2:** Results of 1000 simulations of a roe deer population monitored for 5 years using 1 camera trap/km^2^.

| Method | Param | TrueValue | MeanEst | SE | SEEst | SE.SEEst | Pcov | PDec | PsigD |
| --- | --- | --- | --- | --- | --- | --- | --- | --- | --- |
| REM | N1 | 300 | 235.668 | 78.199 | 72.339 | 23.699 | 0.688 |  |  |
| REM | N5 | 239 | 181.565 | 64.232 | 59.381 | 21.176 | 0.638 |  |  |
| REM | $\lambda$ | -0.0547 | -0.057 | 0.099 | 0.097 | 0.026 | 0.87 | 0.732 | 0.186 |
| REM | CR | -0.2033 | -0.161 | 0.383 | 0.417 | 0.358 | 0.871 | 0.738 | 0.193 |
| REM_d | N1 | 300 | 301.727 | 93.669 | 92.96 | 29.521 | 0.868 |  |  |
| REM_d | N5 | 239 | 232.242 | 76.207 | 76.196 | 26.018 | 0.864 |  |  |
| REM_d | $\lambda$ | -0.0547 | -0.057 | 0.099 | 0.097 | 0.026 | 0.872 | 0.732 | 0.192 |
| REM_d | CR | -0.2033 | -0.161 | 0.383 | 0.417 | 0.289 | 0.887 | 0.738 | 0.161 |
| REM_da | N1 | 300 | 301.069 | 97.229 | 95.555 | 31.466 | 0.871 |  |  |
| REM_da | N5 | 239 | 231.893 | 78.276 | 77.683 | 26.871 | 0.861 |  |  |
| REM_da | $\lambda$ | -0.0547 | -0.056 | 0.104 | 0.1 | 0.027 | 0.861 | 0.722 | 0.187 |
| REM_da | CR | -0.2033 | -0.15 | 0.413 | 0.436 | 0.276 | 0.877 | 0.734 | 0.159 |
| IS | N1 | 300 | 54.785 | 20.805 | 18.818 | 7.119 | 0 |  |  |
| IS | N5 | 239 | 42.953 | 16.803 | 15.415 | 6.117 | 0 |  |  |
| IS | $\lambda$ | -0.0547 | -0.053 | 0.115 | 0.111 | 0.034 | 0.856 | 0.701 | 0.159 |
| IS | CR | -0.2033 | -0.107 | 0.503 | 0.628 | 0.908 | 0.848 | 0.701 | 0.166 |
| IS_d | N1 | 300 | 263.562 | 97.551 | 87.342 | 36.81 | 0.768 |  |  |
| IS_d | N5 | 239 | 204.538 | 75.535 | 69.373 | 28.575 | 0.745 |  |  |
| IS_d | $\lambda$ | -0.0547 | -0.053 | 0.113 | 0.108 | 0.03 | 0.851 | 0.699 | 0.157 |
| IS_d | CR | -0.2033 | -0.119 | 0.482 | 0.605 | 1.06 | 0.856 | 0.703 | 0.167 |
| IS_da | N1 | 300 | 301.313 | 120.668 | 105.91 | 50.082 | 0.816 |  |  |
| IS_da | N5 | 239 | 233.349 | 91.489 | 83.106 | 36.841 | 0.826 |  |  |
| IS_da | $\lambda$ | -0.0547 | -0.051 | 0.124 | 0.115 | 0.033 | 0.856 | 0.699 | 0.147 |
| IS_da | CR | -0.2033 | -0.091 | 0.586 | 0.813 | 3.492 | 0.85 | 0.688 | 0.155 |

Note: We use the same abbreviations as in Table 1

### Simulating a large-range roe deer population

The simulations of a large-range roe deer population led to the same conclusions regarding the bias and precision of population size estimates: accounting for imperfect visibility in the detection zone led to less biased estimates by the REM and the IS, and accounting for the reduced detectability of resting animals by sensors led to unbiased estimates for the IS. The precision of population size estimates was comparable to that obtained for standard roe deer, for both 4 traps/km^2^ (Table 4) and 1 trap/km^2^ (Table 5). However, the trend and change rate estimations were much more precise. With 4 traps/km^2^, the simulated decrease was nearly always identified, and was significant in more than 95% of the cases for the REM, and in more than 50% (up to 76%) of the cases for the IS. With 1 trap/km^2^, the decrease was also frequently identified in most cases with the REM, and in more than 80% of the cases for the IS. However, the decrease reached statistical significance in only a minority of cases – from 33% (IS) to 66% (REM) of the cases.

**Table 3:** Results of 500 simulations of a roe deer population monitored for 5 years using 4 camera traps/km^2^, where all animals display a preference for the neighborhood of paths and where all the traps have been placed in this neighborhood.

| Method | Param. | MeanEst | SE | SEEst | SE.SEEst | Pcov | PDec | PsigD |
| --- | --- | --- | --- | --- | --- | --- | --- | --- |
| REM | -0.0547 | -0.08 | 0.03 | 0.03 | 0.00 | 0.73 | 1.00 | 0.87 |
| REM_d | -0.0547 | -0.08 | 0.03 | 0.03 | 0.00 | 0.73 | 1.00 | 0.86 |
| REM_da | -0.0547 | -0.08 | 0.03 | 0.03 | 0.00 | 0.75 | 1.00 | 0.83 |
| IS | -0.0547 | -0.08 | 0.04 | 0.04 | 0.01 | 0.77 | 0.99 | 0.70 |
| IS_d | -0.0547 | -0.08 | 0.04 | 0.04 | 0.01 | 0.79 | 0.99 | 0.71 |
| IS_da | -0.0547 | -0.08 | 0.04 | 0.04 | 0.01 | 0.81 | 0.99 | 0.69 |

Note: We use the same abbreviations as in Table 1

**Table 4:** Results of 500 simulations of a large-range roe deer population monitored for 5 years using 4 camera traps/km^2^.

| Method | Param | TrueValue | MeanEst | SE | SEEst | SE.SEEst | Pcov | PDec | PsigD |
| --- | --- | --- | --- | --- | --- | --- | --- | --- | --- |
| REM | N1 | 300 | 207.511 | 40.895 | 40.456 | 4.87 | 0.325 |  |  |
| REM | N5 | 240 | 165.729 | 33.028 | 32.409 | 4.006 | 0.315 |  |  |
| REM | $\lambda$ | -0.0543 | -0.055 | 0.013 | 0.012 | 0.002 | 0.875 | 1 | 0.989 |
| REM | CR | -0.2 | -0.2 | 0.047 | 0.046 | 0.009 | 0.885 | 1 | 0.976 |
| REM_d | N1 | 300 | 296.257 | 52.281 | 57.931 | 6.602 | 0.916 |  |  |
| REM_d | N5 | 240 | 236.537 | 41.961 | 46.383 | 5.297 | 0.916 |  |  |
| REM_d | $\lambda$ | -0.0543 | -0.055 | 0.013 | 0.012 | 0.002 | 0.874 | 1 | 0.991 |
| REM_d | CR | -0.2 | -0.2 | 0.047 | 0.046 | 0.009 | 0.937 | 1 | 0.993 |
| REM_da | N1 | 300 | 296.516 | 52.561 | 57.848 | 6.605 | 0.921 |  |  |
| REM_da | N5 | 240 | 236.556 | 42.166 | 46.411 | 5.377 | 0.916 |  |  |
| REM_da | $\lambda$ | -0.0543 | -0.055 | 0.015 | 0.015 | 0.003 | 0.879 | 1 | 0.968 |
| REM_da | CR | -0.2 | -0.2 | 0.057 | 0.054 | 0.011 | 0.931 | 0.999 | 0.964 |
| IS | N1 | 300 | 51.406 | 10.502 | 10.236 | 1.301 | 0 |  |  |
| IS | N5 | 240 | 41.187 | 8.642 | 8.3 | 1.119 | 0 |  |  |
| IS | $\lambda$ | -0.0543 | -0.054 | 0.022 | 0.022 | 0.005 | 0.888 | 0.988 | 0.761 |
| IS | CR | -0.2 | -0.195 | 0.086 | 0.084 | 0.02 | 0.872 | 0.982 | 0.704 |
| IS_d | N1 | 300 | 259.127 | 48.484 | 47.458 | 7.392 | 0.748 |  |  |
| IS_d | N5 | 240 | 208.346 | 39.598 | 39.027 | 6.823 | 0.76 |  |  |
| IS_d | $\lambda$ | -0.0543 | -0.053 | 0.026 | 0.026 | 0.007 | 0.892 | 0.974 | 0.672 |
| IS_d | CR | -0.2 | -0.19 | 0.101 | 0.099 | 0.034 | 0.889 | 0.959 | 0.611 |
| IS_da | N1 | 300 | 295.255 | 57.31 | 55.822 | 10.413 | 0.879 |  |  |
| IS_da | N5 | 240 | 237.832 | 47.503 | 46.435 | 10.191 | 0.874 |  |  |
| IS_da | $\lambda$ | -0.0543 | -0.053 | 0.031 | 0.03 | 0.008 | 0.893 | 0.957 | 0.583 |
| IS_da | CR | -0.2 | -0.187 | 0.12 | 0.117 | 0.045 | 0.897 | 0.936 | 0.51 |

Note: We use the same abbreviations as in Table 1

**Table 5:** Results of 500 simulations of a large-range roe deer population monitored for 5 years using 1 camera traps/km^2^.

| Method | Param | TrueValue | MeanEst | SE | SEEst | SE.SEEst | Pcov | PDec | PsigD |
| --- | --- | --- | --- | --- | --- | --- | --- | --- | --- |
| REM | N1 | 300 | 210.484 | 80.766 | 78.826 | 19.945 | 0.655 |  |  |
| REM | N5 | 240 | 167.931 | 65.534 | 62.97 | 15.982 | 0.658 |  |  |
| REM | $\lambda$ | -0.0543 | -0.055 | 0.028 | 0.025 | 0.009 | 0.835 | 0.979 | 0.679 |
| REM | CR | -0.2 | -0.197 | 0.103 | 0.105 | 0.058 | 0.846 | 0.967 | 0.63 |
| REM_d | N1 | 300 | 299.684 | 104.803 | 113.481 | 27.342 | 0.902 |  |  |
| REM_d | N5 | 240 | 239.059 | 84.642 | 90.663 | 21.937 | 0.895 |  |  |
| REM_d | $\lambda$ | -0.0543 | -0.055 | 0.028 | 0.025 | 0.01 | 0.835 | 0.979 | 0.673 |
| REM_d | CR | -0.2 | -0.197 | 0.103 | 0.105 | 0.059 | 0.88 | 0.967 | 0.645 |
| REM_da | N1 | 300 | 299.633 | 104.979 | 112.892 | 27.098 | 0.902 |  |  |
| REM_da | N5 | 240 | 238.704 | 84.816 | 90.233 | 22.085 | 0.903 |  |  |
| REM_da | $\lambda$ | -0.0543 | -0.055 | 0.034 | 0.028 | 0.01 | 0.817 | 0.95 | 0.582 |
| REM_da | CR | -0.2 | -0.194 | 0.134 | 0.123 | 0.069 | 0.88 | 0.936 | 0.561 |
| IS | N1 | 300 | 51.964 | 20.603 | 19.837 | 5.205 | 0 |  |  |
| IS | N5 | 240 | 41.302 | 16.859 | 15.898 | 4.35 | 0 |  |  |
| IS | $\lambda$ | -0.0543 | -0.056 | 0.05 | 0.043 | 0.017 | 0.828 | 0.878 | 0.423 |
| IS | CR | -0.2 | -0.185 | 0.199 | 0.227 | 0.348 | 0.816 | 0.85 | 0.376 |
| IS_d | N1 | 300 | 260.332 | 95.011 | 91.322 | 25.821 | 0.823 |  |  |
| IS_d | N5 | 240 | 208.851 | 80.911 | 74.454 | 22.685 | 0.808 |  |  |
| IS_d | $\lambda$ | -0.0543 | -0.055 | 0.055 | 0.047 | 0.018 | 0.844 | 0.858 | 0.365 |
| IS_d | CR | -0.2 | -0.178 | 0.224 | 0.249 | 0.382 | 0.841 | 0.834 | 0.34 |
| IS_da | N1 | 300 | 295.397 | 110.777 | 106.145 | 33.628 | 0.862 |  |  |
| IS_da | N5 | 240 | 238.746 | 98.29 | 87.876 | 31.69 | 0.854 |  |  |
| IS_da | $\lambda$ | -0.0543 | -0.054 | 0.065 | 0.054 | 0.02 | 0.824 | 0.819 | 0.328 |
| IS_da | CR | -0.2 | -0.164 | 0.277 | 0.325 | 0.726 | 0.813 | 0.791 | 0.289 |

Note: We use the same abbreviations as in Table 1

We also compared various descriptive statistics between the standard and large-range roe deer populations (Table 6). The mean number of encounters with large range roe deer was more than double that of the standard roe deer. However, the mean speed of large-range roe deer was also more than twice as high as that of the standard roe deer, so that the resulting number of animal-trap associations (and its standard deviation) was similar for the two species. As a result, the population size estimates were comparable between the two populations, whether estimated with REM or by IS. However, since the home-range size was 36 times larger for the large-range roe deer during year 1, individuals of this species visited 7 times more traps, on average, than standard roe deer. Consequently, each camera trap detected 7 times more large-range individuals on average. Therefore, the death of an individual (or the recruitment of a new one) between years affected the number of encounters and associations across a greater number of traps than for the standard species. Therefore, when using bootstrap to calculate the standard deviation of the trends, it was much easier to detect large-range roe deer population changes than standard roe deer population changes.

**Table 6:** Mean value (SD in parentheses) of various statistics calculated for each simulation of camera trap monitoring of roe deer and red deer population (300 individuals,100 camera traps).

|  | Standard Roe Deer | Large-Range Roe Deer |
| --- | --- | --- |
| Number of associations | 19172 (SD = 3476) | 17951 (SD = 3667) |
| Number of encounters | 684 (SD = 101) | 1584 (SD = 279) |
| Travel speed ($\text{m.s}^{-1}$) | 0.0165 (SD = 0.0001) | 0.0387 (SD = 0.0002) |
| Number of traps visited by an animal | 0.459 (SD = 0.056) | 3.492 (SD = 0.62) |
| Number of animals seen by a trap | 1.376 (SD = 0.167) | 10.475 (SD = 1.86) |

## Discussion

We have simulated different settings of camera trap monitoring of a roe deer population. Our simulations revealed the importance of accounting for imperfect visibility in the detection zone to estimate the population size correctly with the REM method and even more with the IS method. Moreover, the limited detectability of animals due to their activity rhythm needed to be considered, but exclusively for the IS method. This was less important for the REM, as long as the mean speed was correctly estimated by including inactivity periods. Although the population trend or the change rate were accurately estimated, even without accounting for imperfect visibility or activity rhythm (Table 1), the estimates were very imprecise and often not significantly different from zero. With 1 camera trap/km^2^, which is already at the upper end of the density of camera traps typically deployed for this species (e.g. Marcon et al., 2019; Gaudiano et al., 2021; Henrich et al., 2022; Palencia et al., 2022), the roe deer population size and trend estimates were so imprecise that they were virtually useless (Table 2). When we simulated animals with a much larger home range, the trend estimates were much more precise, even with only 1 trap/km^2^. We also showed that placing the traps preferentially in the habitat selected by the species led to a strong overestimate of the population decrease. Finally, we showed that using a stratified sampling did not improve the precision of population size estimation compared to random sampling.

### Generality of findings across estimation methods

We used the REM and IS to estimate the population size of the large-range and standard roe deer populations. We did not compare the two methods to determine which one was the more interesting for the monitoring. Such a comparison would be difficult, as the two methods do not require the same elements to be used. Thus, our results seem to indicate that the two methods give comparable results, though the REM seems more precise than the IS in our simulations. However, as Nakashima (2022) noted, the mean movement speed, which we supposed known in our simulations, is never known in practical studies and has to be estimated. The uncertainty of this parameter estimation results in a larger uncertainty of the population size estimates, so that comparing IS and REM is not fair without accounting for the speed estimation. There are numerous other methods available to estimate the population size based on camera trap data collected on unmarked populations (Nakashima et al., 2017; Moeller et al., 2018), and we chose IS and REM because they were the most easily automated (easily computed estimator that does not rely on the complex fit of a model) and because they belonged to the two main families of estimation methods: methods relying on the encounters and methods relying on associations. We think that our simulation results, that are similar for REM and IS, also generalize to other methods such as camera traps distance sampling, space-to-event or time-to-event models (Howe et al., 2017; Moeller et al., 2018; Calenge, 2026).

### Detectability: a prerequisite for unbiased population estimation

Our results are consistent with previous research highlighting the importance of accounting for the imperfect visibility in the detection zone to obtain unbiased population size estimates (Moeller et al., 2018, 2023). While we assumed perfect knowledge of detection probabilities in our simulations, real-world applications require estimating these probabilities, which can introduce additional uncertainty into population size estimates. Accounting for this imperfect detectability of animals in the detection zones with methods relying on detection is basically the rationale behind the camera-trap distance sampling (Howe et al., 2017). Thus, IS without accounting for detectability is very similar to camera-trap distance sampling where the detection probability is set equal to 1.

This stresses the necessity of estimating detectability when designing a camera-trap studies aimed at absolute population density estimation. Moeller et al. (2023) reviewed various approaches to integrate imperfect detectability into the estimation process. These range from distance sampling frameworks used to modele the detection probability of associations (e.g., Howe et al., 2017) or encounters (e.g., Rowcliffe et al., 2011), to the use of experimental calibration to directly estimate these probabilities (McIntyre et al., 2020). We refer the reader to this body of literature for specific implementation details.

### Robustness of estimates to temporal changes in home-range size

In our study, the roe deer population was characterized by a home-range size increasing with time, as a result of the decreasing animal density. This correlation can occur under various ecological conditions. For instance, in the hypothetical case where reduced habitat productivity (e.g., caused by shifts in land management practices) triggers a population decline due to diminished foraging resources, the habitat productivity hypothesis predicts that resource scarcity should also drive an increase in home-range sizes, as individuals compensate for limited resources (Harestad and Bunnel, 1979). Alternatively, this correlation might emerge from density-dependent behavioral mechanisms: declining populations could reduce competition among males, potentially allowing for larger individual territories. Such an inverse relationship between male home-range size and density has been identified in the roe deer by Vincent et al. (1995). In this study, we simulated a negative correlation between home-range size and population, but it is noteworthy that different conditions may lead to a positive correlation, e.g. under habitat productivity hypothesis. For example, if population reductions stem from non-habitat factors (e.g., hunting pressure), lower densities could increase resource availability per capita, thereby shrinking home ranges. Regardless of the underlying mechanism, and whatever their sign, such correlations do exist in nature, and may bias density estimation with the REM when the increase in home-range size is also correlated with increased mean animal travel speed: in our simulations, the mean speed was 30% smaller in year 1 than in year 5. As the population size estimator is inversely related to this mean speed, using a single value of mean speed for all years, as commonly done in studies using REM, would result in a substantially biased estimation (e.g., using the mean speed of year 1 for all years in equation 1 would result to a 30% underestimation in population size during year 5).

### Robustness of REM versus sensitivity of IS to resting periods

Accounting for activity patterns had a minimal impact on REM population size estimates. The random camera placement meant resting areas were sampled as frequently as active ones. While resting animals are undetectable by motion sensors, their entry and exit from a trap’s detection zone, which requires movement, allow encounter detection. Therefore, resting periods within encounters did not affect REM results. On the other hand, since association-based methods strongly rely on the detection of animals at any moment of the encounters, the presence of a resting period during an encounter may lead to a large number of associations missed by a motion-triggered sensor, which explains why association-based methods are more sensitive to activity patterns. Avoiding to include the monitoring periods during which a substantial fraction of the animals are resting allows to remove this small bias, which is also consistent with the findings of previous authors (Howe et al., 2017). This is the simplest approach to account for the activity rhythm of animals, and although other approaches are possible (e.g., by estimating the proportion of the day during which the animal is active and by modifying the duration of the study period in the equations, Nakashima et al., 2017), our approach led to unbiased population size estimates.

Our results raise the question of whether motion-sensors or time-lapse programming of camera traps are preferable for camera trap studies. Association-based methods require time-lapse data, which can be obtained by direct time-lapse programming or post-processing of motion-triggered trap data. Direct time-lapse programming eliminates sensor sensitivity bias, but image analysis, often AI-assisted (Rigoudy et al., 2023), may still introduce detectability issues related to distance and visibility. Nevertheless, contrary to motion sensors, direct time-lapse programming is not affected by the activity of the animal and detects both active and resting animals equally. However, the short time-lapses needed for association-based methods (Howe et al., 2017) raise storage challenges: trap’s memory fills up quickly and the batteries are rapidly depleted. Therefore, direct time-lapse programming might not be adequate with association-based methods.

### Biased population trends: when temporal shifts in behavior and detection go unmodeled

The fact that it was not necessary to account for the activity rhythm of animals and imperfect visibility in the detection zones in our trend estimate is probably a result of our simulation design. Indeed, as long as the simulated detection probability and activity rhythms of animals were the same across years, the resulting bias in the population size estimate was constant, allowing for unbiased estimation of the population trends. However, we do not expect this result to hold if the detectability in the detection zones or the activity rhythm changes across years. For example, the encroachment of shrubs into the study area would lead to an average detection probability decreasing with time. Considering constant detection, or not considering detection at all, would result in an increasing underestimation of population size with time and, consequently in our example of declining population, in an overestimation of the decreasing trend. Not accounting for the activity rhythm or the imperfect detection in trend estimation is only possible under the assumption that these phenomena do not vary with time.

This variation of animal behaviour across years does explain our overestimation of the decline when the traps are placed only in highly-selected habitats. Despite the intuitive aim of maximising detections by positioning all traps in the preferred path neighborhood, this placement led to an overestimation of the decreasing trend (simulated 20% decline vs. estimated 28% over 5 years). This bias stemmed from a simulated change in space use over time: as home-range size increased in later years, individuals spent more time travelling between selected patches, and consequently less time in paths neighborhood in year 5 than in year 1 (Fig. 4). This between-year variability of the mean proportion of the population located in the paths neighborhood at a given moment directly resulted in the overestimation of the population decrease. Appendix S1 demonstrates mathematically that combining the actual population decrease with this decrease in the proportion of the population in the paths neighborhood results in an annual decrease of 8% of the number of encounters/associations in this habitat. This finding has important consequences for camera trap monitoring, especially given that (i) placing camera traps in good habitats is a common practice to maximize the number of animals detections in population monitoring (e.g., Engeman, 2005; Bengsen et al., 2011); (ii) there are compelling reasons to expect space use to vary from one year to the next (e.g., Uboni et al., 2015). Thus the common practice of placing camera traps in highly selected habitats carries a high risk of bias in trend estimation.

**Figure 4.**
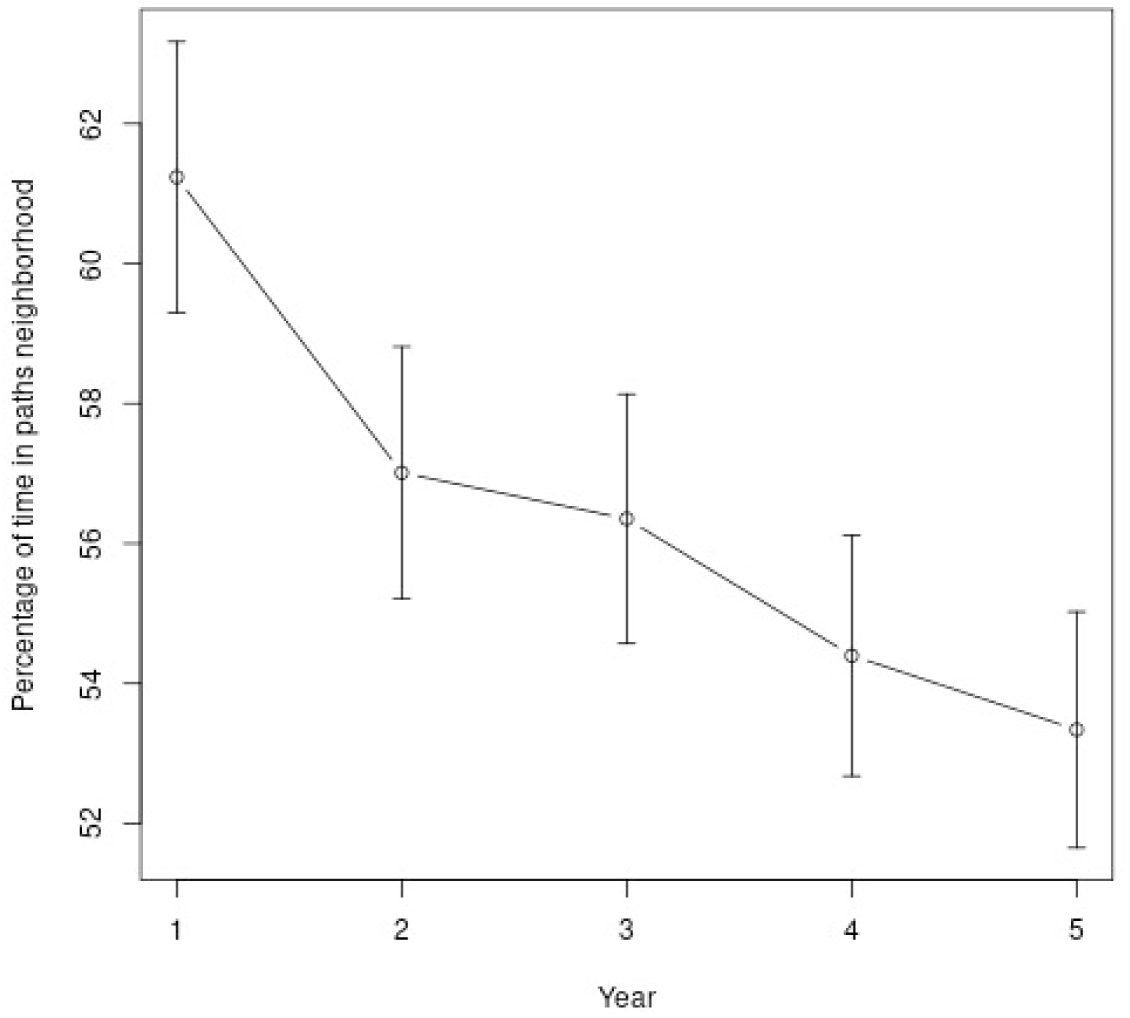
Estimated mean time spent by the roe deer in the paths neighborhood every year, in our simulations of a camera trap monitoring over 5 years. Since the home-range size of the animals increases with time, animals spend less time in average in “good” habitats.

### Evaluating alternative sampling strategies: stratification and grid designs

While altering the sampling strategy could potentially enhance estimate precision, we found no suitable approach; specifically, stratification did not improve precision compared to simple random sampling. Another common strategy involves placing camera traps at the nodes of a regular grid, thereby ensuring a good spatial coverage of the population. We did not consider this grid-based strategy, as it would have yielded results similar to random sampling: given that our simulation design generated a random distribution of home ranges across the study area, the spatial relationship between animals and traps would have been equally random for both grid-based and simple random sampling. The estimates would therefore have been equivalent. Furthermore, density estimation theory assumes random trap placement (e.g. Rowcliffe et al., 2008), making grid-based sampling potentially unsuitable.

### Limited value of roe deer monitoring even under idealized simulation scenarios

Several aspects of our simulations were oversimplistic: our simulated animals corresponded to a point moving in a 2D space, and not to a volume moving in a 3D space. This simplification may affect some of our results. For example, when the point representing the animal was located behind a tree, it was supposed to be completely hidden by the tree, whereas in reality animals can be detected by traps when they are located behind small trees. We think that this did not affect strongly our results, as we simulated the presence of trees only for the Coppice habitat type (which represents 76% of the study area), and always at a low density (see Appendix S1 for details): there was in average 0.37 trees simulated in the detection zone of a camera trap located in the Coppice habitat. Simulations showed that the trees obscured in average only 1.2% of the detection zone of a trap in the Coppice habitat, and these trees had more chance to be located far from the traps, i.e., where the detection probability is already low, since this is where the area of the detection zone is the largest.

More generally, simplifying assumptions in our simulations led to an overestimation of precision in our results: our camera traps did not deteriorate with time. We did not simulate any camera trap effect on the detectability of the encounters/associations. Although our simulated animals were characterized by a variable home range size, they all had the same activity rhythm. There were no differences between male and female, or young and adults. Animals were not gregarious (simulating gregarious animals would have led to a greater variance of the encounter rate across camera traps). In other words, it is expected that the uncertainty characterizing the population size and trends estimates in actual studies will be much more important than those obtained in our simulations. And yet our results concerning the roe deer show that the estimation of population size and trends are very imprecise. Even with 4 camera traps/km^2^, the coefficient of variation of the population size estimate was about 15 to 20% and increased to 30 to 40% when only 1 trap/km^2^ were used. Similar coefficients of variations were obtained in previous studies (e.g., Palencia et al., 2022, obtained CV comprised between 34% to 75% when estimating the population size of various species with the REM, on areas comprised between 1400 to 6600 ha with 7 to 37 camera traps – corresponding to 0.3 to 1.3 camera traps/km^2^ – during periods covering 15 to 138 days). Our study showed that a strong population decrease of 20% in 5 years was not significant in 62% to 75% of the simulations when 4 traps/km^2^ were used, and this proportion rose to 82% when a density of only 1 trap/km^2^ was used. As our simulations were very optimistic, this proportion is expected to be much smaller in real studies. This shows that the use of camera traps for the monitoring of roe deer (and other ecologically similar species) population trends is probably of very limited value.

This conclusion remains robust even when relaxing the assumption of a spatially uniform animal distribution. Additional tests incorporating habitat selection under a random sampling design (Appendix S1, Section S5.4) showed that this added layer of realism only marginally affected estimation precision. Habitat selection slightly decreased the precision of population size estimates and slightly improved that of trend estimation, but ultimately, these marginal effects did not alter our main conclusion: even when accounting for more complex spatial behaviours, the monitoring design remains largely insufficient to detect meaningful biological changes within a five-year timeframe.

### Potential for monitoring large-range species

The results obtained for the large-range roe deer were more encouraging. Even with only 1 camera trap/km^2^, we were able to statistically detect a 20% decrease of the population size in 33% to 68% of the simulations, depending on the methods used. This is still a low sensitivity, given the strong simulated population decrease and the perfect knowledge of some parameters in our simulations. However, these results indicate that camera traps monitoring might not be immediately dismissed for larger species in areas of size similar to the Chizé forest. An important quantity governing the choice to use or not camera traps for population monitoring is the number of different camera traps that can capture a given animal in a given study area. This metric depends on multiple factors, including the number of traps deployed, the species’ home-range size, heterogeneity of space use within the home range, and the size of the study area. For example, considering the large-range roe deer in our study, randomly placing 100 camera traps over the 2600 ha study area resulted in a density of approximately 4 traps/km^2^, meaning that roughly 18 traps were included within the average 4.5 km^2^ home-range. As the home-range use of simulated large-range deer was highly heterogeneous – with activity concentrated around a limited number of attraction points (see Appendix S1 for an illustration) – the actual number of traps detecting a given individual was considerably lower than this value, averaging 3.5 traps per animal (Table 4). Consequently, the death of a large-range roe deer between two years is detected by an average of 3.5 traps, resulting in a clear decrease in the number of encounters/associations detected by the traps. This explains the effectiveness of camera trap monitoring for this species under these conditions. In contrast, with a larger study area of 20000 ha and the same 100 traps, the trap density would decrease to 0.5 traps/km^2^. This would result in approximately 0.5×3.5/4 = 0.44 different traps encountered by a given individual in average, similar to the value obtained for the standard roe deer in our simulations (Table 4). Indeed, for the standard roe deer, with only 0.45 traps visited per animal in Chizé, most individuals remain undetected, resulting in low power to statistically detect population declines.

We further explored this effect by conducting a small additional simulation in Appendix S1 (section S5.6). Since the the large-range roe deer encountered 7.6 times more traps on average than the standard roe deer, we simulated a design where the trap density for the standard roe deer was increased 7.6-fold (760 traps) while the study duration was reduced to 3.95 days to maintain a constant total sampling effort (as detailed in Appendix S1, section S5.6). Despite the limited number of iterations carried out for this specific scenario, this high-density, short-term design led to nearly identical precision in trend estimation to that of the large-range scenario. These findings further confirm that the number of unique traps encountered per individual is a fundamental driver of monitoring performance, regardless of whether it is achieved through larger animal movements or higher trap density.

### Management implications

While this study provided unbiased estimates of population sizes, it also revealed important limitations for monitoring roe deer populations with camera traps: low precision and a high susceptibility to spatiotemporal variations in animal behavior. The results for the large-range virtual species were more encouraging, which suggests that camera-trap population monitoring might be more suitable for species with high mobility. However, other factors may complicate this potential. Consider the red deer as an example: population units for this species are generally much larger than 2600 ha (in France, typical red deer management units used by hunters associations often exceed 10 000 ha; Maryline Pellerin, com. pers.). Should a camera-trap study be designed to monitor the red deer in such contexts, the number of camera traps per red-deer home range would likely be too low, leading to low-precision estimates. Moreover, the gregarious behaviour of this species could further exacerbate this imprecision. Thus, even if camera trap monitoring might be promising for scientific studies of such species in small areas, its practical ability for day-to-day population monitoring by wildlife managers remain questionable, even for the large-range species.

In fact, we advise the reader to proceed to simulations to determine whether or not camera traps are a suitable choice to design a monitoring a given population of a given species, and to proceed to the best adaptations. Our R package `simCTChize` can be useful to carry out such simulations. Appendix S1, also available as a vignette of the package, contains a tutorial explaining how it can be used to reach this aim.

Should simulations indicate that camera-trap monitoring is viable for a given species and study area, researchers must tailor their study design to their specific objectives. If the goal is the estimation of absolute population size, formally estimating detection probability is needed. We recommend Moeller et al. (2023) as a comprehensive starting point for integrating these protocols. However, if the objective is limited to trend estimation, detectability corrections are not necessarily required, provided that detection remains constant over time. Similarly, while encounter-based methods (e.g. REM) require an annual estimation of mean movement speed (e.g. using the framework developed by Palencia et al., 2019), practitioners can reduce operational complexity by opting for association-based approaches, such as IS or camera-trap distance sampling, which circumvent the need for speed data.

Regardless of the chosen estimator, managers must resist the temptation to place traps only in high-use habitats to maximize detections. While placing cameras in less-frequented areas may seem like a ‘wasted’ effort, our results demonstrate this spatial randomness is a prerequisite for unbiased trend detection. Ultimately, while the automated nature of camera traps is economically attractive, a scientifically sound program – requiring a high number of units and random sampling – represents a significant investment, not only financially but also in terms of human labor. Managers should therefore understand that reducing costs by limiting study objectives or trap density is only possible if the resulting loss in precision and the risk of bias remain acceptable for their conservation goals.

## Acknowledgements

We warmly thank our colleagues of the French biodiversity management organisation (*Office français de la biodiversité*), and in particular the members of the MAS network (*Mathématique appliquée et statistiques*), who provided feedback and constructive criticisms on these results.

