## Appendix S1 for "Evaluating camera trap methods for monitoring population trends in ungulates: insights from simulation"

### S1 Introduction

This vignette contains the supplementary material of Calenge et al. (2026). The aim of this paper is to assess the bias and precision in animal population size estimation based on camera traps data. We rely on simulations of a virtual population of roe deer in the study area of Chizé. A companion package named `simCTChize` contains the data and functions used for this paper, and is required to reproduce the calculations in this document. The present document is also available as a vignette of this package.

Several options are available to install this package. For all options, if the user is using Windows, it is required that they install the Rtools (available at this URL: https://cran.r-project.org/bin/windows/Rtools/).

Then, one possibility is to download the tarball from Zenodo (URL: https://doi.org/10.5281/zenodo.16989360), and then to place it in the R working directory. The installation is then simple:

```
install.packages("simCTChize_1.0.tar.gz", repos = NULL, type="source")
```

Another possibility is to install it from Github with the function `install_github` from the package `devtools` (which also needs to be installed):

```
## If devtools is not yet installed, type
install.packages("devtools")

## Install the package badgertub
devtools::install_github("ClementCalenge/simCTChize", ref="main")
```

Throughout this vignette, we suppose that the reader is familiar with the models and simulations developed in the main paper. We give the R code used for our simulations.

Our approach consists in generating realistic simulations of the movement of $N$ animals in the study area of Chize in March of each year of a 5-year period. During this period, the average home-range size of the animals increases and the population size decreases progressively. Each year, a set of $R$ camera traps are placed on the study area and record the movements of the animals. Similarly, we simulate a realistic detection process (presence of obstacles, decreasing detection probability with distance, detectability depending on the habitat type). Finally, we use the random encounter model (REM, Rowcliffe et al., 2008) and the instantaneous sampling (IS, Moeller et al., 2018) to estimate the population size of each year as well as the decreasing trend. We have considered various parameterizations for these two methods in order to estimate the population size (accounting for activity rythm of the animals or not, accounting for imperfect detection of animals or not). In this document, we present the R code that we used for these simulations.

We first load the package:

```
library(simCTChize)
```

### S2 The components of the simulations: movement and detection

#### S2.1 Simulating roe deer movements

##### S2.1.1 Description of the simulated movement process

To simulate the movements of a roe deer, we first simulate a set of attraction points in its home range. These points correspond to places where the animal will search food more intensively. The animal will alternate between three types of movements:

- *Patch-level movements*: these movements simulate an intensive use of the habitat patches in the neighborhood of the attraction point. Formally, we simulate an Ornstein-Uhlenbeck with foraging process (Fleming et al., 2014) with a very small variance, centred on the attraction point (before generating a sequence of relocations with this type of movement, the attraction point for this sequence is randomly sampled among the set of attraction points simulated at the beginning of the process).

- *Between-patch movements*: these movements simulate a less intensive space use describing movements between patches. They are also simulated with Orntein-Uhlenbeck with foraging processes characterized by a larger variance, and also centred on one randomly sampled attraction points (again, one point is sampled before generating each sequence).

- *Resting*: in this case, the animal does not move at all, and remains at the same location during the whole resting period.

The movement simulation is based on the following daily pattern. The day is supposed to start at 17:00 in the evening; for each day of the month:

- We first simulate the presence of a resting period in the middle of the day with a probability equal to 0.85. If a resting period is simulated during a particular day, its starting time $m$ is randomly drawn from a uniform distribution bounded by 8:00 and 15:00. The duration of the resting period (in hours) is equal to $d = 15 - m + e$, where $e$ is a residual randomly drawn from a Gaussian distribution with mean 0 and standard deviation 1 (truncated so that the resting period never ends after 17:00);

- We then randomly place patch-level movements in the remaining periods of the 24-h sequence: we simulate the start time of these patches movement – if any – by simulating a Poisson process with constant intensity $\lambda'(t) = 0.66/(15 \times 3600)$ (which guarantees a mean number of patches equal to 0.66 per night). Their duration is randomly drawn from a uniform distribution bounded by 0 and 5 hours.

- Finally, we randomly place "resting patches" in the remaining periods except between 6:00 and 8:00 in the morning where all animals are supposed active. We simulate the start time of these resting patches with a Poisson process with constant intensity $\lambda'(t) = 0.33/(15/3600)$. The duration of each patch is randomly drawn from a uniform distribution bounded by 0 and 5 hours.

- "between-patch" type of movements are simulated in the remaining periods.

When the animal switches to a resting period, it stops moving immediately at the beginning of the period and does not move until the end of the resting period. When the animal switches to "between-patches movement" type, an attraction point is randomly drawn from the simulated set, and an Ornstein-Uhlenbeck with foraging process is simulated with the following parameters: $\tau_r = \tau_v = 4000$ seconds and a value of $\sigma^2$ randomly drawn from a uniform distribution bounded by 1000 and 10000 $m^2$. When the animal switches to "patch movement", a new attraction point is again randomly drawn from the simulated set, and an Ornstein-Uhlenbeck with foraging process is simulated with the following parameters: $\tau_r = \tau_v = 300$ seconds, and a value of $\sigma^2$ equal to 200 $m^2$. All these parameters (of the Ornstein-Uhlenbeck processes, of the distributions controlling the occurrence and duration of the different sequences, etc.) have been set after an analysis of GPS data collected in the Chizé study area.

##### S2.1.2 Simulation in R

The package `simCTChize` contains all the required elements implementing this process. The first step of the simulation process consists in simulating a set of attraction points for the Ornstein-Uhlenbeck process. We used GPS data collected on 25 roe deer to estimate 25 sets of attraction points: we used the kernel method to estimate the utilization distribution for each animal, and we identified the local modes in this distribution (each mode corresponding to an attraction point). We used this list of 25 sets of attraction points as a basis for our simulations. Thus, to simulate the movement of a roe deer, we first sampled one of the 25 available sets that we randomly shifted over the study area. The dataset `listPatches` from the package is a list of the 25 matrices containing the coordinates of the attraction points of these 25 animals (centred so that their mean x and y coordinates are equal to zero). For example, we can plot the set of attraction points for the first animal:

```
plot(listPatches[[1]])
```

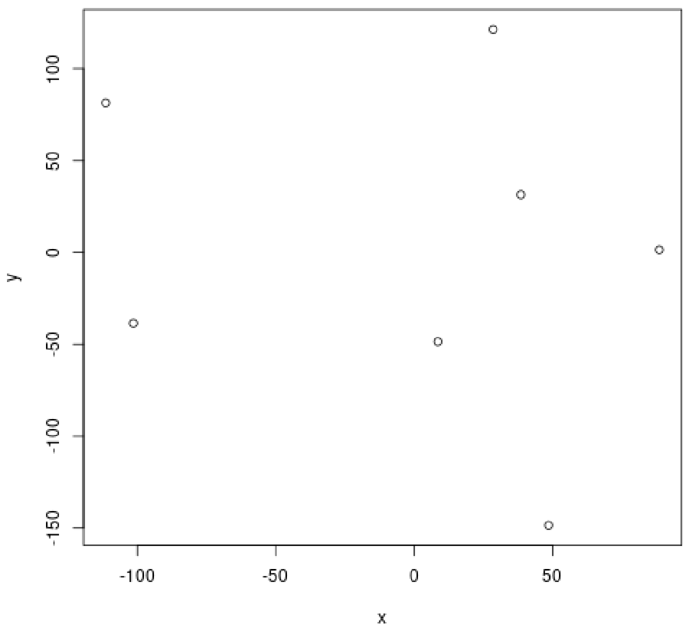

The simulation of roe deer movements requires the pre-calculation of matrices that will be used for the simulation of Ornstein-Uhlenbeck processes. These calculations are carried out by functions of the package `ctmm`, and the wrapper function `prepareMvt` from our package can be used to calculate these matrices from the values of the parameters $\tau_v, \tau_r, \sigma$ (see the help page of this function). For example, to prepare the matrices required for our simulations:

```r
## A list of matrices with different values of sigma comprised between
## 1000 and 10000 for the "between patches movement" (remember that
## sigma is randomly drawn between 1000 and 10000 m2 for this movement
## type, so that we need a list of matrices -- explaining the use of
## lapply here)
listMatBetween <- lapply(seq(1000,10000, by=1000),
                         function(x) prepareMvt(tau = c(4000,4000),
                                                sigma = x))

## The list of matrices for the "patch movement"
MatWithin <- prepareMvt(tau = c(300,300), sigma = 200)

## We store the results in a list
mvtMatrices <- list(listMatBetween=listMatBetween,
                    MatWithin=MatWithin)
```

Note that we use here internal functions from `ctmm`, which may not be available to the user in the future. To make sure that our package will still be usable even if these functions are deprecated, we stored the results of these calculations as a dataset of the package (the dataset `mvtMatrices`), so that the user does not have to worry if the above command lines no longer work on their computer.

Finally, one can use the function `roeDeerMovement` to simulate the movements of one animal during 30 days using the process described in the previous section (see the help page of this function for further information about its parameters):

```r
## Simulation of movement for 30 days:
set.seed(777) ## for reproducibility

## Lets say that we simulate movements between the attraction points
## of the first set of patches
setpat <- list(listPatches[[1]])

## We will simulate an animal with sigma=1000 for between-patches
## movements (first matrix of listMatBetween).
lipt <- list(ptg=mvtMatrices$listMatBetween[[1]],
             ptp=mvtMatrices$MatWithin)

## Simulation
rd <- roeDeerMovement(30, setpat, lipt = lipt, verbose=FALSE)

## the result:
plot(rd, ty="l", asp=1, xlab="X coordinate (metres)",
     ylab="Y coordinate (metres)")
```

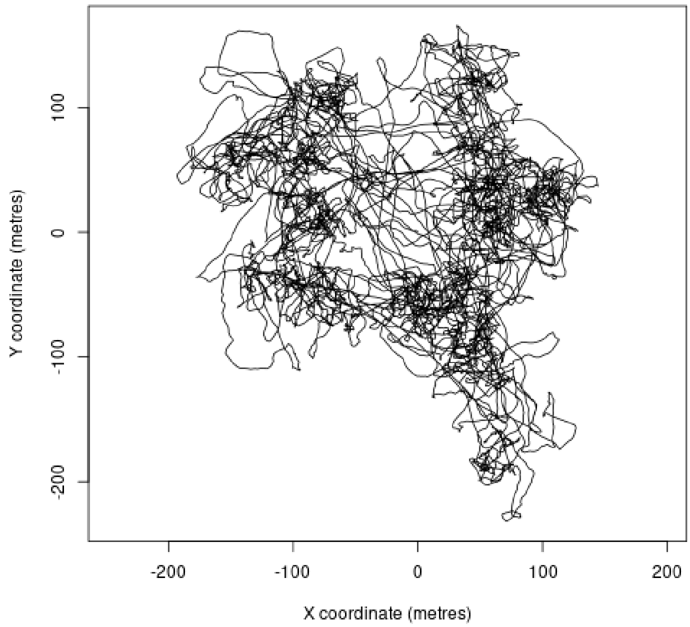

The roe deer alternates between several attraction points. Note that here the relocations are centred on (0,0).

After this simulation, the next step consists in positionning this movement in space and time. First, we have to define the coordinates of the home range centroid. In the following, we will suppose that this centroid is characterized by the coordinates 0,0:

```
## The coordinates of the centroid
coordinatesHRcentroid <- c(0,0)

## Shift the movement to this centroid
rd[,1] <- rd[,1]+coordinatesHRcentroid[1]
rd[,2] <- rd[,2]+coordinatesHRcentroid[2]
```

Of course, in this particular example, this code is useless (adding 0 to the coordinates does not change the coordinates), but this code is there to illustrate that the next step after simulation of the movement is to position it in space (in practice, `coordinatesHRcentroid` will correspond to the Lambert II coordinates of a point sampled on the study area). Finally, we must position this movement in time by defining a variable named `date` and containing the date and time of each relocation (using the format `POSIXct`, i.e. as the number of seconds elapsed since January 1st 1:00). For this example, we define simply:

```
## Transform in data.frame and add a date
rd <- as.data.frame(rd)
rd$date <- 1:nrow(rd)
```

So that the data.frame stores one relocation per second.

#### S2.2 The detection by camera traps

##### S2.2.1 Description of the simulated detection process

The previous subsection showed how we simulated random movement of the animals on the study area. The other aspect of our simulations correspond to the detection of these movements by a camera trap. In this section we describe how we simulated this detection process. First, we simulate the presence of obstacles in the neighborhood of the trap (the area located within 20 m from the trap), which will affect the detectability of the animals. The density of obstacles depends on the habitat type where the trap is placed:

- *Coppice*: in this habitat type, we simulate three types of stems: big, medium and small trees. For each type of trees we first randomly place $T$ trees with a diameter randomly drawn from a uniform distribution (comprised between 17.5 and 27.5 cm for small trees, between 27.5 and 47.5 cm for medium trees, and between 47.5 cm and 1 m for large trees), with $T$ drawn from a Poisson distribution parameterized by mean calculated from a density of stems per hectare itself randomly drawn from a uniform distribution (bounded by 50 and 70 stems per hectares for small trees, by 25 and 35 stems/ha for medium trees, and by 12 and 17 trees for large trees). These values were based on CRPF (2004).

- *Regeneration* and *Open* habitat types are characterized by the absence of obstacles. This may seem surprising for the regeneration type, but we simulated the lower detectability differently for this habitat type (by simulating a detection probability decreasing much more rapidly with the distance to the trap than for other habitat types, see below).

Once the presence of obstacle is simulated, we build a map of the detection probability in the detection zone of the trap. The orientation of the trap relative to the East direction is randomly drawn from a uniform distribution bounded by 0 and $2\pi$. The radius of the detection zone is fixed at 20 m. The angle of the detection zone is set to 0.175 radians. The detection zone is discretized in pixels of 10×10 cm. The probability of detection of an animal whose centre of gravity is located in a given pixel depends on: (i) whether this centre of gravity is located behind an obstacle or not (the animal is considered invisible behind the obstacle); (ii) whether the animal is active or not (resting animals are considered invisible); (iii) the distance between the animal and the trap. We model the detection process as a time-to-event model (aka "survival model", Therneau & Grabsch , 2000), where the detection is modeled with a hazard function. If the animal is not resting, the hazard $\lambda(t)$ that the animal is detected $t$ seconds after it stayed active at a distance $d(t)$ from the camera trap is modeled by a Cox proportional hazard model:

$$\lambda(t) = \lambda_0 \times \exp(\beta_d \times d(t))$$

where $\lambda_0$ is a baseline hazard. This can be reformulated as

$$\lambda(t) = \exp(\beta_0 + \beta_d \times d(t))$$

with $\beta_0 = \log \lambda_0$. The parameter $\beta_d$ describes how the detectability decreases when the distance to the trap increases. The cumulative risk $\Lambda(T)$ can be calculated from the hazard function:

$$\Lambda(T) = \int_0^T \lambda(t)dt$$

The probability that the animal is detected after $t$ seconds in the detection zone of the trap is then:

$$D(t) = 1 - \exp(-\Lambda(t))$$

When T is small enough, then $\Lambda(T) \approx T \times \lambda(0)$. Here, we consider time intervals of $T = 1$ seconds, so that:

$$D(t) = 1 - \exp(-\exp(\beta_0 + \beta_d \times d(t)))$$

where $d(t)$ is the distance between the animal and the trap at time $t$. In our simulations, we have fixed $\beta_0, \beta_d$ such that the probability to detect the animal in one second at small distance $d_1$ is nearly certain, set to $p_1$ (e.g. $p_1 = 0.99$), and the probability to detect it at a large distance $d_2$ is nearly 0, set to $p_2$ (e.g. $p_2 = 0.01$). We can show that:

$$\begin{aligned} \beta_d &= (\log(-\log(1-p_1)) - \log(-\log(1-p_2)))/(d_1 - d_2) \\ \beta_0 &= \log(-\log(1-p_2)) - \beta_d \times d_2 \end{aligned}$$

In our simulations, we defined different values of $\beta_0, \beta_d$ for the different habitat types:

- *Coppice* and *Open* habitat types: we chose these parameters so that the probability to detect the animal is $p_1 = 0.99$ at $d_1 = 5$ metres from the trap, and $p_2 = 0.01$ at $d_2 = 18$ metres from the trap (calibrated visually from the results of Howe et al., 2017, who worked on a species with a similar size in a similarly open environment).

- *Regeneration* habitat type: we chose these parameters so that the probability to detect the animal is $p_1 = 0.99$ at $d_1 = 0.5$ metres from the trap, and $p_2 = 0.01$ at $d_2 = 3$ metres from the trap (subjective choice).

##### S2.2.2 Implementation in R

The package `simCTChize` provides several functions to map the detectability of animals in the detection zone of a camera trap. The function `organiseCT` carries out all different steps described in the previous section (simulation of obstacles, random orientation of a trap, detectability, etc.). It takes only one argument, corresponding to the habitat type where the trap is located (see the help page of this function). For example, in the coppice:

```
set.seed(777)  ## for reproducibility
ct <- organiseCT("Coppice")
head(ct)

##         x   y       vu
## 80401 0.0 0.0 48.61074
## 80803 0.1 0.1 45.47615
## 81205 0.2 0.2 42.54369
## 81607 0.3 0.3 39.80032
## 82009 0.4 0.4 37.23385
## 82010 0.5 0.4 35.94695
```

The result is a data.frame with three columns: the coordinates x and y of the pixels in the detection zone, and a column named `vu` containing the value of the hazard function $\lambda(1)$ for each one. We can show the result on a map:

```
## For a clearer depiction of the detectability:
## log scale and rescale between 0 and 1
u <- log(ct[,3]+0.001)
v <- (u-min(u))/(max(u)-min(u))

## Then show this detectability in grey levels
plot(ct[,1],ct[,2], col=grey(v), pch=16, cex=0.7, asp=1,
     xlab="X", ylab="Y", main="Detection zone of a camera trap")
```

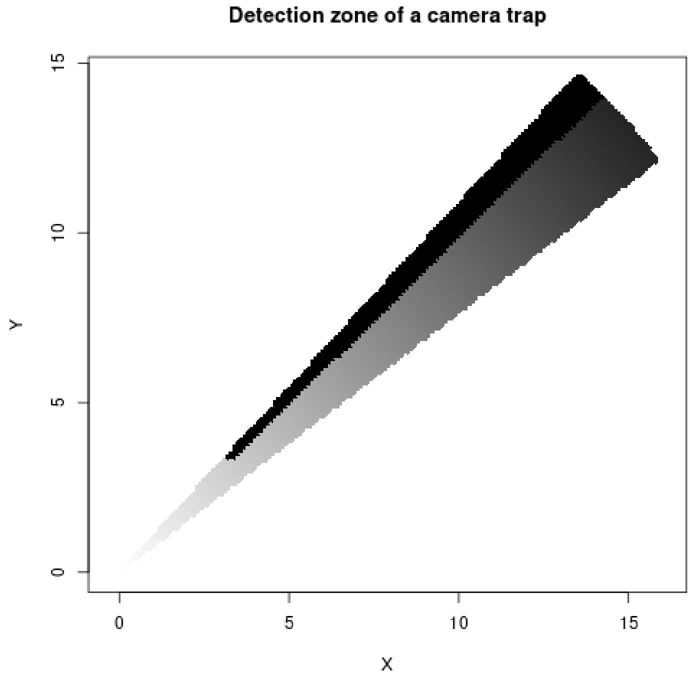

On this map, we can see that there is a tree located on the boundary of the detection zone at about 3 or 4 metres of the trap (which is located at (0,0)), that hides every pixel behind (detection risk equal to 0 – corresponding to the black area on this map). Also note that the detectability decreases with the distance to the trap and is nearly 0 at 20 metres.

*Remark*: in the examples of the help page of the function `organiseCT`, we show how the user can use the other functions of the package to simulate camera traps in other environment types (e.g. partial obstacles, different types of obstacles, different detection functions, larger detection zones, etc.).

Once the map of the detection zone has been simulated, our approach consist in placing this trap randomly on the map. This requires to sample a location for this trap. In practice in our simulations, we first sample the location of the trap, which allows to determine the habitat type where it is located (coppice, regeneration, open), and then we use `organiseCT` to simulate the detection zone. In this section, we do the reverse. Thus, after the use of `organiseCT`, we have to manually set the coordinates of the trap. To place this simulated trap in the same space as the roe deer simulated in the previous section, we place the trap at the coordinates (0,0). To define the coordinates of the trap, we set the attribute `cooori` for the object returned by `organiseCT`:

```
attr(ct, "cooori") <- c(0,0)
```

Now, for this example, we will place the movement simulated in the previous section in the same space as this trap.

```
u <- log(ct[,3]+0.001)
v <- (u-min(u))/(max(u)-min(u))
plot(ct[,1],ct[,2], col=grey(v), pch=16, cex=0.7, asp=1,
     xlab="X", ylab="Y", main="Detection zone of a camera trap")
lines(rd[,1:2], col="red")
```

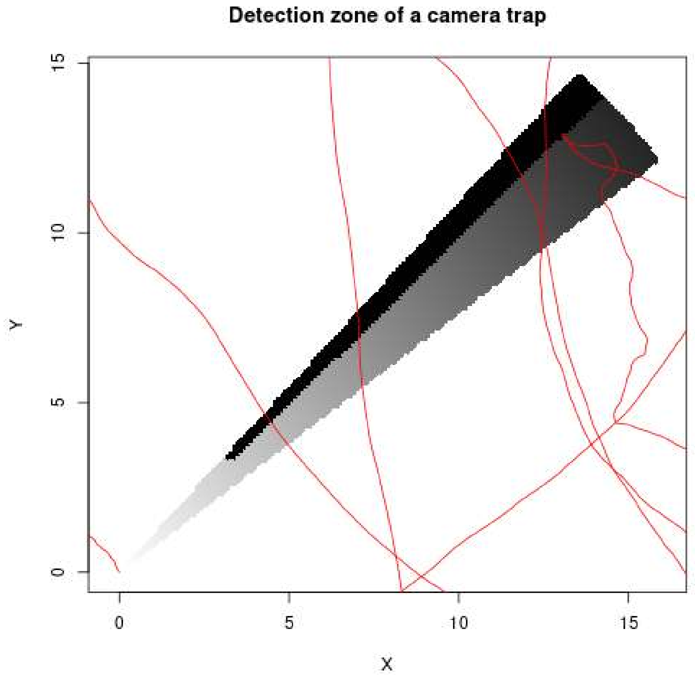

During one month, the animal crossed 5 times the detection zone of this camera trap. We can simulate the detection process with the function `whenDetection` (see the help page of this function):

```
wd <- whenDetection(as.data.frame(rd), ct)
str(wd)

## 'data.frame': 5 obs. of  7 variables:
##  $ noPassage: int  1 2 3 4 5
##  $ beginning: int  303156 312582 1526489 2022068 2370253
##  $ end      : int  304622 312942 1526546 2022142 2370440
##  $ duration : int  1467 361 58 75 188
##  $ detection: logi  TRUE TRUE TRUE ...
##  $ Pdetect  : num  1 0.992 ...
##  $ When     :List of 5
##   ..$ 1: int  303203 303317 303365 303524 303872 ...
##   ..$ 2: int  312600 312603 312659 312700 312805
##   ..$ 3: int  1526489 1526490 1526491 1526492 1526493 ...
##   ..$ 4: int  2022093 2022095 2022097 2022102 2022110 ...
##   ..$ 5: int 2370293
```

The result is a data.frame describing the number of encounters between the animal and the trap. For each encounter, this data.frame indicates the date/time of the beginning and the end of the encounter, its duration, whether the encounter was detected or not, the theoretical probability that this encounter was detected based on the map of the detection zone, as well at the dates/times when the animal has been detected (using the terminology of our paper, the date/time of the *associations*). We can show these encounters on the map:

```
u <- log(ct[,3]+0.001)
v <- (u-min(u))/(max(u)-min(u))
plot(ct[,1],ct[,2], col=grey(v), pch=16, cex=0.7, asp=1,
     xlab="X", ylab="Y", main="Detection zone of a camera trap")
lines(rd[,1:2], col="red")
tmp <- lapply(wd$When, function(x) {
    points(rd[rd$date%in%x, 1:2], pch=21, bg="yellow", col="green")
```

```
})
```

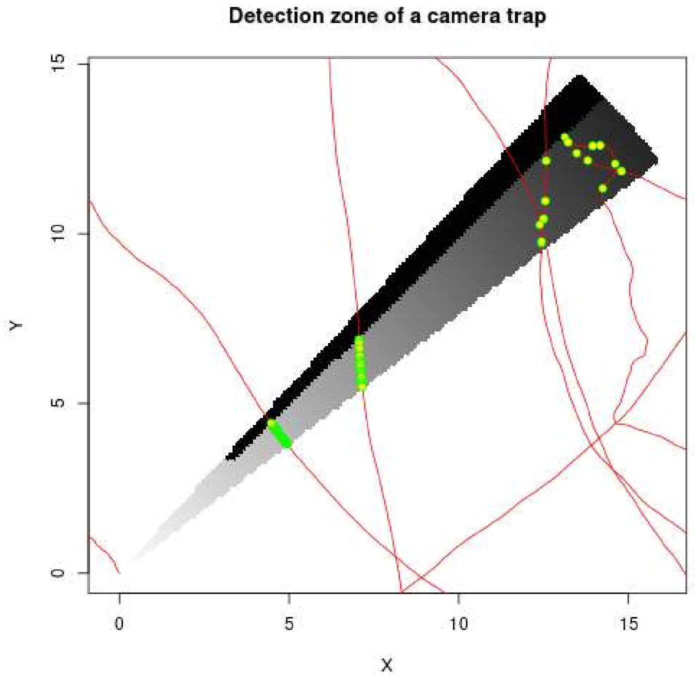

Green/yellow points correspond to the animal relocations that were detected by the camera trap. We can see that encounter occurring close to the trap are detected at every second, whereas the encounters occurring far from it are more rarely detected.

### S3 Simulating the movements of a roe deer population during five years

#### S3.1 Changes in population size

We explained in the previous section how we simulated the movement of one roe deer and how these movements were monitored using one camera trap. We now expand a bit on how we simulated the monitoring of a whole population during five years.

First, we simulated a simplified population dynamics model to generate a decrease in population size. We started with an initial population size of $N_0 = 300$ individuals with a balanced sex ratio (we ignored the slightly larger proportion of females in this population; Pellerin, com. pers). The camera trap monitoring is carried out in March. For all years, we supposed that births occur in spring, after this monitoring. We supposed that 90% of the $N_t/2$ females participate to the reproduction (slightly smaller than the 95% observed nowadays in Chizé, but our aim was to simulate a decrease in population size), and that each participating female had a probability of 0.3 to give birth to a fawn and a probability of 0.7 to give birth to two fawns (corresponding to the observed mean number of embryos per pregnant femal equal to 1.7). We suppose a balanced sex ratio among fawns, and we suppose that they survive until March of the following year with a probability of 0.3 (corresponding to fawn survival during the "bad" years in Chizé, Pellerin, com. pers. – again, our aim was to simulate a population decrease). If the fawns survive until the following year, they immediately become adult (we do not account for different age classes in our model, to keep it simple). We suppose that the adults of one year survive until the following year with a probability of 0.7 (such a survival probability is frequent in Chizé).

For the record, we show below the R code used to simulate this population dynamics:

```
set.seed(77) ## For the reproducibility
survival <- 0.7
survivalFawn <- 0.3
propParticipatingFemales <- 0.9

## What we have in March during year 1
Nmales <- 150
Nfemales <- 150
Ntot <- Nmales+Nfemales

## Reproductive females
Nfemalesrepro <- rbinom(1, Nfemales, propParticipatingFemales)

## Births: all participating females have by definition at least one fawn
## They have a second one with a probability of 0.7
Nbirths <- Nfemalesrepro+rbinom(1,Nfemalesrepro,0.7)

## Half males and half females in the births
Nyoungmales <- rbinom(1,Nbirths,0.5)
Nyoungfemales <- Nbirths-Nyoungmales

## We will need that
Nfemalesrepro <- 0
Newmales <- 0
Newfemales <- 0
Oldmales <- 0
Oldfemales <- 0

## For each year after the first one
for (i in 2:5) {

    ## Following March, survival of fawns of last year + male survival
    Newmales[i] <- rbinom(1, Nyoungmales[i-1], survivalFawn)
    Oldmales[i] <- rbinom(1, Nmales[i-1], survival)
    Nmales[i] <- Oldmales[i]+Newmales[i]

    ## Same for females
    Newfemales[i] <- rbinom(1, Nyoungfemales[i-1], survivalFawn)
    Oldfemales[i] <- rbinom(1, Nfemales[i-1], survival)
    Nfemales[i] <- Oldfemales[i]+Newfemales[i]

    ## So total population size
    Ntot[i] <- Nmales[i]+Nfemales[i]

    ## And next spring, births (same as year 1)
    Nfemalesrepro[i] <- rbinom(1, Nfemales[i], propParticipatingFemales)
    Nbirths[i] <- Nfemalesrepro[i]+rbinom(1,Nfemalesrepro[i],0.7)
    Nyoungmales[i] <- rbinom(1,Nbirths[i],0.5)
    Nyoungfemales[i] <- Nbirths[i]-Nyoungmales[i]
}

data.frame(Ntot, Old=Oldmales+Oldfemales, New=Newmales+Newfemales)

##   Ntot Old New
```

```
## 1  300   0   0
## 2  274 214  60
## 3  266 199  67
## 4  246 188  58
## 5  239 182  57
```

The resulting data.frame is actually stored in the dataset `dfPopSize` of the package:

```
dfPopSize
```

```
##   Ntot Old New
## 1  300   0   0
## 2  274 214  60
## 3  266 199  67
## 4  246 188  58
## 5  239 182  57
```

Thus, in our study, we simulated 300 individuals on the study area during the first year. Among these 300, 214 survived to the following year and 60 new animals enter the population (birth). Among the 274 individuals present on the study area during the second year, 199 survive to the third year et 67 animals enter the population. And so on.

Using this approach, we simulate a 20% decrease of the population size in 5 years. When an animal survived from one year to the next, the location of the centroid of its home range was kept constant. When a new animal appeared on the study area, its home-range centroid was randomly sampled within the study area (avoiding the immediate neighborhood of the study area boundary, to avoid the simulation of movements outside the study area, see our discussion of the function `simulateCTStudy5years` below).

#### S3.2 Changes in space use

We simulated an increasing average home-range size of the roe deer with time. Indeed, the negative correlation between home-range size and population size has been previously observed in other roe deer populations (e.g. Kjellander et al., 2004).

We simulated a change in the mean home-range size of the animals using two approaches:

- we varied the parameter $\sigma^2$ of the mixed Ornstein-Uhlenbeck process (which controls the dispersion of the movements during the between-patch sequences). For a given year $t$ comprised between 1 and 5, the value of $\sigma^2$ for a given animal was drawn from a uniform distribution bounded by 1000 $m^2$ and $(3 + \lfloor (t-1) \times 1.75 \rfloor) \times 1000$ m$^2$ (i.e., between 1000 and 3000 m$^2$ during the first year, and between 1000 and 10000 m$^2$ during year 5).

- we also varied the list of sets of attraction points used to simulate the patches. We distinguished the 19 "small sets" made by 10 attraction points or less and the 6 "large sets" made by more than 10 attraction points. During a given year $t$, we sampled for each animal a set of attraction points in a list built by the 19 small sets and a number of $\lfloor (t-1) \times 6/4 \rfloor$ randomly sampled "large" sets. Thus, as $t$ increases, the number of "large" sets increases too in the sampled list.

We illustrate how this approach led to a change in home range size by comparing the home-range size (estimated by the minimum convex polygon) of 200 animals simulated during year 1 and year 5. WARNING!!! This code takes several minutes to execute. Fortunately, the resulting data.frame `simHRspeed` is available as a dataset in the package, so that the user does not need to execute this code to access to the result:

```r
## list of Patches used for year 1 and year 5
patchesYear1 <- listPatches[sapply(listPatches,nrow)<=10]
patchesYear5 <- listPatches

## remember that the list of movement matrices contain one element per
## value of sigma2 comprised between 1000 and 10000. Thus:
mvtMatricesYear1 <- mvtMatrices
mvtMatricesYear1$listMatBetween <- mvtMatricesYear1$listMatBetween[1:3]
mvtMatricesYear5 <- mvtMatrices

## Load adehabitatHR for home-range size estimation
library(adehabitatHR)

## Initialization
simHRspeed <- list()

## Simulation
set.seed(777) ## for reproducibility

## For each year
for (year in c(1,5)) {

    homeRangeSizey <- numeric(0)
    speedy <- numeric(0)

    ## And for each animal
    for (i in 1:200) {
        cat("year", year,"animal", i,"\r")

        if (year==1) {
            lipt <- list(ptg=mvtMatricesYear1$listMatBetween[[sample(1:3,1)]],
                         ptp=mvtMatrices$MatWithin)
            patch <- patchesYear1
        } else {
            lipt <- list(ptg=mvtMatricesYear5$listMatBetween[[sample(1:10,1)]],
                         ptp=mvtMatrices$MatWithin)
            patch <- patchesYear5
        }

        z <- roeDeerMovement(30, lixy=patch, verbose=FALSE, lipt=lipt)

        ## Calculation of average speed
        speedy[i] <- mean(sqrt(rowSums((z[-1,]-z[-nrow(z),])^2)))

        ## Home-range size estimation if the animal had been monitored by
        ## GPS with one relocation every 20 min
        m <- as.integer(seq(1,nrow(z),by=60*20))
        zb <- z[m,]
        homeRangeSizey[i] <- mcp(SpatialPoints(zb))[[2]]
    }
    simHRspeed[[year]] <- data.frame(year=year, HRsize=homeRangeSizey,
                                     speed=speedy)
}

HRspeed <- do.call(rbind, simHRspeed)
```

The resulting plot shows how the home-range size varies from year 1 to year 5:

```
library(ggplot2)
ggplot(HRspeed, ggplot2::aes(y = HRsize, x = factor(year))) +
    geom_violin(draw_quantiles = c(0.1, 0.9)) +
    geom_boxplot(width = 0.2, fill = "grey",
                 outlier.shape = NA) + ylab("Home-range size (hectares)") +
    xlab("year")
dvm <- round(tapply(HRspeed$HRsize, HRspeed$year, mean), 1)
dvs <- round(tapply(HRspeed$HRsize, HRspeed$year,sd),1)
```

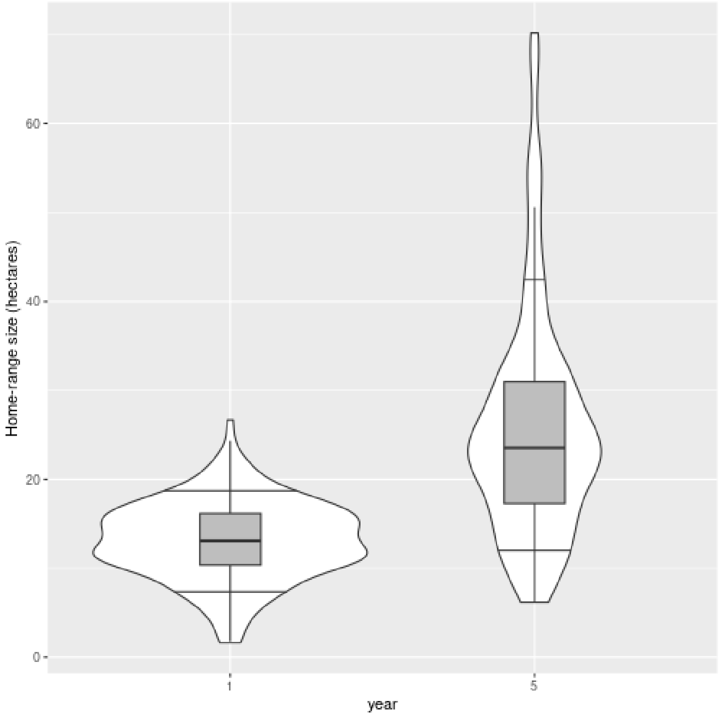

This approach leads to a mean home range size multiplied by two between year 1 and year 5. With the parameters described at the beginning of this section, the mean home-range size during year 1 is 13.1 ha (SD = 4.4 ha) and 25.8 ha (SD = 12.9 ha) during year 5.

Note that the changes in space use does not only affect the home-range size; it also affects the average movement speed of animals:

```
ggplot(HRspeed, ggplot2::aes(y = speed, x = factor(year))) +
    geom_violin(draw_quantiles = c(0.1, 0.9)) +
    geom_boxplot(width = 0.2, fill = "grey",
                 outlier.shape = NA) + ylab("Mean speed (m/s)") +
    xlab("year")
```

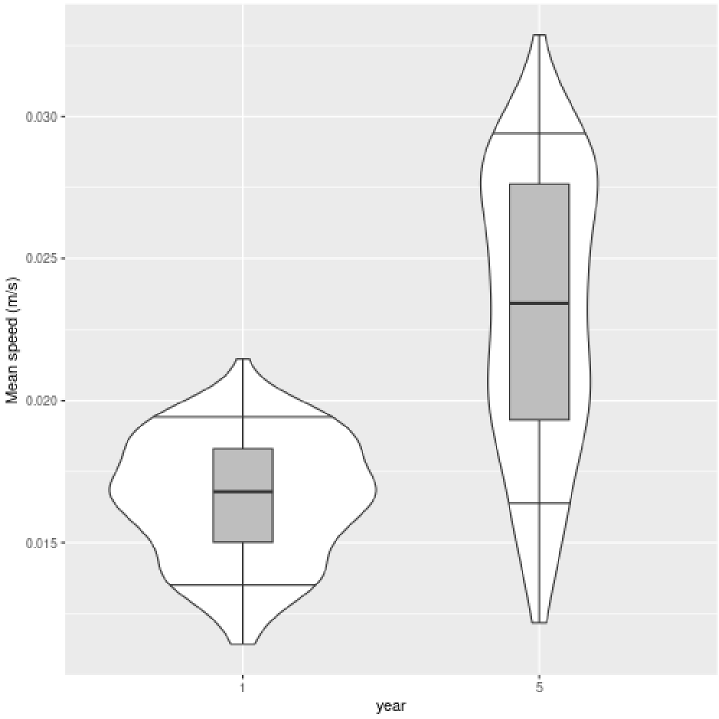

This may be important when the population size is estimated using methods relying on the mean speed (e.g., random encounter models).

##### S3.3 The maps required for the simulations

Several other elements are required for this simulation. As noted before, we simulate roe deer movements in the Chizé study area, so that we need a map of the contour of the study area (this is required to place both animals and camera traps). The coordinates of the contour of this area are available in the dataset `contourChize`. Note that in this area, the animals home ranges are not randomly distributed. A larger proportion of animals is present in the northern part of the study area (64%, Pettorelli et al., 2001). We reproduced this heterogenous distribution of animals in our study, by placing each simulated animal in the northern part with a probability of 0.64. The coordinates of the northern and southern parts of the study area are available as the datasets `northChize` and `southChize` respectively. We show these three maps below:

```
plot(contourChize, type="n", xlab="X coordinates (Lambert II)",
     ylab="Y coordinates (Lambert II)", main="Chize study area")
polygon(contourChize, lwd=3)
polygon(northChize, col="lightblue")
polygon(southChize, col="orange")
```

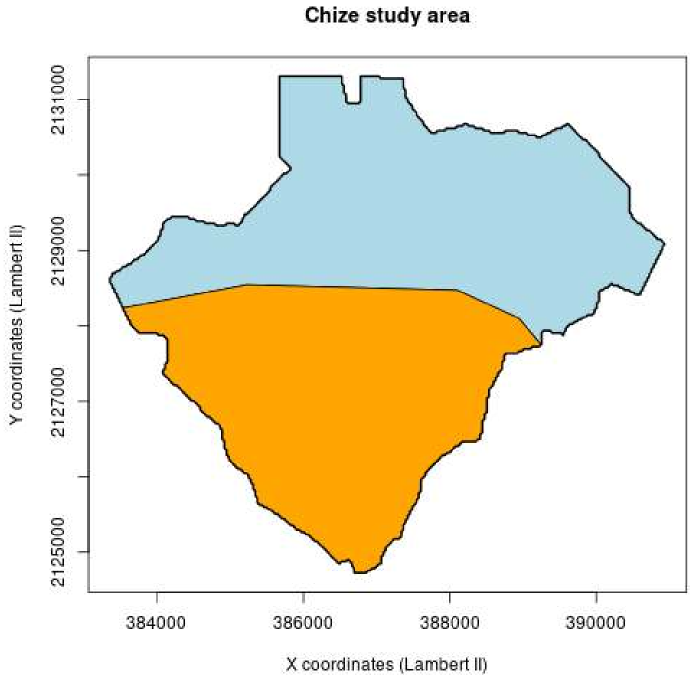

When randomly placing camera traps over the study area, we need to account for the habitat type where they are located. The dataset `habitatMap` contains an object of the class `sf` (from the package of the same name) with this information.

```
library(sf)
plot(habitatMap)
```

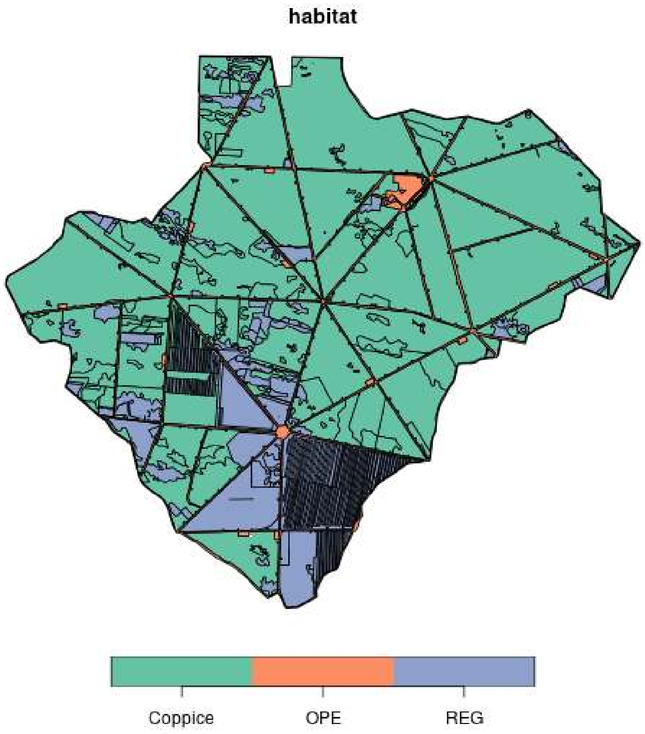

#### S3.4 The function used for the simulation

We now have all the elements required to simulate the monitoring of the population during 5 years by a camera trap study carried out every year in March. The function `simulateCTStudy5years` implements this procedure. It uses internally the functions described in the previous sections to implement the monitoring. The code below shows how to use this function to simulate 4 iterations of such a study. WARNING!!! THIS FUNCTION IS VERY SLOW!!!!! It took 40 minutes to carry out these 4 simulations. We provide the results of this simulations in the dataset `sim4iter` of the package, so that the user does not need to execute this command to access the result:

```
sim4iter <- simulateCTStudy5years(nCT = 100, duration = 30,
                                  niter = 4, dfPopSize,
                                  listPatches, contourChize, contourN=northChize,
                                  contourS=southChize,
                                  dff=relationAnimalSigmaMaxd, habitatMap,
                                  listptg=mvtMatrices$listMatBetween,
                                  ptp=mvtMatrices$MatWithin, nofi = 0, backup = TRUE)
```

We simulate here the monitoring of the population with 100 camera traps during 30 days (one month). Four iterations of the 5-year study are carried out, and the data.frame `dfPopSize` is passed as an argument to indicate how the population changes over the 5 years. We also pass the list of sets of attraction points (for the patches), the different maps, the movement matrices.

*Remark*: the argument `dff` (set to the dataset `relationAnimalSigmaMaxd`, available in the package) is a data.frame containing the maximum predicted distance between the animal movement and its home-range centroid as a function of the animal id (each id corresponding to the corresponding set of attraction points in the argument `listPatches`) and of a value of $\sigma$ (where the actual value of sigma used for the OUf process is this value times 1000). This data.frame is used internally to select a centroid for each animal so that all its simulated relocations are located within the limits of the study area.

We refer the reader to the help page of the function for further details on this function. The `print` method allows to give a short description of this list:

```
sim4iter

##
## ******* Object of class simCT:
##
## Number of simulations of a 5-year study: 4
## Number of camera traps: 100
## Mean radius of the detection zone: 20 metres
## Mean angle of the detection zone: 0.175 radians
## Number of simulated animals for each year:
## [1] 300 274 266 246 239
## Size of the object:
## 10.1 Mb
##
## This object is a list with the following structure:
## List of 3
##  $ encounters :List of 4
##  $ speed      :List of 4
##  $ activeSpeed:List of 4
##  - attr(*, "class")= chr "simCT"
##  - attr(*, "CTparameters")=List of 2
## See ?simulateCTStudy5years for further details on these components
##
## *******
```

This list has three elements: the component `speed` contain the mean speed of the simulated animals on the study area for each iteration. The component `activeSpeed` contains the mean speed of these animals during active periods (i.e. excluding the period between 8:00 and 17:00 every day). Finally, the component `encounter` contains the main results of the study:

```
str(sim4iter$encounters,2)

## List of 4
##  $ :List of 3
##   ..$ creationCT : Named int [1:2] 687857406 695267170
##   .. ..- attr(*, "names")= chr [1:2] "position" "content"
##   ..$ indicetraps:'data.frame': 100 obs. of  3 variables:
##   ..$ results    :List of 5
##  $ :List of 3
##   ..$ creationCT : Named int [1:2] 492192608 10700035
##   .. ..- attr(*, "names")= chr [1:2] "position" "content"
##   ..$ indicetraps:'data.frame': 100 obs. of  3 variables:
##   ..$ results    :List of 5
##  $ :List of 3
##   ..$ creationCT : Named int [1:2] 345115572 345015853
##   .. ..- attr(*, "names")= chr [1:2] "position" "content"
##   ..$ indicetraps:'data.frame': 100 obs. of  3 variables:
##   ..$ results    :List of 5
##  $ :List of 3
##   ..$ creationCT : Named int [1:2] 995049911 172049480
##   .. ..- attr(*, "names")= chr [1:2] "position" "content"
##   ..$ indicetraps:'data.frame': 100 obs. of  3 variables:
##   ..$ results    :List of 5
```

This component is itself a list containing three elements: (i) a vector named `creationCT` with two components (the seed used by the random number generator to generate the position of the camera traps on the area, and the seed used by this generator to generate the position of trees – obstacles – in the detection zone), (ii) a data.frame named `indicetraps` containing the coordinates x and y of the camera traps, as well as the habitat type in which they fall, (iii) a list named `results` containing the encounter and association detections. This component contains the result for the five years of the study, each year being a list of data.frames. Each data.frame corresponds to the set of encounters of one animal by one camera trap. For example, considering the first pair of animal/camera trap during the first year:

```
str(sim4iter$encounters[[1]]$results[[1]][[1]])

## 'data.frame': 3 obs. of  9 variables:
##  $ noPassage: int  1 2 3
##  $ beginning: int  268719 277145 598774
##  $ end      : int  268755 277194 598861
##  $ duration : int  37 50 88
##  $ detection: logi  TRUE FALSE TRUE
##  $ Pdetect  : num  0.194 0.46 ...
##  $ When     :List of 3
##   ..$ 1: int 268743
##   ..$ 2: int
##   ..$ 3: int  598775 598785 598798 598803 598804 ...
##  $ whichani : int  3 3 3
##  $ whichCT  : int  16 16 16
```

The columns `whichani` and `whichCT` indicate respectively the animal and camera traps ID. Each row of this data.frame correspond to one encounter of the animal by the camera trap. This data.frame is similar to the one returned by the function `whenDetection` (actually it is, we just added the two columns `whichani` and `whichCT`).

### S4 Estimating the population size from the simulations

#### S4.1 The population size itself

The function `simulateCTStudy5years` simulates a certain number of 5-year camera trap studies and stores all encounters and associations detected by the camera traps, as well as the mean speed of animals during the whole period and during the active period. These data can then be used to estimate the population size for each year of the study period. Two methods are implemented in `simCTChize`: random encounter model (REM, Rowcliffe et al., 2008), implemented in the function `estimRem` and instantaneous sampling (IS, Moeller et al., 2018), implemented in the function `estimIS`.

In our paper we considered six methods to estimate population size:

- `REM_raw`: the data used for the random encounter model correspond to all detected encounters. However, we do not account for imperfect detection in the estimation, and we do not account for the activity rythm of animals.
- `REM_detect`: the data used for the random encounter model correspond to all detected encounters, and we estimate population size while accounting for the detection probability estimated from the data.
- `REM_detect_active`: the data used for the random encounter model correspond only to the detected encounters during the most active periods (i.e. excepting the period comprised between 8:00 and 17:00), and we estimate the detection probability from these data. In addition we use the mean speed of animals calculated during active periods.
- `IS_raw`: this approach uses the instantaneous sampling approach on all the detected associations, but supposes that the detection probability is equal to 1.
- `IS_detect`: this approach is similar to the previous one, but corrects for the imperfect detection using the estimated detection probability for each trap.
- `IS_detect_active`: this approach is similar to the previous one, but relies only on the data collected during active periods (i.e., excluding the period comprised between 8:00 and 17:00 every day).

##### S4.1.1 Random encounter model

We first illustrate the use of `estimRem` to estimate the population size based on the results of `simulateCTStudy5years`. As an example, we estimate population size during the first year of the study carried out in the first iteration of the object `sim4iter` returned in the last section:

```
firstyear <- sim4iter$encounters[[1]]$results[[1]]
str(head(firstyear,7),1)

## List of 7
##  $ :'data.frame': 3 obs. of  9 variables:
##  $ :'data.frame': 3 obs. of  9 variables:
```

```
##  $ :'data.frame': 2 obs. of  9 variables:
##  $ :'data.frame': 1 obs. of  9 variables:
##  $ :'data.frame': 5 obs. of  9 variables:
##  $ :'data.frame': 2 obs. of  9 variables:
##  $ :'data.frame': 11 obs. of  9 variables:
```

Remember that `firstyear` is a list of data.frame, each one containing the set of encounters between one animal of the population and one camera trap. We bind the rows of the different data.frames to obtain a unique data.frame containing all the encounters between the animals of the population and the camera traps:

```
firstyeardf <- do.call(rbind, firstyear)
str(head(firstyeardf))

## 'data.frame': 6 obs. of  9 variables:
##  $ noPassage: int  1 2 3 1 2 ...
##  $ beginning: int  268719 277145 598774 1597812 1612362 ...
##  $ end      : int  268755 277194 598861 1598025 1612535 ...
##  $ duration : int  37 50 88 214 174 ...
##  $ detection: logi  TRUE FALSE TRUE ...
##  $ Pdetect  : num  0.194 0.46 ...
##  $ When     :List of 6
##   ..$ 1: int 268743
##   ..$ 2: int
##   ..$ 3: int  598775 598785 598798 598803 598804 ...
##   ..$ 1: int
##   ..$ 2: int
##   ..$ 3: int
##  $ whichani : int  3 3 3 6 6 ...
##  $ whichCT  : int  16 16 16 34 34 ...
```

To estimate population size with random encounter models, we need to calculate the mean speed of animals over the study area. We have this information in the component `speed` of the object returned by `simulateCTStudy5years`. Let us calculate the mean speed of the animals during the first year of the first iteration:

```
(dr1 <-  mean(sim4iter$speed[[1]][[1]]))

## [1] 0.01657088
```

This speed is expressed in metres/second. We can calculate the same mean speed, but focusing only on active periods (excluding the monitoring between 8:00 and 17:00):

```
(dra1 <-  mean(sim4iter$activeSpeed[[1]][[1]]))

## [1] 0.01885334
```

We can now estimate the population size on the study area from these data with `estimRem`. We pass as arguments the data.frame containing all the encounters `firstyeardf`, the mean speed `dr1`, the radius of the detection zone (20 metres), the angle of this zone (0.175 radians), as well as the duration of the study (30 days) and the number of traps:

```
(estd <- estimRem(firstyeardf, dr1, 20, 0.175, duration=30, nCT=100))

## [1] 220.6012
```

Remember that we simulated 300 animals on the study area during the first year. The population size seems underestimated here. Indeed, this approach does not account for imperfect detectability. The function `estimRem` has a parameter named `Pdetect` that can be used to specify a probability of detection of encounters during the monitoring. In an actual camera trap study, this probability should be estimated with a field study. Here, each row of the data.frame `firstyeardf` corresponds to an encounter between an animal and a camera trap, which may be either detected or not. Therefore, in the particular case of our simulations, we can estimate the probability of detection by calculating the proportion of detected encounters:

```
(pdetect <- mean(firstyeardf$detection))

## [1] 0.6653491
```

We estimate again the population size accounting for this probability:

```
(estd2 <- estimRem(firstyeardf, dr1, 20, 0.175,
                   Pdetect=pdetect, duration=30, nCT=100))

## [1] 331.5571
```

Finally, note that the mean speed here is underestimated, leading to an overestimation of the population size, as this speed is calculated over the whole study period, including during the day, when most animals are inactive. A better estimate can be obtained by replacing the mean speed `dr1` by the mean speed during active periods `dra1`, and excluding all detections that occurred between 8:00 and 17:00 (when most animals are inactive). We also indicate that our monitoring is carried out only 15 hours per day:

```
firstyeardfactive <- removeHour(firstyeardf, 8,17)
## recalculate detection probability
(pdetect <- mean(firstyeardfactive$detection))

## [1] 0.6730769

(estd3 <- estimRem(firstyeardfactive, dra1, 20, 0.175, hoursPerDay=15,
                   Pdetect=pdetect, duration=30, nCT=100))

## [1] 319.4459
```

This leads to a better population size estimate. We can loop over all four iterations to estimate population size during the first year of each study:

```
(edr <- sapply(1:4, function(i) {
    firstyear <- sim4iter$encounters[[i]]$results[[1]]
    firstyeardf <- do.call(rbind, firstyear)
    firstyeardfactive <- removeHour(firstyeardf, 8,17)
    dra1 <-  mean(sim4iter$activeSpeed[[i]][[1]])
    (pdetect <- mean(firstyeardfactive$detection))
    estimRem(firstyeardfactive, dra1, 20, 0.175, hoursPerDay=15,
             Pdetect=pdetect, duration=30, nCT=100)
}))

## [1] 319.4459 264.9511 329.3585 353.2001
```

The mean of these four estimates is not far from the true simulated value of 300 animals:

```r
mean(edr)

## [1] 316.7389
```

##### S4.1.2 Instantaneous sampling

We now illustrate how to use the function `estimIS` to estimate the population size using instantaneous sampling. As for the random encounter model, we first focus on the first year of the first simulation:

```r
firstyeardf <- do.call(rbind, sim4iter$encounters[[1]]$results[[1]])
```

Instantaneous sampling requires an estimation of the probability of detection of associations for each trap. Be careful here: the probability of detection of associations is not the same as the probability of detection of encounters. As for the REM, in real studies, this mean probability of detection for each trap should be estimated with field studies (or alternatively, using approaches implying camera trap distance sampling). In our simulations, we have all the information required to estimate directly this mean probability of detection for each trap. The idea is to use the function `organiseCT` to map the detectability in the detection zone of each trap. Because this function requires the habitat type for each trap, we need to get this information from our simulations. This information is stored in the column `hab` of the data.frame `indicetraps` from the object returned by `simulateCTStudy5years`

```r
## habitat of the camera traps for the first simulation
habitat4sim <- sim4iter$encounters[[1]]$indicetraps$hab
str(habitat4sim)

##  chr [1:100] "Coppice" "Coppice" "Coppice" "Coppice" ...
```

Then, we need to simulate again the maps of surroundings of camera traps. This implies random elements, such as the orientation of the camera trap or the presence of obstacles. To reproduce the random environments simulated by `simulateCTStudy5years`, we need to set the random number generator of R to the same state as that used by this function. The object returned by this function includes, in the vector `creationCT`, the seeds used to simulate these environments. Therefore, we can build again a list of 100 elements corresponding to the raster maps of the detection zones of the 100 camera traps:

```r
## Seed used for the "content" of the detection zone
set.seed(sim4iter$encounters[[1]]$creationCT[2])

## Calculate raster maps of the 100 CT
list100CT <- lapply(1:100, function(j) {
    cat("Traps: ", j,"\r")
    organiseCT(habitat4sim[j])
})

## Calculation of the mean detectability for each trap:
prop <- sapply(1:length(list100CT), function(trap)
    mean(1-exp(-list100CT[[trap]]$vu)))
```

We can then use the function `estimIS` to estimate the population size using the data.frame of detected encounters `firstyeardf`, the radius (20 m) and angle (0.175 radians) of the detection zone, estimated detection probability `prop`, the number of camera traps (100) and the duration in days of the monitoring (30):

```
(edi1 <- estimIS(firstyeardf, r=20, theta=0.175, detct=prop,
                 nCT=100, duration =30))

## [1] 277.7033
```

We can improve this estimation by removing the period between 8 and 17 hour, during which most animals are inactive:

```
firstyeardfactive <- removeHour(firstyeardf, 8, 17)
```

We estimate again the population size, but this time, we indicate that there are only 15 hours of monitoring per day:

```
(edi1 <- estimIS(firstyeardfactive, r=20, theta=0.175, detct=prop,
                 nCT=100, duration =30, hoursPerDay=15))

## [1] 313.3894
```

As for the REM, we can loop over all the iterations in `sim4iter` to estimate the population size for each one:

```
edi <- sapply(1:4, function(i) {
    cat("Iteration:", i, "\n")
    firstyeardf <- do.call(rbind, sim4iter$encounters[[i]]$results[[1]])
    fyda <- removeHour(firstyeardf, 8, 17)

    ## habitat
    habitat4sim <- sim4iter$encounters[[i]]$indicetraps$hab

    ## Camera traps
    set.seed(sim4iter$encounters[[i]]$creationCT[2])
    list100CT <- lapply(1:100, function(j) {
        cat("Traps: ", j,"\r")
        organiseCT(habitat4sim[j])
    })
    prop <- sapply(1:length(list100CT), function(trap)
        mean(1-exp(-list100CT[[trap]]$vu)))

    ## estimation
    estimIS(fyda, r=20, theta=0.175, detct=prop,
            nCT=100, duration =30, hoursPerDay=15)

})
```

The results are characterized by a much larger variance than REM:

```
edi

## [1] 313.3894 182.2096 259.0404 322.5617

mean(edi)

## [1] 269.3003
```

##### S4.1.3 Uncertainty on the population size estimation

Consider the data collected by a camera trap study during the first year of the study. As noted before, we can estimate the population size using `estimRem`, for example, accounting for imperfect detection and focusing only on active periods:

```
(estd3 <- estimRem(firstyeardf, dra1, 20, 0.175,
                   Pdetect=pdetect, duration=30, nCT=100))

## [1] 288.0718
```

However, this is a point estimate, and there is an associated uncertainty to this estimate. The package `simCTChize` provides the function `bootstrapREM` to estimate the uncertainty on this estimate using a bootstrap approach (by bootstraping the camera traps). For example, using 1000 bootstrap samples:

```
br <- bootstrapREM(firstyeardf, Nboot=1000, dra1, 20, 0.175)
head(br)

## [1] 275.8369 217.5533 233.1341 271.7974 344.5075 283.9158
```

We can estimate the coefficient of variation of our estimation:

```
sd(br)/mean(br)

## [1] 0.1407838
```

Similarly, the function `bootstrapIS` can be used to estimate the uncertainty on the population size estimation by instantaneous sampling:

```
bi <- bootstrapIS(firstyeardfactive, Nboot=1000, r=20, theta=0.175, detct=prop,
                  hoursPerDay = 15, nCT = 100)
sd(bi)/mean(bi)

## [1] 0.1479277
```

#### S4.2 A summary of the changes in population density

##### S4.2.1 Comparing two years

Now imagine that we want to compare two years, to assess whether population size increased or decreased between the two years. For example, consider the IS estimation of population size in year 1 and year 5, for the first iteration of our simulations:

```
## Elements required for year 1
year1df <- do.call(rbind, sim4iter$encounters[[1]]$results[[1]])
fyda1 <- removeHour(year1df, 8, 17)

## Elements required for year 5
year5df <- do.call(rbind, sim4iter$encounters[[1]]$results[[5]])
fyda5 <- removeHour(year5df, 8, 17)

## Common elements for the two years:
## Camera traps detection probabilities
```

```
habitat4sim <- sim4iter$encounters[[1]]$indicetraps$hab
set.seed(sim4iter$encounters[[1]]$creationCT[2])
list100CT <- lapply(1:100, function(j) {
    cat("Traps: ", j,"\r")
    organiseCT(habitat4sim[j])
})
prop <- sapply(1:length(list100CT), function(trap)
    mean(1-exp(-list100CT[[trap]]$vu)))

## estimation year1
estyear1 <- estimIS(fyda1, r=20, theta=0.175, detct=prop,
                    nCT=100, duration =30, hoursPerDay=15)

## estimation year5
estyear5 <- estimIS(fyda5, r=20, theta=0.175, detct=prop,
                    nCT=100, duration =30, hoursPerDay=15)

## estimation CV year1
byear1 <- bootstrapIS(fyda1, Nboot=1000, r=20, theta=0.175, detct=prop,
                      hoursPerDay = 15, nCT = 100)
cvyear1 <- sd(byear1)/mean(byear1)

## estimation CV year5
byear5 <- bootstrapIS(fyda5, Nboot=1000, r=20, theta=0.175, detct=prop,
                      hoursPerDay = 15, nCT = 100)
cvyear5 <- sd(byear5)/mean(byear5)
```

The resulting estimations for the two years (with their coefficients of variation) are:

```
## Year 1
estyear1

## [1] 313.3894

## CV year 1:
cvyear1

## [1] 0.1493469

## Year 5
estyear5

## [1] 238.929

## CV year 5:
cvyear5

## [1] 0.2457194
```

The estimated population size decreases between the two years. It is not clear from the coefficients of variation whether this decrease is significant. We can calculate the rate of change between the two estimates:

```
(estyear5-estyear1)/estyear1

## [1] -0.237597
```

Remember that the simulated rate of change was equal to 20% so that this estimate is not very far from the truth. The function `bootstrapIScompar` can be used to calculate 1000 bootstrap replicates of this rate of change:

```
brc <- bootstrapIScompar(fyda1, fyda5, Nboot=1000, r=20, theta=0.175,
                         detct=prop, hoursPerDay=15,
                         nCT=100)
```

We can derive a 90% confidence interval on this rate of change:

```
quantile(brc, c(0.05,0.95))

##         5%        95%
## -0.5300082  0.1597203
```

The confidence interval on the rate of change includes 0, so that we are unable to conclude from these estimate that the population decreased between year 1 and year 5. This confirms the large uncertainty on this estimation.

Note that we can similarly use `bootstrapREMcompar` to compare the population sizes by the REM between the two years for the first simulation. The population size estimates are, for year 1 and year 5:

```
## mean active speed
dra1 <-  mean(sim4iter$activeSpeed[[1]][[1]])
dra5 <-  mean(sim4iter$activeSpeed[[1]][[5]])

(esty1 <- estimRem(fyda1, dra1, 20, 0.175,
                   Pdetect=pdetect, duration=30,
                   nCT=100, hoursPerDay=15))

## [1] 319.4459

(esty5 <- estimRem(fyda5, dra5, 20, 0.175,
                   Pdetect=pdetect, duration=30,
                   nCT=100, hoursPerDay=15))

## [1] 236.5377
```

And the estimated rate of change is:

```
(esty5-esty1)/esty1

## [1] -0.2595375
```

Not far from the truth. The bootstrap confidence interval on this estimate is:

```
brcr <- bootstrapREMcompar(fyda1, fyda5, Nboot=1000, dra1, dra5,
                           r=20, theta=0.175, hoursPerDay=15,
                           nCT=100,
```

```
                             detection=TRUE)

quantile(brcr, c(0.05,0.95))

##         5%        95%
## -0.5346948 -0.1021273
```

The REM approach seems more precise: we can conclude that the population decreased from the REM estimates (but remember that we suppose here that we know exactly the mean speed of animals, which will never be the case in real studies).

##### S4.2.2 A trend summary over 5 years

Our study also estimates the trend of population size over 5 years. We used the classical theoretical framework developed by Gerrodette (1987), assuming:

$$N_t = N_0 \mu^t$$

Here, $\mu$ is a growth rate for the population, and $N_0$ is the population size during the first year. This rate can be estimated using a simple linear regression. Taking the logarithm of $N_t$ in this equation, we can fit the linear regression of $\log N_t$ on $t$:

$$\log N_t = \log N_0 + (\log \mu)t + \epsilon_t$$

where $\epsilon_t$ is a Gaussian residual. The coefficient associated with $t$ is $\log \mu$. Thus, based on the classical equations of linear regression, the growth rate can be estimated with:

$$\mu = \exp \left\{ \frac{\mathrm{Cov}(\log N_t, t)}{\mathrm{Var}(t)} \right\} \quad (1)$$

Thus, considering the simulated changes in population size:

```
dfPopSize

##   Ntot Old New
## 1  300   0   0
## 2  274 214  60
## 3  266 199  67
## 4  246 188  58
## 5  239 182  57
```

We can estimate $\mu$:

```
exp(cov(log(dfPopSize$Ntot), 1:5)/var(1:5))

## [1] 0.945309
```

We found clearer to present $\lambda = 1 - \mu$, which corresponds to the proportion of the population disappearing in one year, that is:

```
1-exp(cov(log(dfPopSize$Ntot), 1:5)/var(1:5))

## [1] 0.05469102
```

The package `simCTChize` provides functions to estimate this growth rate from camera trap data, using the object returned by the function `simulateCTStudy5years`. The function `estimTrendIS` estimates the population size for each year using camera trap data, and then uses the equation 1 on estimated population sizes to estimate $\hat{\lambda}$. We illustrate the process below. We consider the first simulation from the object `sim4iter` calculated before. We extract from this object the five data.frames containing the encounters detected by the camera traps during the five years of the study:

```
listencountersAll <- lapply(sim4iter$encounters[[1]]$results,
                            function(x) do.call(rbind,x))
```

Note, as before, that it may be suitable to remove all detections that occurred between 8AM and 5PM, as the animals are less active at that time:

```
listencounters <- lapply(listencountersAll, function(x)
                         removeHour(x, 8, 17))
```

We again calculate the mean detection probability for each trap:

```
## Common elements for the five years:
## Camera traps detection probabilities
habitat4sim <- sim4iter$encounters[[1]]$indicetraps$hab
set.seed(sim4iter$encounters[[1]]$creationCT[2])
list100CT <- lapply(1:100, function(j) {
    cat("Traps: ", j,"\r")
    organiseCT(habitat4sim[j])
})
prop <- sapply(1:length(list100CT), function(trap)
    mean(1-exp(-list100CT[[trap]]$vu)))
```

The vector `prop` contains the mean detection probability for each trap. We can use the function `estimTrendIS` to estimate the detection:

```
(tr <- estimTrendIS(listencounters, r=20, theta=0.175, detct=prop,
                    hoursPerDay=15))

##         N[1]         N[2]         N[3]         N[4]
## 313.38935378 322.65506702 208.96031966 209.56266005
##         N[5]       lambda
## 238.92896852  -0.09281794
```

This function returns the population size estimate for each year as well as the estimate of $\lambda$. Here, we estimate a decrease of 9.28% per year. The function `bootstrapTrendIS` can be used to calculate the same vector for `Nboot` = 1000 bootstrap samples:

```
set.seed(777) ## for reproducibility

bos <- bootstrapTrendIS(listencounters, r=20, theta=0.175, detct=prop,
                        hoursPerDay=15)
```

The result is a matrix with as many rows as bootstrap samples:

```
head(bos)

##          N[1]     N[2]     N[3]     N[4]     N[5]
## [1,] 277.7425 256.1204 256.1702 214.0945 222.2812
## [2,] 281.5916 264.3492 287.5678 271.3410 349.1259
## [3,] 278.2501 314.3660 223.8777 229.2652 302.1663
```

```
## [4,] 305.3056 290.6494 145.3702 194.8218 234.9621
## [5,] 347.3423 393.0681 184.9712 166.1065 279.5342
## [6,] 250.3980 244.2740 215.0479 161.1085 321.7679
##             lambda
## [1,] -0.060561842
## [2,]  0.046661472
## [3,] -0.014963346
## [4,] -0.088242326
## [5,] -0.121529207
## [6,]  0.008571026
```

So that a 90% confidence interval can simply be calculated with:

```
qutis <- apply(bos,2,quantile, c(0.05, 0.95))
```

Which is equal to:

```
qutis

##         N[1]     N[2]     N[3]     N[4]     N[5]
## 5%  237.5351 232.2584 146.1053 143.3246 154.0484
## 95% 388.2611 426.6904 270.8308 276.3615 341.5609
##           lambda
## 5%  -0.188478907
## 95%  0.008169133
```

We have one confidence interval for each population size estimate, as well as for the trend $\lambda$.

We can similarly use the function `estimTrendRem` to estimate the trend using the random encounter model. We first need to calculate the mean speed for each year (remember that we are interested here in active periods):

```
vectorSpeed <- sapply(sim4iter$activeSpeed[[1]], mean)
vectorSpeed

## [1] 0.01885334 0.01957492 0.02244776 0.02449569 0.02662553
```

Then we use the function `estimTrendRem`:

```
estimTrendRem(listencounters, vectorSpeed, r=20, theta=0.175,
              hoursPerDay=15,
              Pdetect=mean(do.call(rbind,listencounters)$detection))

##        N[1]        N[2]        N[3]        N[4]        N[5]
## 291.6958701 272.1135294 228.1892469 232.8452590 215.9898598
##      lambda
##  -0.0728877
```

And we can use the function `bootstrapTrendRem` to estimate confidence intervals:

```
set.seed(777)
bosr <- bootstrapTrendRem(listencounters, vectorSpeed, r=20,
                          theta=0.175,
                          hoursPerDay=15,
                          Pdetect=mean(do.call(rbind,listencounters)$detection))
citr <- apply(bosr,2,quantile,c(0.05,0.95))
```

Which gives:

```
citr

##         N[1]     N[2]     N[3]     N[4]     N[5]
## 5%  229.1896 196.6602 165.8572 166.0386 151.6650
## 95% 356.7024 350.0151 299.6209 309.2095 282.7048
##          lambda
## 5%  -0.15595303
## 95%  0.01005886
```

#### S4.3 A summary

The function `processCTStudy` provides a summary of the population size estimates and trends obtained by IS or REM for each simulation. The help page of this function describes how to use this function. We illustrate it in this section. For example, consider the object `sim4iter` generated by the function `simulateCTStudy5years`. We can estimate the population size and trend for each iteration using the random encounter model, accounting for imperfect detectability, and focusing only on active periods:

```
pro <- processCTStudy(sim4iter, method="REM",
                      detectability=TRUE, active=TRUE)
```

Which gives:

```
pro

## [[1]]
##                  PtEst       EstSE      EstCIl       EstCIu
## N1         291.6958701 40.43220176 227.4811078 359.24429941
## N2         272.1135294 46.17264877 203.0416438 357.19917571
## N3         228.1892469 39.77055979 163.0923145 296.78601443
## N4         232.8452590 41.82574730 168.7005596 306.00228878
## N5         215.9898598 37.74538098 156.3565939 277.95416389
## lambda      -0.0728877  0.05095896  -0.1579674   0.01459537
## changeRate  -0.2595375  0.13331468  -0.5267023  -0.10304277
##
## [[2]]
##                  PtEst       EstSE      EstCIl
## N1         242.9619916 40.89002446 182.1830504
## N2         314.0330379 56.93117054 227.0092372
## N3         270.3653266 42.87063271 205.3246113
## N4         242.3454573 37.11275432 184.9478490
## N5         180.5553759 29.24477076 137.2594788
## lambda      -0.0817512  0.04414245  -0.1542824
## changeRate  -0.2568575  0.13706820  -0.4816596
##                   EstCIu
## N1         317.542096652
## N2         414.674518296
## N3         349.434431608
## N4         302.670923382
## N5         231.329897987
## lambda      -0.007888987
## changeRate  -0.031743001
##
## [[3]]
```

```
##                    PtEst       EstSE      EstCIl
## N1          336.48727129 61.12067588 240.0642228
## N2          270.75279775 49.25237322 194.7124287
## N3          230.47422765 38.90969801 169.1095726
## N4          233.61987505 47.45897886 157.8366431
## N5          265.23643941 53.46238399 184.5348914
## lambda       -0.06043551  0.04797757  -0.1359176
## changeRate   -0.21174897  0.17412984  -0.4587347
##                   EstCIu
## N1          439.67388300
## N2          352.29530702
## N3          293.33732194
## N4          311.28097793
## N5          356.93857704
## lambda        0.01752443
## changeRate    0.11370361
##
## [[4]]
##                   PtEst       EstSE      EstCIl       EstCIu
## N1          414.7676900 73.73066898 301.5556654 537.29397657
## N2          302.9244810 52.50542347 219.4260664 390.35508846
## N3          337.4753046 64.95328922 229.3841924 445.60767300
## N4          230.1374994 52.79830523 147.2879996 320.65824923
## N5          192.6507901 44.41650058 121.3770546 266.78253192
## lambda       -0.1654382  0.05688493  -0.2660750  -0.07739656
## changeRate   -0.5355212  0.18906698  -0.5940049   0.01374056
```

For each simulation, this function returns the point estimate and 90% confidence interval on the population size estimate for each year (N1 to N5), on the trend $\lambda$, and on the rate of change between the first year $N_1$ and the last year $N_5$, i.e. $(N_5 - N_1)/N_1$.

Using a suitable parameterization, the function `processCTStudy` can be used to summarize the estimates obtained for any of the 6 methods considered in our paper (see section S4.1). To make things simpler, we provide a function `process_all` implementing all these 6 methods (see next section for an example of use).

### S5 The main simulations carried out in the paper

#### S5.1 Monitoring of the roe deer during five years: no habitat selection and random sampling of traps)

##### S5.1.1 Using 100 traps

The first set of simulations was carried out to compare random encounter models and instantaneous sampling on 1000 simulations of a five-year study. These simulations can be easily carried out using the function `simulateCTStudy5years`. WARNING: THIS CALCULATION TAKES A VERY LONG TIME!!!! (see below):

```
simrd5nohs <- simulateCTStudy5years(nCT = 100, duration = 30,
                                    niter = 1000, dfPopSize,
                                    listPatches, contourChize, contourN=northChize,
                                    contourS=southChize,
                                    dff=relationAnimalSigmaMaxd, habitatMap,
                                    listptg=mvtMatrices$listMatBetween,
```

```
                        ptp=mvtMatrices$MatWithin, nofi = 0, backup = TRUE)
```

We did not include the result of these simulations in the package, as the resulting object size was too large to fit in a R package (2.5 Gb), though it is of course available from the authors upon request. Note that, in practice, we did not exactly use this R code to simulate the 1000 studies; we used this code with `niter=100` in 10 parallel sessions on a workstation, which took 4 days of calculation. We then combined our results together using the function `c.simCT` (see the help page of this function) into an object `simrd5nohs`. Then, for each simulation, we estimated, using the six methods described in section S4.1, the population size for each year, the variation rate between year 1 and year 5 and the trend $\lambda$ (together with their 90% bootstrap confidence interval). We use the function `process_all` (again, recall that this code will not work since the main object `simrd5nohs` is not available in the package. This code is provided only for information purposes):

```
roeNoHS <- process_all(simrd5nohs)
```

The reader can test this code by replacing `simrd5nohs` by the dataset `sim4iter` containing only 4 simulations (but they should be warned that even in this case, this code is rather slow). Anyway, the resulting object `roeNoHS` is available as a dataset of the package for the 1000 simulations:

```
str(roeNoHS,1)

## List of 6
##  $ REM_raw           :List of 1000
##  $ REM_detect        :List of 1000
##  $ REM_detect_active:List of 1000
##  $ IS_raw            :List of 1000
##  $ IS_detect         :List of 1000
##  $ IS_detect_active :List of 1000
```

We can then build the table summarizing the main results of these simulations. The function `present_results_CT` from the package can be used:

```
oldopt <- options(width = 100) ## to allow a cleaner display of the results

present_results_CT(roeNoHS, dfPopSize)

##          method Param TrueValue MeanEst     SE  SEEst SE.SEEst  Pcov  PDec PsigD
## 1       REM_raw    N1       300  237.31 37.875 38.687    6.215 0.511
## 2       REM_raw    N5       239 181.767 33.792 31.553    6.114 0.444
## 3       REM_raw   lam   -0.0547  -0.063  0.049  0.048    0.007 0.869 0.906 0.364
## 4       REM_raw    CR   -0.2033   -0.22  0.165  0.165    0.043 0.859 0.906 0.378
## 5
## 6       REM_det    N1       300  303.86  45.68  49.51    7.853 0.923
## 7       REM_det    N5       239 232.533 40.045 40.352    7.572 0.882
## 8       REM_det   lam   -0.0547  -0.063  0.049  0.048    0.007 0.867 0.906 0.367
## 9       REM_det    CR   -0.2033   -0.22  0.165  0.169    0.042   0.9 0.906 0.327
## 10
## 11 REM_det_act    N1       300 303.224 47.154 50.959    8.427 0.928
## 12 REM_det_act    N5       239 232.706 41.018 41.146     7.83 0.882
## 13 REM_det_act   lam   -0.0547  -0.062  0.051   0.05    0.007 0.867 0.889 0.344
## 14 REM_det_act    CR   -0.2033  -0.217  0.172  0.176    0.045 0.904 0.891 0.311
## 15
## 16       IS_raw    N1       300  54.903  9.955 10.084    1.897     0
## 17       IS_raw    N5       239  42.983  8.975    8.3    1.844     0
## 18       IS_raw   lam   -0.0547  -0.058  0.055  0.055    0.008  0.88 0.843 0.289
```

```
## 19      IS_raw    CR   -0.2033   -0.198   0.195   0.198    0.057  0.87 0.848 0.279
## 20
## 21      IS_det    N1       300 262.512 47.812 46.688   11.689 0.737
## 22      IS_det    N5       239 205.374 40.169 37.611    9.195 0.682
## 23      IS_det   lam   -0.0547   -0.057   0.056   0.054    0.008 0.884 0.854 0.279
## 24      IS_det    CR   -0.2033   -0.197   0.196   0.197    0.056 0.882 0.841 0.276
## 25
## 26  IS_det_act    N1       300 298.869 57.784 56.794   15.407 0.881
## 27  IS_det_act    N5       239  233.97 49.233 45.015   12.228 0.849
## 28  IS_det_act   lam   -0.0547   -0.057   0.059   0.057    0.009 0.885 0.833 0.272
## 29  IS_det_act    CR   -0.2033   -0.193   0.211   0.214    0.067 0.891 0.834 0.257
## 30

options(oldopt)
```

In this table, for each method and each parameter of interest (population size of year 1 `N1` and 5 `N5`, trend $\lambda$ `lam` and change rate `CR`), we present the true value of the parameter, the mean of the point estimate calculated over the 1000 simulations (`MeanEst`), their standard deviation `SE`, the mean value of standard deviation estimated by bootstrap (`SEEst`), the coverage probability of the 90% confidence interval estimated by bootstrap (probability that the true value is in the confidence interval, `Pcov`), and for the trend $\lambda$ and the rate of change, the probability that the parameter is negative (`PDec`, that is, the probability that the point estimate indicates a population decrease) and the probability that this decrease is significant (`PsigD`, that is, the probability that the upper limit of the confidence interval on the parameter is negative). These results are discussed in the paper.

##### S5.1.2 Using 25 traps

We then use the function `sampleCTs` to subsample the number of camera traps (i.e. keep only the encounter detections that occurred in the first 25 simulated traps for each simulation). Here too, this calculation will not work on the user's computer, as the dataset `simrd5nohsrd` is not available in the package. This code is provided only for information purpose:

```
simrd5nohs25 <- sampleCts(simrd5nohs, 25)
```

And we use again the function `process_all` provided in the package to calculate the population size and trends estimates for each one (this code is also provided for information purpose. However the resulting object `roeNoHS25` is available in the package as a dataset):

```
roeNoHS25 <- process_all(simrd5nohs25)
```

And finally, we use the function `present_results_CT` from the package to summarize the study with only 25 traps:

```
oldopt <- options(width = 100)

present_results_CT(roeNoHS25, dfPopSize)

##          method Param TrueValue MeanEst      SE   SEEst SE.SEEst  Pcov   PDec PsigD
## 1       REM_raw    N1       300 235.668  78.199 72.339   23.699 0.688
## 2       REM_raw    N5       239 181.565  64.232 59.381   21.176 0.638
## 3       REM_raw   lam   -0.0547   -0.057   0.099   0.097    0.026  0.87 0.732 0.186
## 4       REM_raw    CR   -0.2033   -0.161   0.383   0.417    0.358 0.871 0.738 0.193
## 5
## 6       REM_det    N1       300 301.727  93.669  92.96   29.521 0.868
```

```
## 7       REM_det    N5        239 232.242  76.207 76.196    26.018 0.864
## 8       REM_det   lam    -0.0547  -0.057   0.099  0.097     0.026 0.872 0.732 0.192
## 9       REM_det    CR    -0.2033  -0.161   0.383  0.417     0.289 0.887 0.738 0.161
## 10
## 11 REM_det_act    N1        300 301.069  97.229 95.555    31.466 0.871
## 12 REM_det_act    N5        239 231.893  78.276 77.683    26.871 0.861
## 13 REM_det_act   lam    -0.0547  -0.056   0.104    0.1     0.027 0.861 0.722 0.187
## 14 REM_det_act    CR    -0.2033   -0.15   0.413  0.436     0.276 0.877 0.734 0.159
## 15
## 16      IS_raw    N1        300  54.785  20.805 18.818     7.119     0
## 17      IS_raw    N5        239  42.953  16.803 15.415     6.117     0
## 18      IS_raw   lam    -0.0547  -0.053   0.115  0.111     0.034 0.856 0.701 0.159
## 19      IS_raw    CR    -0.2033  -0.107   0.503  0.628     0.908 0.848 0.701 0.166
## 20
## 21      IS_det    N1        300 263.562  97.551 87.342     36.81 0.768
## 22      IS_det    N5        239 204.538  75.535 69.373    28.575 0.745
## 23      IS_det   lam    -0.0547  -0.053   0.113  0.108      0.03 0.851 0.699 0.157
## 24      IS_det    CR    -0.2033  -0.119   0.482  0.605      1.06 0.856 0.703 0.167
## 25
## 26  IS_det_act    N1        300 301.313 120.668 105.91    50.082 0.816
## 27  IS_det_act    N5        239 233.349  91.489 83.106    36.841 0.826
## 28  IS_det_act   lam    -0.0547  -0.051   0.124  0.115     0.033 0.856 0.699 0.147
## 29  IS_det_act    CR    -0.2033  -0.091   0.586  0.813     3.492  0.85 0.688 0.155
## 30

options(oldopt)
```

#### S5.2 Monitoring of the roe deer during five years: selection of paths and traps preferentially placed on paths

##### S5.2.1 Simulated habitat selection process

We also examined the effect on the estimates of the common practice consisting in placing camera traps only in "good" habitats. This practice is generally used when the aim is not to estimate the population size, but rather to provide an index of its changes with time. We simulated habitat selection in the roe deer, and we placed all the 100 traps in the habitat selected by the roe deer to assess how this practice affects the trend estimates.

Roe deer is known to be a species selecting ecotones. We simplified this behaviour and simulated a selection of paths neighborhood (defined here as the area located at less than 20 m from a path) in our study area. The sf object `pathsMap` contains the map of these paths:

```
plot(pathsMap)
```

id

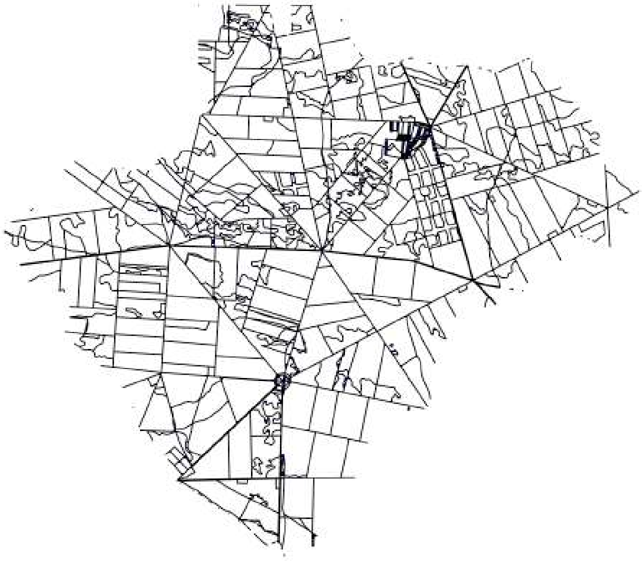

We simulated habitat selection using the following approach. For a given roe deer, we first sampled the location of a centroid for the home-range. As previously, this centroid is sampled in the northern or southern part of the study area, with respective probability 193/300 and 1-193/300 (to simulate the difference in population densities between these two parts of the study area). We selected the centroid so that: (i) it was located within 20 m from the paths (within the “paths neighborhood”), (ii) it was located at more than 600 m from the study area boundary (so that the movements of the animal around this center do not lead the animal outside the study area). Then a random number of patches (uniform between 3 and 10) are selected within a random distance from this centroid (this distance is drawn randomly from a uniform distribution bounded by 100 m and 300 m), making sure that all these patches centers are located in the areas at less than 20 m from the paths. These patches are used as center of attractions in the function `roeDeerMovement`. This habitat selection is simulated by the R function `HSlocation` from the package. We illustrate this process, also using different R objects provided in the package: `contourChize` contains the coordinates of the boundary of the study area, `pathsBuffer` contains a 20-m buffer from the paths:

```
## Chize map
chizeMap <- sf::st_sfc(sf::st_polygon(list(as.matrix(contourChize))))

## Erode this map by 600 m
erodedMap <- sf::st_buffer(chizeMap, -600)

## Intersects with areas located < 20 m from paths
sf::st_crs(pathsBuffer) <- sf::st_crs(erodedMap)
corePaths <- sf::st_intersection(st_geometry(pathsBuffer),
                                  st_geometry(erodedMap))

## Plot the result
plot(sf::st_geometry(chizeMap), col="grey")
plot(corePaths, col="red", add=TRUE)

## A random centroid will be randomly drawn from the maps in
## corePaths and patches will be drawn from pathsBuffer
set.seed(777) ## for reproducibility
(locs <- HSlocation(corePaths, pathsBuffer))
```

```
## $centroide
##              X       Y
## [1,] 388028.5 2127551
##
## $patchs
##                X          Y
##  [1,]   21.75545  185.88996
##  [2,] -189.54821 -127.32704
##  [3,]  130.06417  -69.60880
##  [4,]  155.44426  -49.24965
##  [5,]  -60.34629  190.74063
##  [6,]  -76.42551  -59.58044
##  [7,]   74.28959   17.72199
##  [8,] -109.25065  -91.37951
##  [9,]  -47.96388  182.71930
## [10,]    8.13194  -25.83990

## add these points on the map:
points(sweep(locs$patchs, 2, locs$centroide, "+"), pch=21,
       bg="yellow")
points(locs$centroide[1], locs$centroide[2], pch=3, cex=2, lwd=4, col="blue")
points(locs$centroide[1], locs$centroide[2], pch=3, cex=2, lwd=2, col="green")
```

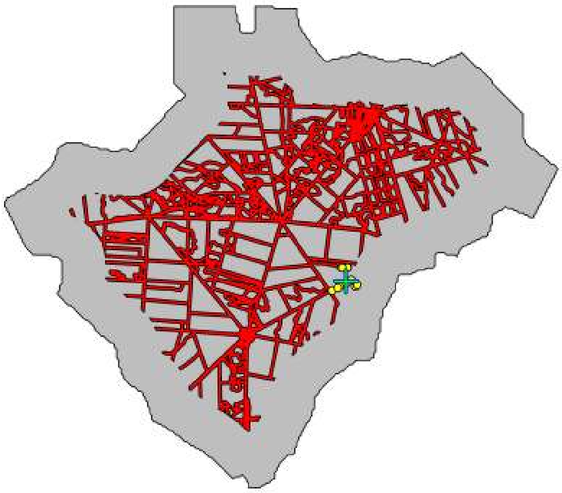

The grey area corresponds to the Chize Study Area. The red areas correspond to the areas located at a distance greater than 600 m from the boundary of this study area and at less than 20 m from a path. The green cross corresponds to the simulated centroid for the roe deer and the yellow points correspond to the simulated centres of attraction for the roe deer movements.

We can then use the function `roeDeerMovement` as before to simulate the movements of the animals based on these attraction points. As all attraction points are located in the paths neighborhood, this habitat type will appear as selected by the animals.

##### S5.2.2 Simulation

The function `simulateCTStudy5years` can be used to simulate a camera trap monitoring of the roe deer population, with a habitat selection by the animals (this is controlled by the parameter `habitatSelection = TRUE`). We then placed all the 100 traps on the paths for the monitoring (this is controlled by the parameter `propPaths = 1`). We used the following code. WARNING: THIS CALCULATION TAKES A VERY LONG TIME!!!! To limit the computation time, we only simulated 500 times this study:

```
simrd5withhs <- simulateCTStudy5years(nCT = 100, duration = 30,
                                      niter = 500, dfPopSize,
                                      listPatches, contourChize, contourN=northChize,
                                      contourS=southChize,
                                      dff=relationAnimalSigmaMaxd, habitatMap,
                                      listptg=mvtMatrices$listMatBetween,
                                      ptp=mvtMatrices$MatWithin, stratSampling=TRUE,
                                      propPaths=1, pathsMap=pathsMap,
                                      pathsBuffer=pathsBuffer, offPathsMap=offPathsMap,
                                      offPathsBuffer=offPathsBuffer,
                                      habitatSelection=TRUE,
                                      nofi = 0, backup = TRUE)
```

As before, we did not include the result of these simulations in the package, as the resulting object size was too large to fit in a R package (1.7 Gb), though it is of course available from the authors upon request. Then, for each simulation, we estimated, using each possible assessed method, the trend $\lambda$ (together its 90% bootstrap confidence interval). The 6 assessed methods were the six methods described in section S4.1. We used the code below. Note that this code will not work on the readers' computer, as the main object `simrd5withhs` is not available in the package. This code is provided for information purposes. Note also that the resulting list `roeHSBiasedSam` is available as a dataset in the package:

```
## we use the function compute_all written above:
roeHSBiasedSam <- process_all(simrdwithhs)
```

We can then format the results:

```
trend1 <- exp(coef(lm(log(dfPopSize[,1])~c(1:5)))[2])-1
nam <- c("REM_raw","REM_det","REM_det_act",
         "IS_raw","IS_det","IS_det_act")
sel <- 6

oldopt <- options(width = 100)

do.call(rbind,lapply(1:length(roeHSBiasedSam), function(i) {
    x <- roeHSBiasedSam[[i]]
    method <- nam[i]
    MeanEst <- mean(sapply(x, function(y) y[sel,1]), na.rm=TRUE)
    SE <- sd(sapply(x, function(y) y[sel,1]), na.rm=TRUE)
    SEEst <- mean(sapply(x, function(y) y[sel,2]), na.rm=TRUE)
    SE.SEEst <- sd(sapply(x, function(y) y[sel,2]), na.rm=TRUE)
    pcover <- mean(sapply(x, function(y)
        trend1>=y[sel,3]&trend1<=y[sel,4]))

    Psigla <- mean(sapply(x, function(y) y[6,4]<0))
    PDecla <- mean(sapply(x, function(y) y[6,1]<0))

    data.frame(method=method,
               MeanEst=round(MeanEst,3),
               SE=round(SE,3),
```

```
              SEEst=round(SEEst,3), SE.SEEst=round(SE.SEEst,3), Pcov=round(pcover,3),
              PDec=round(PDecla,3),
              PsigD=round(Psigla,3))
}))

##        method MeanEst    SE SEEst SE.SEEst  Pcov  PDec PsigD
## 1     REM_raw  -0.084 0.030 0.029    0.004 0.732 0.996 0.870
## 2     REM_det  -0.084 0.030 0.029    0.004 0.728 0.996 0.862
## 3 REM_det_act  -0.084 0.031 0.031    0.004 0.750 0.996 0.830
## 4      IS_raw  -0.081 0.036 0.035    0.005 0.770 0.994 0.696
## 5      IS_det  -0.083 0.035 0.035    0.005 0.788 0.988 0.714
## 6  IS_det_act  -0.083 0.037 0.038    0.006 0.814 0.986 0.692

options(oldopt)
```

These results are also discussed in the paper.

##### S5.2.3 Proportion of the time spent near the paths

In these simulations, we supposed that the mean home-range size of animals increased with time. Therefore, the dispersion of the movements around attraction points was larger in year 5 than in year 1. As attraction points were located in the neighborhood of paths (within 20 m from the paths), this implies that the animals spent less time in this habitat type in year 5 than in year 1. Even if the number of animals on the area had been constant during the five-year period, the mean number of animals present within 20 m from the paths at any moment would have decreased with time.

We illustrate this idea using simulations: we simulated the home range of 1000 animals selecting the neighborhood of paths during the 5 years (while increasing their home-range size in the same way as in our previous simulations), to assess how the proportion of the time that the animals spent in this selected habitat changed with time. WARNING: this calculation is long! The resulting object `propTimePathsHS` is available as a dataset in the package (see below):

```
## Number of simulated animals
ntotani <- 1000

## Required for the simulation: the centroid of animals is located at
## at least 600 m from the boundary of the study area (to avoid
## movement outside the study area).
cartec <- sf::st_sfc(sf::st_polygon(list(as.matrix(contourChize))))
sf::st_crs(cartec) <- sf::st_crs(pathsBuffer)
cartec <- sf::st_buffer(cartec, -600)

## We then intersected the map of the paths neighborhood with this
## "eroded map"
corePaths <- sf::st_intersection(pathsBuffer, cartec)
po1 <- sf::st_sfc(sf::st_polygon(list(as.matrix(northChize))))
sf::st_crs(po1) <- sf::st_crs(corePaths)

## Because 193/300 animals were located in the northern part, we
## intersected this paths map with the limits of the north an south
## of Chize
northPaths <- sf::st_intersection(corePaths, po1)
po1 <- sf::st_sfc(sf::st_polygon(list(as.matrix(southChize))))
sf::st_crs(po1) <- sf::st_crs(corePaths)
```

```r
southPaths <- sf::st_intersection(corePaths, po1)

## Parameters of movement to simulate the increase in home range size
## (through the use of sigma) and the habitat selection (through
## HSlocation)
paramani <- lapply(1:ntotani, function(i) {
    cat(i,"\r")
    north <- (runif(1) < 193/300)
    minsigan <- round(4 * (1:5)/5)
    maxsigan <- round(3 + 7 * (0:4)/4)
    sigmaan <- sapply(1:5, function(t) sample(minsigan[t]:maxsigan[t],
                                              1))
    zone <- southPaths
    if (north)
        zone <- northPaths
    lxyz <- HSlocation(zone, pathsBuffer)
    return(list(lxyz = lxyz, sigmaan = sigmaan))
})

## Simulation of the movements of the 1000 animals +
## calculation of the proportion of relocations in paths neighborhood.
propTimePathsHS <- matrix(0, nrow=1000, ncol=5)
for (iAni in 1:ntotani) {
    cat(iAni,"\r")
    for (year in 1:5) {

        ## Simulate the movement
        paAni <- paramani[[iAni]]
        lipt <- list(ptg=mvtMatrices$listMatBetween[[paAni$sigmaan[year]]],
                     ptp=mvtMatrices$MatWithin)
        lixyt <- list(paAni$lxyz$patchs, paAni$lxyz$patchs)
        z <- roeDeerMovement(30, lixyt, lipt = lipt,
                             verbose = FALSE)
        z <- t(t(z)+as.vector(paAni$lxyz$centroide))

        ## To speed up the calculation, consider one reloc every 1000
        ## seconds
        z <- z[seq(1,nrow(z), by=1000),]

        ## Join with the map of paths Buffer, and calculate the proportion of
        ## relocations that are in this habitat type
        z <- as.data.frame(z)
        jo <- st_join(st_as_sf(z,coords=c("x","y"), crs=st_crs(pathsBuffer)),
                      pathsBuffer)
        propTimePathsHS[iAni, year] <- mean(!is.na(jo$id))
    }
}
```

Using this object `propTimePathsHS`, we can calculate the mean percentage of time spent by the animals in the neighborhood of the paths (+/- SE):

```r
moa <- 100*apply(propTimePathsHS,2,mean)
soa <- 100*apply(propTimePathsHS,2,mean)/sqrt(1000)

plot(1:5, moa, ylim=range(c(moa+soa, moa-soa)),
     xlab="Year",ylab="Percentage of time in paths neighborhood",
     ty="b")
```

```
arrows(1:5, moa-soa, 1:5, moa+soa, angle=90,
       length=0.05, code=3)
```

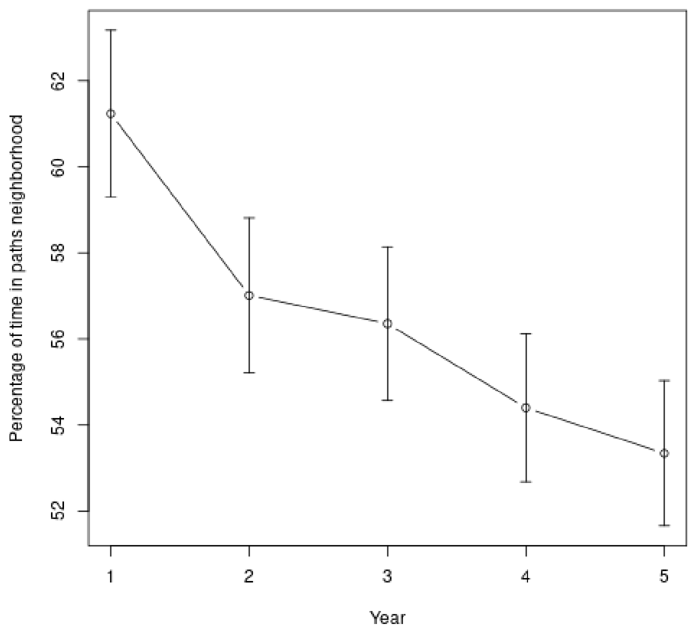

For each simulated animal $r$, we calculated the proportion $p_t^{(r)}$ of the time spent close to a path during year $t$. The mean proportion spent by the 1000 animals in this habitat type is classically estimated with:

$$\overline{p_t} = \frac{1}{1000} \sum_{r=1}^{1000} p_t^{(r)}$$

With classically, a standard error equal to:

$$\mathrm{SE}(\overline{p_t}) = \sqrt{\frac{\mathrm{Var}(p_t^{(r)})}{1000}}$$

Moreover, in our study simulating a population of $N_t$ animals over five years, there are in average $\hat{H}_t = N_t \times \overline{p_t}$ animals present in this habitat type at a given time of the year $t$. We can use these simulations of 1000 animals during 5 years to estimate the rate of decrease $\mu'$ with time of the number of animals present in this habitat type using a linear regression:

$$\log \hat{H}_t = a + (\log \mu') \times t + e_t \tag{2}$$

with $e_t$ a residual with zero mean. Note that the variance of $\log(\hat{H}_t)$ – and therefore of $e_t$ – is not constant (see previous figure). We can account for this heteroskedasticity by weighing the linear regression by:

$$w_t = 1/\mathrm{Var}(\log \hat{H}_t)$$

Note that, $\mathrm{Var}(\log \hat{H}_t) = \mathrm{Var}(\log N_t + \log \overline{p_t}) = \mathrm{Var}(\log \overline{p_t})$ (because $\mathrm{Var}(\log N_t) = 0$ since $N_t$ is known). Using the delta-method, we can show that:

$$\mathrm{Var}(\log \overline{p_t}) \approx \frac{\mathrm{Var}(\overline{p_t})}{\overline{p_t}^2}$$

And $\mathrm{Var}(\overline{p_t})$ can be estimated from the data simulated above. We can therefore estimate $\lambda' = 1 - \exp \mu'$ expected for the number of animals in paths neighborhood using equation 2, weighting each year with $1/\mathrm{Var}(\log \overline{p_t})$:

```r
## moa and soa as a proportion
moap <- moa/100
soap <- soa/100

## Real number in each habitat type
ht <- c(300,274,266,246,239)*moap
## variance of \bar{ht}
voa <- soap^2

## variance of log(ht) with the delta-method
vh <- voa/(moap^2)

## Weighted regression
su <- summary(lm(log(ht)~c(1:5), weight=1/vh))
cothb <- exp(round(su$coefficients[2,1],2))-1
## standard error
scothb <- round(sqrt(((su$coefficients[2,2])^2)*
                      exp(2*exp(round(su$coefficients[2,1],2)))),2)

## slope:
cothb

## [1] -0.08606881

## standard error
scothb

## [1] 0.03
```

With this regression, we estimate $\lambda$ = -0.086 (SE = 0.03), which corresponds roughly to the value we estimated by placing all the traps in paths neighborhood.

#### S5.3 Habitat selection and stratification

##### S5.3.1 Implementing stratification in the simulated monitoring

In this section, we also simulated a roe deer population during five years where animals preferentially use the neighborhood of paths, but we tried to use a stratified sampling of the traps and a corresponding estimator to obtain an unbiased estimation. We placed a proportion $p$ of the traps in the selected habitat and monitored the population for five years. Then, we estimated the population size for each year (by estimating the population size in each habitat – close to paths and far from paths – separately and by summing the two estimates). We simulated this study for the following values of $p$: 0.25, 0.5, 0.75. We used the following code. WARNING: THIS CALCULATION TAKES A VERY LONG TIME!!!! Since this calculation was very long, we only simulated 500 times this study for each value of $p$:

```r
listSimulated <- list()
propsim <- c(0.25,0.5,0.75)

for (i in 1:3) {
    listSimulated[[i]] <- simulateCTStudy5years(nCT = 100, duration = 30,
                                                niter = 500, dfPopSize,
                                                listPatches, contourChize,
                                                contourN=northChize,
                                                contourS=southChize,
```

```
                                                  dff=relationAnimalSigmaMaxd,
                                                  habitatMap=habitatMap,
                                                  listptg=mvtMatrices$listMatBetween,
                                                  ptp=mvtMatrices$MatWithin,
                                                  stratSampling=TRUE,
                                                  propPaths=propsim[i],
                                                  pathsMap=pathsMap,
                                                  pathsBuffer=pathsBuffer,
                                                  offPathsMap=offPathsMap,
                                                  offPathsBuffer=offPathsBuffer,
                                                  habitatSelection=TRUE,
                                                  nofi = 0, backup = TRUE)
}
```

As before, we did not include the result of these simulations in the package, as the resulting object size was too large to fit in a R package (6.6 Gb), though it is available from the authors upon request. Then, for each simulation, we estimated, using each possible assessed method, the trend $\lambda$ (together its 90% bootstrap confidence interval). The 6 assessed methods were the six methods described in section S4.1. We used the code below. Note that this code will not work on the readers' computer, as the main object `listSimulated` is not available in the package. This code is provided for information purposes. Note also that the resulting list `roeStratSam` is available as a dataset in the package:

```
roeStratSam <- lapply(listSimulated, function(x) {
    process_all(x,stratified=TRUE, pathsBuffer=pathsBuffer)
})
```

Consider the population size the first year (in theory equal to 300). We can consider how the six methods estimate this parameter, for each proportion of traps in the selected habitat type:

```
oldopt <- options(width = 100)
do.call(rbind,
        lapply(1:3,
               function(i) {

            df1 <- do.call(rbind,
                           lapply(1:6,
                                  function(j) {
                               data.frame(
                                   p = c(0.25, 0.5, 0.75)[i],
                                   Esp.Dech=round(mean(sapply(roeStratSam[[i]][[j]],
                                                              function(x) x[1,1])),1),
                                   SE.Dech=round(sd(sapply(roeStratSam[[i]][[j]],
                                                           function(x) x[1,1])),1),
                                   Couv.IC=round(mean(sapply(roeStratSam[[i]][[j]],
                                                             function(x)
                                   (x[1,3]<300)&(x[1,4]>300))),3),
                                   Estim.SE=round(mean(sapply(roeStratSam[[i]][[j]],
                                                              function(x) x[1,2]),
                                                       na.rm=TRUE),1),
                                   SE.Estim.SE=round(sd(sapply(roeStratSam[[i]][[j]],
                                                               function(x) x[1,2]),
                                                        na.rm=TRUE),1)
                               )
                           }))
            df1$Method <- c("REM_raw","REM_det","REM_det_act",
                            "IS_raw","IS_det","IS_det_act")
```

```
            df1 <- df1[,c(1,7,2:6)]
            if (i==3)
                return(df1)
            return(rbind(df1,
                         data.frame(p="",Method="",Esp.Dech="",SE.Dech="",
                                    Couv.IC="",Estim.SE="",SE.Estim.SE="")))

        }))
```

```
##       p      Method Esp.Dech SE.Dech Couv.IC Estim.SE SE.Estim.SE
## 1  0.25     REM_raw    235.9    53.3   0.574     49.6        11.2
## 2  0.25     REM_det    300.1    62.7   0.868       58        12.1
## 3  0.25 REM_det_act      299    63.8   0.862     59.1        12.5
## 4  0.25      IS_raw     54.7    13.9       0     12.7         3.3
## 5  0.25      IS_det    263.7    69.3    0.73     59.4        23.8
## 6  0.25  IS_det_act    300.7    85.7    0.84     71.5        35.1
## 7
## 8   0.5     REM_raw    235.6      45   0.526     42.6         7.4
## 9   0.5     REM_det    298.7    51.7   0.888     49.7         8.2
## 10  0.5 REM_det_act    298.9    52.9    0.88       51         8.4
## 11  0.5      IS_raw     54.3    11.6       0     10.9         2.2
## 12  0.5      IS_det    259.5    53.3   0.712     49.8        11.8
## 13  0.5  IS_det_act      298    64.3   0.868     60.5        17.7
## 14
## 15 0.75     REM_raw    236.7    47.1   0.598     46.9        13.2
## 16 0.75     REM_det      299    54.7   0.876     56.9        14.4
## 17 0.75 REM_det_act    299.4    57.4   0.876     58.6        15.8
## 18 0.75      IS_raw     54.8      12       0     11.8         3.7
## 19 0.75      IS_det      262      58    0.73     54.8        19.8
## 20 0.75  IS_det_act    298.9    71.8   0.862     66.3        27.8
```

```
options(oldopt)
```

Note that the precision of the population size estimates is not strongly affected by the the level of allocation in the stratified sample. The estimated precision for stratified sample was comparable to the precision estimated without stratification, though slightly better.

We can similarly estimate the trend $\lambda$ for each method and each proportion of traps in paths neighborhood:

```
oldopt <- options(width = 100)
do.call(rbind,
        lapply(1:3,
               function(i) {

            df1 <- do.call(rbind,
                           lapply(1:6,
                                  function(j) {
                               data.frame(
                                   p = c(0.25, 0.5, 0.75)[i],
                                   Esp.Dech=round(mean(sapply(roeStratSam[[i]][[j]],
                                                              function(x) x[6,1])),3),
                                   SE.Dech=round(sd(sapply(roeStratSam[[i]][[j]],
                                                           function(x) x[6,1])),4),
                                   Couv.IC=round(mean(sapply(roeStratSam[[i]][[j]],
                                                             function(x)
                                   (x[6,3]< -0.054)&(x[6,4]>-0.054))),3),
```

```
                                    Estim.SE=round(mean(sapply(roeStratSam[[i]][[j]],
                                                           function(x) x[6,2]),
                                                    na.rm=TRUE),3),
                                    SE.Estim.SE=round(sd(sapply(roeStratSam[[i]][[j]],
                                                            function(x) x[6,2]),
                                                     na.rm=TRUE),3)
                                )
                            }))
            df1$Method <- c("REM_raw","REM_det","REM_det_act",
                            "IS_raw","IS_det","IS_det_act")
            df1 <- df1[,c(1,7,2:6)]

            if (i==3)
                return(df1)
            return(rbind(df1,
                         data.frame(p="",Method="", Esp.Dech="",SE.Dech="",
                                    Couv.IC="",Estim.SE="",SE.Estim.SE="")))
        }))

##        p      Method Esp.Dech SE.Dech Couv.IC Estim.SE SE.Estim.SE
## 1  0.25     REM_raw   -0.057  0.0392   0.894     0.04       0.009
## 2  0.25     REM_det   -0.052  0.0378     0.9    0.038       0.007
## 3  0.25 REM_det_act   -0.051  0.0395   0.904     0.04       0.008
## 4  0.25      IS_raw   -0.048  0.0483   0.884    0.048        0.01
## 5  0.25      IS_det   -0.051  0.0506    0.89    0.048       0.011
## 6  0.25  IS_det_act   -0.051  0.0551   0.868    0.052       0.011
## 7
## 8   0.5     REM_raw   -0.058  0.0355    0.89    0.035       0.006
## 9   0.5     REM_det   -0.053  0.0332   0.892    0.033       0.005
## 10  0.5 REM_det_act   -0.053  0.0348   0.892    0.035       0.005
## 11  0.5      IS_raw   -0.052   0.043   0.878    0.042       0.007
## 12  0.5      IS_det   -0.053  0.0437   0.886    0.043       0.008
## 13  0.5  IS_det_act   -0.055  0.0484   0.894    0.047       0.009
## 14
## 15 0.75     REM_raw   -0.057  0.0413   0.856    0.039       0.009
## 16 0.75     REM_det   -0.052  0.0401   0.844    0.037       0.008
## 17 0.75 REM_det_act   -0.052  0.0427   0.856    0.038       0.008
## 18 0.75      IS_raw   -0.051   0.049    0.86    0.046        0.01
## 19 0.75      IS_det   -0.052  0.0513   0.862    0.047       0.011
## 20 0.75  IS_det_act   -0.052  0.0551   0.866    0.051       0.012

options(oldopt)
```

We can see that placing half of the traps in the paths neighborhood leads to a slightly more precise estimate, but the difference with other allocations was not very strong, suggesting that this factor may not have a large effect on the estimation. Comparing the standard error of the estimates of the trend in the stratified case ($\approx 0.035$ to 0.055) with the standard errors of the trend in the simulations carried out previously without habitat selection ($\approx 0.05$ to 0.06) suggests that stratification improves only slightly the estimate precision.

##### S5.3.2 Simulating the stratification itself

The problem with the approach described in the previous section is that it is very long to implement (it took several days of calculation just to simulate the three levels of allocation). An easier and quicker approach consists in combining the detections by traps in simulations where all traps are placed in the neighborhood of paths with the detections by traps in other simulations where all traps are not placed in this neighborhood. For example, to simulate a stratification where 10% of the traps are placed in the neighborhood of paths, we can extract the detections from the 10 first traps in the simulations carried out in section S5.2.2, and extracting the detections from the 90 first traps in simulations (that we carry out below) of a monitoring in which all traps are located far from the paths. The limit of this approach is that the animals that are simulated in the first set of simulations and in the second set are not the same. However, we will see below that the results obtained for the three levels of allocation (p = 0.25, p = 0.5, p = 0.75) tested in the previous section are the same with this approach and with the approach implementing a stratification directly within the simulated study.

Therefore, to implement this approach, we first need to simulate 500 studies where all the camera traps are located far from the paths. We use the same code as in section S5.2.2, just changing the parameter `propPaths=0`. We used the following code. WARNING: THIS CALCULATION TAKES A VERY LONG TIME!!!! To limit the computation time, we only simulated 500 times this study:

```
simrd5withhs0 <- simulateCTStudy5years(nCT = 100, duration = 30,
                                       niter = 500, dfPopSize,
                                       listPatches, contourChize,
                                       contourN=northChize,
                                       contourS=southChize,
                                       dff=relationAnimalSigmaMaxd, habitatMap,
                                       listptg=mvtMatrices$listMatBetween,
                                       ptp=mvtMatrices$MatWithin,
                                       stratSampling=TRUE,
                                       propPaths=0, pathsMap=pathsMap,
                                       pathsBuffer=pathsBuffer,
                                       offPathsMap=offPathsMap,
                                       offPathsBuffer=offPathsBuffer,
                                       habitatSelection=TRUE,
                                       nofi = 0, backup = TRUE)
```

As before, we did not include the result of these simulations in the package, as the resulting object size was too large to fit in a R package (941.2 Mb), though it is of course available from the authors upon request.

Now, the idea is to combine the detections from a given proportion of traps (e.g., 10%) from the simulations in `simrd5withhs` from the section S5.2.2 with the detections from the complementary proportion (e.g., 90%) from this object `simrd5withhs0`. We estimate the animal abundance in the paths neighborhood with the 6 methods using only the detections from the traps in the paths neighborhood; we then estimate the animal abundance in the habitat distant from paths with the six methods using only the detections from the traps in the habitat distant from paths. The abundance estimate on the whole study area is the sum of the abundance estimates in these two habitat types.

We carry out these calculations. First, to speed up the calculations, we first begin by calculating the mean detection probability for each trap in `simrd5withhs` and `simrd5withhs0` (this code is slow, and the objects `simrd5withhs` and `simrd5withhs0` are too large to fit in the package, and the result are not included in the package; this code is provided for information purposes):

```
prop0 <- lapply(1:500, function(j) {
    cat(j,"\r")
    habitat4sim <- simrd5withhs0$encounters[[j]]$indicetraps$hab
    set.seed(simrd5withhs0$encounters[[j]]$creationCT[2])
    list100CT <- lapply(1:100, function(k) {
        organiseCT(habitat4sim[k])
    })
    sapply(1:length(list100CT),
           function(trap)
```

```
        mean(1 -
             exp(-list100CT[[trap]]$vu)))
})

prop1 <- lapply(1:500, function(j) {
    cat(j,"\r")
    habitat4sim <- simrd5withhhs$encounters[[j]]$indicetraps$hab
    set.seed(simrd5withhhs$encounters[[j]]$creationCT[2])
    list100CT <- lapply(1:100, function(k) {
        organiseCT(habitat4sim[k])
    })
    sapply(1:length(list100CT),
           function(trap)
        mean(1 -
             exp(-list100CT[[trap]]$vu)))
})
```

Then we consider a variable proportion of traps in the paths neighborhood (from 5 to 95%). For each proportion, we use the function `sampleCTs` to select the corresponding proportion of traps in `simrd5withhhs` and `simrd5withhhs0`. We then use the functions `Remhab` and `IShab` to estimate abundance in the two habitat types with the six methods (close and far from paths). This code is also very slow, and cannot be executed by the user since they do not have the objects `simrd5withhhs` and `simrd5withhhs0`. However, the resulting object is available as a dataset from the package:

```
liche <- list()
lipas <- list()
litot <- list()

for (i in 1:length(pro)) {

    cat("##############################\n## Iteration",i, "\n\n")
    cat("## rem_raw\n")
    reb <- sampleCTs(simrd5withhhs, pro[i])
    rec <- sampleCTs(simrd5withhhs0, 100-pro[i])

    ## REM_raw:
    rem_raw_che <- Remhab(reb, paths=TRUE, detection=FALSE, active=FALSE)
    rem_raw_pas <- Remhab(rec, paths=FALSE, detection=FALSE, active=FALSE)
    rem_raw <- rem_raw_che+rem_raw_pas

    ## REM_detect:
    cat("## rem_det\n")
    rem_det_che <- Remhab(reb, paths=TRUE, detection=TRUE, active=FALSE)
    rem_det_pas <- Remhab(rec, paths=FALSE, detection=TRUE, active=FALSE)
    rem_det <- rem_det_che+rem_det_pas

    ## REM_detect_activ
    cat("## rem_det_act\n")
    rem_det_act_che <- Remhab(reb, paths=TRUE, detection=TRUE, active=TRUE)
    rem_det_act_pas <- Remhab(rec, paths=FALSE, detection=TRUE, active=TRUE)
    rem_det_act <- rem_det_act_che+rem_det_act_pas

    ## IS_raw:
    cat("## is_raw\n")
    is_raw_che <- IShab(reb, paths=TRUE, detection=FALSE, active=FALSE, prop=prop1)
    is_raw_pas <- IShab(rec, paths=FALSE, detection=FALSE, active=FALSE, prop=prop0)
```

```
    is_raw <- is_raw_che+is_raw_pas

    ## IS_detect:
    cat("## is_det\n")
    is_det_che <- IShab(reb, paths=TRUE, detection=TRUE, active=FALSE, prop=prop1)
    is_det_pas <- IShab(rec, paths=FALSE, detection=TRUE, active=FALSE, prop=prop0)
    is_det <- is_det_che+is_det_pas

    ## IS_detect_activ
    cat("## is_det_act\n")
    is_det_act_che <- IShab(reb, paths=TRUE, detection=TRUE, active=TRUE, prop=prop1)
    is_det_act_pas <- IShab(rec, paths=FALSE, detection=TRUE, active=TRUE, prop=prop0)
    is_det_act <- is_det_act_che+is_det_act_pas

    ##
    liche[[i]] <- data.frame(rem_raw=rem_raw_che, rem_det=rem_det_che,
                             rem_det_act=rem_det_act_che,
                             is_raw=is_raw_che, is_det=is_det_che,
                             is_det_act=is_det_act_che)
    lipas[[i]] <- data.frame(rem_raw=rem_raw_pas, rem_det=rem_det_pas,
                             rem_det_act=rem_det_act_pas,
                             is_raw=is_raw_pas, is_det=is_det_pas,
                             is_det_act=is_det_act_pas)

    litot[[i]] <- data.frame(rem_raw=rem_raw, rem_det=rem_det,
                             rem_det_act=rem_det_act,
                             is_raw=is_raw, is_det=is_det,
                             is_det_act=is_det_act)
}

stratData <- list(inPaths=liche, offPaths=lipas, all=litot)
```

We can then estimate the coefficient of variation, for each method and each proportion of traps in the paths neighborhood:

```
par(mfrow=c(3,2))
un <- unique(sapply(strsplit(names(stratData$all[[1]]),"\\."),function(x) x[1]))
pro <- seq(5,95,by=5)
tmp <- lapply(1:6, function(j) {
    cv <- sapply(1:19, function(i) {
        o <- stratData$all[[i]][,paste0(un[j],".1")]
        sd(o,na.rm=TRUE)/mean(o,na.rm=TRUE)
        })
    plot(pro/100,cv, ty="b",
         xlab="Proportion of traps close to paths",
        ylab="Coefficient of variation",
        main=un[j], ylim=c(0.15,0.5))
    abline(h=seq(0.15,0.5, by=0.1),
           v=seq(0.05,0.95,by=0.1), col="grey")
    points(pro/100, cv, ty="b")
    })
```

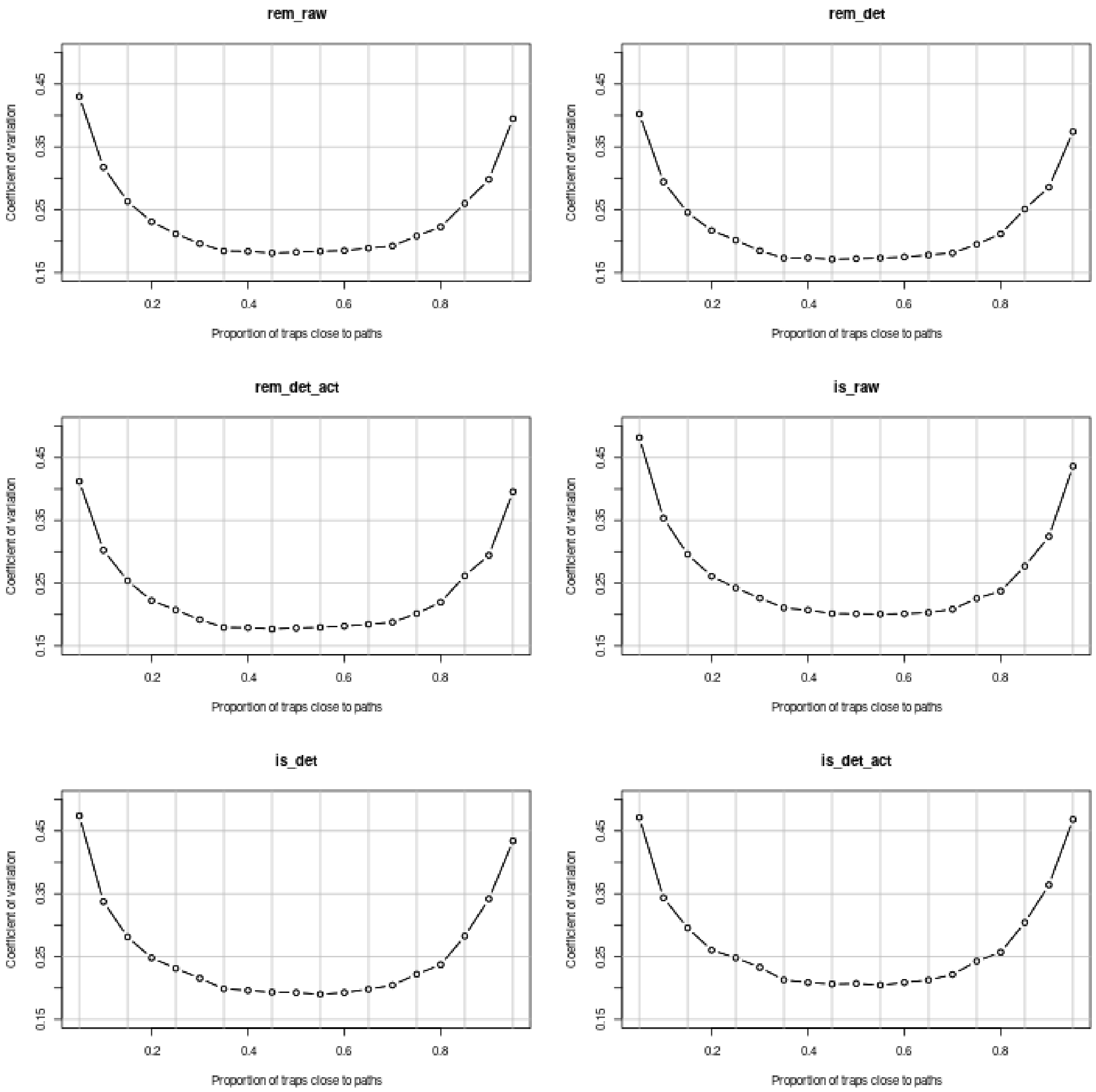

These results are interpreted in the paper. Note how the results are similar to the results of the previous section, where the stratification was implemented directly in the simulation of a five-year study.

#### S5.4 Habitat selection and random placement of camera traps

##### S5.4.1 The simulations

Finally, we conducted simulations under a habitat selection scenario while maintaining random camera-trap placement. Unlike the scenarios in section S5.2 – which combined habitat selection with non-random camera trap placement – this approach aims to isolate the specific influence of habitat selection on the estimators. By comparing these results with the baseline results in section S5.1 (random placement of camera traps without habitat selection), we can quantify the bias or imprecision introduced solely by the animals' habitat preferences, independently of any observer-driven sampling bias.

The function `simulateCTStudy5years` can also be used to simulate a camera trap monitoring of the roe deer population, with a habitat selection by the animals (this is controlled by the parameter `habitatSelection = TRUE`) and a random placement of the traps (we just let the option `stratSampling` set to the default value `FALSE`). We used the following code. WARNING: THIS CALCULATION TAKES A VERY LONG

TIME!!!! To limit the computation time, we only simulated 500 times this study:

```
roeDeerHSRandomSam <- simulateCTStudy5years(nCT = 100, duration = 30,
                                            niter = 500, dfPopSize,
                                            listPatches, contourChize,
                                            contourN=northChize,
                                            contourS=southChize,
                                            dff=relationAnimalSigmaMaxd, habitatMap,
                                            listptg=mvtMatrices$listMatBetween,
                                            ptp=mvtMatrices$MatWithin,
                                            pathsMap=pathsMap,
                                            pathsBuffer=pathsBuffer,
                                            offPathsMap=offPathsMap,
                                            offPathsBuffer=offPathsBuffer,
                                            habitatSelection=TRUE,
                                            nofi = 0, backup = TRUE)
```

Again, we did not include the result of these simulations in the package, as the resulting object size was too large to fit in a R package (123 Mb), though it is of course available from the authors upon request. We then used the function `process_all` of the package to estimate population size, trend and population decrease from these data. Note that this code will not work on the readers' computer, as the main object `roeDeerHSRandomSam` is not available in the package. This code is provided for information purposes. Note also that the resulting list `HSRSroe` is available as a dataset in the package:

```
HSRSroe <- process_all(roeDeerHSRandomSam)
```

We then present the results:

```
oldopt <- options(width = 100)
present_results_CT(HSRSroe, dfPopSize)

##         method Param TrueValue MeanEst     SE  SEEst SE.SEEst  Pcov  PDec PsigD
## 1      REM_raw    N1       300 235.748 46.758 45.599    7.993  0.59            
## 2      REM_raw    N5       239 182.399 30.646 30.547    4.731 0.436            
## 3      REM_raw   lam   -0.0547   -0.06  0.036  0.036    0.006 0.894 0.946 0.486
## 4      REM_raw    CR   -0.2033  -0.212  0.128  0.129    0.032 0.898 0.934  0.47
## 5                                                                              
## 6      REM_det    N1       300 303.782 56.987 58.858    9.775 0.908            
## 7      REM_det    N5       239 235.052  36.39 39.397    6.007 0.906            
## 8      REM_det   lam   -0.0547   -0.06  0.036  0.036    0.006 0.892 0.946 0.484
## 9      REM_det    CR   -0.2033  -0.212  0.128  0.132    0.031 0.912 0.934 0.418
## 10                                                                             
## 11 REM_det_act    N1       300 304.033 59.286 60.207   10.521 0.898            
## 12 REM_det_act    N5       239 234.925 36.631 40.039    6.117 0.906            
## 13 REM_det_act   lam   -0.0547  -0.059  0.038  0.038    0.006 0.878  0.94 0.456
## 14 REM_det_act    CR   -0.2033   -0.21   0.14  0.141    0.035 0.912  0.92 0.376
## 15                                                                             
## 16      IS_raw    N1       300  54.788  12.26 11.732    2.394     0            
## 17      IS_raw    N5       239  43.093  7.933  7.849    1.305     0            
## 18      IS_raw   lam   -0.0547  -0.055  0.042  0.043    0.007 0.894 0.902 0.336
## 19      IS_raw    CR   -0.2033  -0.192  0.158  0.164    0.044 0.894 0.876 0.312
## 20                                                                             
## 21      IS_det    N1       300 262.221 55.394 53.793   12.328 0.752            
## 22      IS_det    N5       239  208.39 37.427 36.195    8.196  0.75            
## 23      IS_det   lam   -0.0547  -0.053  0.043  0.045    0.008 0.882 0.892 0.292
## 24      IS_det    CR   -0.2033  -0.184  0.165  0.172    0.052 0.888 0.868 0.298
```

```
## 25
## 26  IS_det_act    N1       300 301.362 68.144 64.859   16.163 0.876
## 27  IS_det_act    N5       239 237.825 45.158 43.674   12.134  0.87
## 28  IS_det_act   lam   -0.0547  -0.053  0.048  0.049    0.009 0.894 0.864 0.272
## 29  IS_det_act    CR   -0.2033  -0.185  0.184  0.191    0.065 0.894 0.848 0.288
## 30

options(oldopt)
```

A comparison with Section S5.1 reveals that estimation outcomes are very similar between scenarios with and without habitat selection, provided that the camera traps are randomly placed. In terms of bias, the outcomes are virtually identical: both REM and IS provide unbiased population size estimates – provided detectability and activity rhythms are accounted for – as do the resulting change rates and trends. However the introduction of habitat selection leads to two minor differences regarding the precision of the estimates:

- The population size estimates are slightly less precise in the presence of habitat selection.
- Conversely, the trend estimation is slightly more precise with habitat selection.

While the impact of habitat selection on precision remains marginal, we believe that exploring the underlying theoretical drivers of these results is instructive for a deeper understanding of the factors factors affecting camera trap studies. Given the secondary nature of these effects relative to our primary findings, we have focused the main text on the most impactful results and provided a more detailed theoretical examination only in the following two subsections.

##### S5.4.2 A larger variance of the population size estimation with habitat selection

The increased variance of population size estimate is expected, as habitat selection leads to a greater spatial heterogeneity in animal density across the study area. This, in turn, leads to a higher variability in encounter rates among trap locations, which naturally propagates into a higher variance of the estimates.

To provide a more formal explanation of this phenomenon, consider a single year (e.g., Year 1). Let $N = 300$ be the number of animals on the study area of total surface $A = 2598$ ha. We divide the area into two habitat types: $A_1 = 986$ ha of selected habitat (path neighborhoods) and $A_2 = 1612$ ha of non-selected habitat, such that $A_1 \approx 0.38 \times A$ and $A_2 \approx 0.62 \times A$. For simplicity, let $s$ denote the effective sampling area of a single camera trap, assuming $s$ is constant constant across all traps and detectability is perfect.

First, consider the baseline scenario with no habitat selection. In this case, the probability $x$ that a specific animal is located within the detection zone of a given trap at any time $t$ is simply the ratio of the traps detection area to the total study area: $x = s/A$. Assuming $G = 100$ camera traps are randomly distributed across the area, and treating their detection zones as independent, the probability $u_n$that an animal is present within the detection zone of at one trap at time $t$ is equal to

$$u_n = 1 - (1 - x)^G$$

Given that $s$ is very small in comparison to $A$, the probability $s/A$ is typically very small. We can therefore approximate this quantity by a Taylor expansion:

$$u_n \approx G \times x \tag{3}$$

Now, consider the scenario with habitat selection. We analyze the two habitat types separately. With random placement, the expected number of camera traps falling into the selected habitat is $G_1 = G \times$

$(A_1/A)$, and similarly, $G_2 = G \times (A_2/A)$ for the non-selected habitat. Using the same rationale as before, the probability that an individual located within the selected habitat is captured by at least one trap is:

$$c_1 \approx G \times (A_1/A) \times x$$

Similarly, for an individual in the non-selected habitat:

$$c_2 \approx G \times (A_2/A) \times x$$

In the presence of habitat selection, let $p_h$ be the probability that a randomly chosen animal is located within the selected habitat. The overall probability $u_h$ that this animal is detected by at least one trap is:

$$u_h = p_h \times c_1 + (1 - p_h) \times c_2$$

Substituting the expressions for $c_1$ and $c_2$ into the equation for $u_h$, we obtain:

$$u_h = p_h \times G \times (A_1/A) \times x + (1 - p_h) \times G \times (A_2/A) \times x \tag{4}$$

Our simulation results (section S5.2.3) indicate that $p_h \approx 0.55$. We can therefore calculate the ratio of the probabilities $u_h/u_n$ by dividing equation 4 by equation 3, and substituting the parameter values from our study:

$$\begin{aligned} u_h/u_n &= (0.55 \times 100 \times 0.38 \times x + 0.45 \times 100 \times 0.62 \times x)/(100x) \\ &= (20.9 \times x + 27.9 \times x)/(100x) \\ &= 0.488 \end{aligned}$$

Therefore, the probability of an animal being detected by at least one trap is approximately halved when habitat selection is present. This reduction in detection probability explains the increased estimation variance: since the majority of traps (62%) are located in habitats that are less frequently used by the animals, the effective sampling effort is diminished compared to a spatially uniform scenario.

##### S5.4.3 A reduced variance of the trend estimation with habitat selection

The reduced variance of trend estimates is also expected. Indeed, strong habitat selection leads to a spatial aggregation of individuals within specific areas (in our simulations, the path neighborhoods). Consequently, traps located within these highly used habitats record a higher number of different animals, which increases the sensitivity of the monitoring design to detect population changes across successive years.

To formalize this, consider two years from the study period (e.g., Year 1 and Year 2), and the estimated change rate $\hat{N}_2/\hat{N}_1$. It is well-known (e.g., eq. 16 in Powell , 2007) in that the variance of this ratio can be approximated by:

$$\mathrm{Var}\left(\frac{\hat{N}_2}{\hat{N}_1}\right) \approx \frac{E(\hat{N}_2)^2}{E(\hat{N}_1)^2} \cdot \left(\frac{\mathrm{Var}(\hat{N}_2)}{E(\hat{N}_2)^2} - 2\frac{\mathrm{Cov}(\hat{N}_2, \hat{N}_1)}{E(\hat{N}_1) \cdot E(\hat{N}_2)} + \frac{\mathrm{Var}(\hat{N}_1)}{E(\hat{N}_1)^2}\right) \tag{5}$$

The critical term in this expression is the covariance $\mathrm{Cov}(\hat{N}_2, \hat{N}_1)$ between the estimators of the two years. A larger positive covariance reduces the overall variance of the ratio, thereby increasing the precision of the trend estimate.

We compared the covariance between the sampling distributions of population size estimates for Year 1 and Year 5, using the simulations conducted with and without habitat selection (sections S5.4.1 and S5.1, respectively). For example, considering the REM accounting for activity and detectability:

```r
## Covariance with habitat selection:
u <- HSRSroe[[3]]
cov(sapply(u, function(x) x[1,1]),
    sapply(u, function(x) x[5,1]))

## [1] 1164.727

## Covariance between year 1 and 5 without habitat selection
u2 <- roeNoHS[[3]]
cov(sapply(u2, function(x) x[1,1]),
    sapply(u2, function(x) x[5,1]))

## [1] 295.0406
```

The covariance between the two years is much larger when habitat selection is present, which, as explained before, is due to the aggregation of individuals in selected habitats. As a result, the changes of population size across years can be estimated more precisely.

#### S5.5 Monitoring of the large-range roe deer during five years: no habitat selection and random sampling of traps)

##### S5.5.1 With 100 traps

We also simulated the movement of an animal larger than a roe deer to assess the effect of the home-range size on the precision of our estimates. To make things clearer we considered an animal with a home-range size similar to the home-range size of the red deer. We used the same movement algorithm as for the movement of the roe deer, with the following differences: (i) the value of $\sigma^2$ used for both between-patch and patch movement was set to 10000 $m^2$, (ii) the patches location are randomly drawn in a rectangular box of 4 kilometres wide centred on the home range center. An illustration of this process is given below:

```r
simulate_large_range_roe_deer <- function()
{
    ## between 5 ànd 10 patches
    np <- sample(5:10, 1)
    ## the patches:
    cb <- cbind(runif(np,-2000,2000),
                runif(np,-2000,2000))
    ## simulation
    lipt <- list(ptg=mvtMatrices$listMatBetween[[10]],
                 ptp=mvtMatrices$listMatBetween[[10]])
    lixyt <- list(cb, cb)
    z <- roeDeerMovement(30, lixyt, lipt = lipt,
                         verbose = FALSE)

    return(z)
}

plot(simulate_large_range_roe_deer(), ty="l")
```

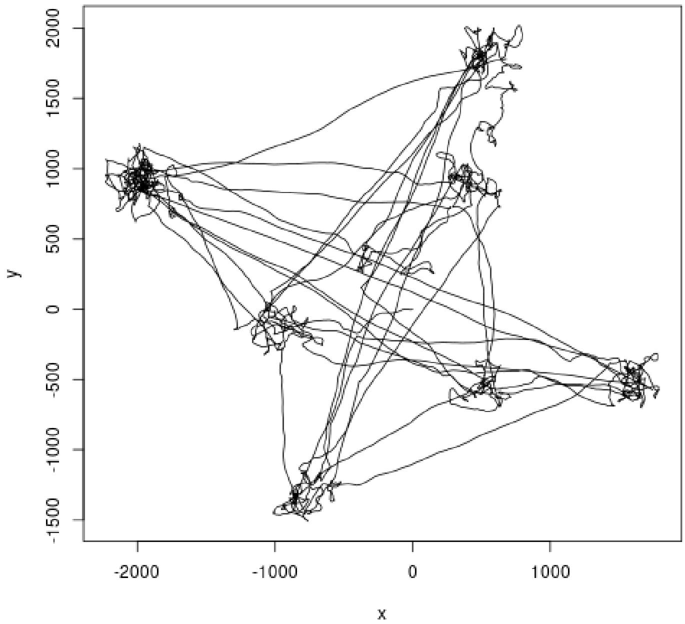

Contrary to what was simulated for the standard roe deer, we did no try to be as realistic as possible for the large-range roe deer. The aim was to generate movements that cover a large area. Simulations of this process showed that the Minimum convex polygon of the simulated animals covered a mean area equal to 469 ha (interquartile range: 383 – 548 ha), which is approximately the home range size of the red deer (Kamler et al., 2004). Note that we simulated a constant home-range size for the five years of the study (i.e., we used the same process for all years).

We then simulated a population decrease for the large-range roe deer that could also be considered as realistic for the red deer. As for the standard roe deer, we simulated a population size of 300 large-range roe deer on the study area during the first year (which would be a very high density for the red deer, but this allowed a comparison with the standard roe deer). We supposed a sex-ratio of 0.5, an adult annual survival of 0.76, and a reproduction equal to 0.5 fawns per female. We simulated this process 1000 times and chose one simulation characterized by a population decrease exactly equal to 20%:

```
set.seed(777)
li <- list()
for (k in 1:1000) {
    N <- 300
    nfem <- N/2
    nmal <- N/2
    nsurf  <- nsurm <- njfem <- njmal <- numeric(0)

    ## For each year
    for (i in 2:5) {

        ## Survival of previous year
        nsurf[i-1] <- rbinom(1,nfem[i-1],0.76)
        nsurm[i-1] <- rbinom(1,nmal[i-1],0.76)

        ## 0.5 young per female * sex ratio of 0.5 = 0.25
        njfem[i-1] <- rbinom(1,nsurf[i-1],0.25)
        njmal[i-1] <- rbinom(1,nsurf[i-1],0.25)

        ## Number of females = number surviving from previous year
```

```
        ## + new females
        nfem[i] <- nsurf[i-1]+njfem[i-1]
        ## same for males
        nmal[i] <- nsurm[i-1]+njmal[i-1]
    }
    N <- nfem+nmal
    li[[k]] <- data.frame(Ntot=N, Old=c(0,nsurm+nsurf), New=c(0,njfem+njmal))
}
liN <- do.call(rbind,lapply(li, function(x) x$Ntot))

## Repeated 1000 times, we select a case resulting in 240 animals
## during the last year:
whi <- c(1:nrow(liN))[liN[,5]==240][
    which.min(apply(apply(liN[liN[,5]==240,],1,diff),2,var))]

(dfPopSizeRedDeer <- li[[whi]])

##   Ntot Old New
## 1  300   0   0
## 2  283 222  61
## 3  267 216  51
## 4  253 208  45
## 5  240 195  45
```

Note that the resulting object `dfPopSizeRedDeer` is available as a dataset from the package (its name comes from the fact that we used the data collected on the red deer to build it).

We then used the function `simulateCTStudy5years` to simulate 1000 times a five-year camera trap study of this large-range roe deer population, simulating for each year the movement of these animals and their monitoring using 100 camera traps randomly placed on the study area. Note the use of `LRroeDeer = TRUE` in this function. WARNING: THIS CALCULATION TAKES A VERY LONG TIME!!!! (see below):

```
simrd5nohsrd <- simulateCTStudy5years(nCT = 100, duration = 30,
                                      niter = 1000,
                                      dfPopSizeRedDeer,
                                      listPatches, contourChize,
                                      contourN=northChize,
                                      contourS=southChize,
                                      dff=relationAnimalSigmaMaxd,
                                      habitatMap=habitatMap,
                                      listptg=mvtMatrices$listMatBetween,
                                      ptp=mvtMatrices$MatWithin,
                                      redDeer=TRUE, nofi = 0,
                                      backup = TRUE)
```

We did not include the result of these simulations in the package, as the resulting object size was too large to fit in a R package (2.5 Gb), though it is of course available from the authors upon request. We then used the function `process_all` of the package to estimate population size, trend and population decrease from these data. Note that this code will not work on the readers' computer, as the main object `simrd5nohsrd` is not available in the package. This code is provided for information purposes. Note also that the resulting list `LRroeNoHS` is available as a dataset in the package:

```
LRroeNoHS <- process_all(simrd5nohsrd)
```

We can use the function `present_results_CT` from the package to summarize these results:

```r
oldopt <- options(width = 100)
present_results_CT(LRroeNoHS, dfPopSizeRedDeer)
```

```
##          method Param TrueValue MeanEst     SE  SEEst SE.SEEst  Pcov  PDec PsigD
## 1       REM_raw    N1       300 207.511 40.895 40.456     4.87 0.325            
## 2       REM_raw    N5       240 165.729 33.028 32.409    4.006 0.315            
## 3       REM_raw   lam   -0.0543  -0.055  0.013  0.012    0.002 0.875     1 0.989
## 4       REM_raw    CR      -0.2    -0.2  0.047  0.046    0.009 0.885     1 0.976
## 5                                                                               
## 6       REM_det    N1       300 296.257 52.281 57.931    6.602 0.916            
## 7       REM_det    N5       240 236.537 41.961 46.383    5.297 0.916            
## 8       REM_det   lam   -0.0543  -0.055  0.013  0.012    0.002 0.874     1 0.991
## 9       REM_det    CR      -0.2    -0.2  0.047  0.046    0.009 0.937     1 0.993
## 10                                                                              
## 11 REM_det_act    N1       300 296.516 52.561 57.848    6.605 0.921            
## 12 REM_det_act    N5       240 236.556 42.166 46.411    5.377 0.916            
## 13 REM_det_act   lam   -0.0543  -0.055  0.015  0.015    0.003 0.879     1 0.968
## 14 REM_det_act    CR      -0.2    -0.2  0.057  0.054    0.011 0.931 0.999 0.964
## 15                                                                              
## 16       IS_raw    N1       300  51.406 10.502 10.236    1.301     0            
## 17       IS_raw    N5       240  41.187  8.642    8.3    1.119     0            
## 18       IS_raw   lam   -0.0543  -0.054  0.022  0.022    0.005 0.888 0.988 0.761
## 19       IS_raw    CR      -0.2  -0.195  0.086  0.084     0.02 0.872 0.982 0.704
## 20                                                                              
## 21       IS_det    N1       300 259.127 48.484 47.458    7.392 0.748            
## 22       IS_det    N5       240 208.346 39.598 39.027    6.823  0.76            
## 23       IS_det   lam   -0.0543  -0.053  0.026  0.026    0.007 0.892 0.974 0.672
## 24       IS_det    CR      -0.2   -0.19  0.101  0.099    0.034 0.889 0.959 0.611
## 25                                                                              
## 26   IS_det_act    N1       300 295.255  57.31 55.822   10.413 0.879            
## 27   IS_det_act    N5       240 237.832 47.503 46.435   10.191 0.874            
## 28   IS_det_act   lam   -0.0543  -0.053  0.031   0.03    0.008 0.893 0.957 0.583
## 29   IS_det_act    CR      -0.2  -0.187   0.12  0.117    0.045 0.897 0.936  0.51
## 30                                                                              
```

```r
options(oldopt)
```

These results are discussed in the paper.

##### S5.5.2 With 25 traps

We reproduced this simulation using only 25 traps. We again used the function `sampleCTs` from the package to subsample the results of the simulations carried out in previous section (again, this calculation will not work on the user's computer, as the dataset `simrd5nohsrd` is not available in the package. This code is provided only for information purposes):

```r
simrd5nohs25rd <- sampleCts(simrd5nohsrd, 25)
```

And we use the function `process_all` to calculate the population size and trends estimates for each one (this code is also provided for information purpose. However the resulting object `LRroeNoHS25` is available in the package as a dataset):

```r
LRroeNoHS25 <- process_all(simrd5nohs25rd)
```

And finally, we use the function `present_results_CT` from the package to summarize the study with only 25 traps:

```
oldopt <- options(width = 100)
present_results_CT(LRroeNoHS25, dfPopSizeRedDeer)

##          method Param TrueValue MeanEst      SE   SEEst SE.SEEst  Pcov  PDec PsigD
## 1       REM_raw    N1       300 210.484  80.766  78.826   19.945 0.655
## 2       REM_raw    N5       240 167.931  65.534   62.97   15.982 0.658
## 3       REM_raw   lam   -0.0543  -0.055   0.028   0.025    0.009 0.835 0.979 0.679
## 4       REM_raw    CR      -0.2  -0.197   0.103   0.105    0.058 0.846 0.967  0.63
## 5
## 6       REM_det    N1       300 299.684 104.803 113.481   27.342 0.902
## 7       REM_det    N5       240 239.059  84.642  90.663   21.937 0.895
## 8       REM_det   lam   -0.0543  -0.055   0.028   0.025     0.01 0.835 0.979 0.673
## 9       REM_det    CR      -0.2  -0.197   0.103   0.105    0.059  0.88 0.967 0.645
## 10
## 11 REM_det_act    N1       300 299.633 104.979 112.892   27.098 0.902
## 12 REM_det_act    N5       240 238.704  84.816  90.233   22.085 0.903
## 13 REM_det_act   lam   -0.0543  -0.055   0.034   0.028     0.01 0.817  0.95 0.582
## 14 REM_det_act    CR      -0.2  -0.194   0.134   0.123    0.069  0.88 0.936 0.561
## 15
## 16       IS_raw    N1       300  51.964  20.603  19.837    5.205     0
## 17       IS_raw    N5       240  41.302  16.859  15.898     4.35     0
## 18       IS_raw   lam   -0.0543  -0.056    0.05   0.043    0.017 0.828 0.878 0.423
## 19       IS_raw    CR      -0.2  -0.185   0.199   0.227    0.348 0.816  0.85 0.376
## 20
## 21       IS_det    N1       300 260.332  95.011  91.322   25.821 0.823
## 22       IS_det    N5       240 208.851  80.911  74.454   22.685 0.808
## 23       IS_det   lam   -0.0543  -0.055   0.055   0.047    0.018 0.844 0.858 0.365
## 24       IS_det    CR      -0.2  -0.178   0.224   0.249    0.382 0.841 0.834  0.34
## 25
## 26   IS_det_act    N1       300 295.397 110.777 106.145   33.628 0.862
## 27   IS_det_act    N5       240 238.746   98.29  87.876    31.69 0.854
## 28   IS_det_act   lam   -0.0543  -0.054   0.065   0.054     0.02 0.824 0.819 0.328
## 29   IS_det_act    CR      -0.2  -0.164   0.277   0.325    0.726 0.813 0.791 0.289
## 30

options(oldopt)
```

These results are discussed in the paper.

##### S5.5.3 With 100 traps and an increasing home-range

The previous simulations were carried out by supposing that the home-range size was constant across the 5 years. We conducted an additional set of simulations (with 100 camera traps) to also assess the impact of a linearly increasing home-range size for a large range species. This increase was modeled by widening the rectangular box around the home-range centroid from which patch centers were sampled. Specifically at year t, patch locations were randomly drawn within a square box of width $W_t$ centrered on the home range centroid, and defined as:

$$W_t = 2 \times \left(1.414 + t \times \frac{4 - 1.414}{5}\right)$$

where $t = 1, ..., 5$ represents the year of the study. Thus the sampling box width expanded from 2.828 km in year 1 to 4 km in year 5.

We simulated this process for 100 animals during year 1 and year 5 to illustrate it. This calculation can take several minutes, so to save some time, we have stored the results in the dataset `LRroeIS2years`:

```r
## This function simulate such movements: this function takes one
## argument si, the half-size of the box in which the patch centers
## are sampled, and returns the coordinates of the animal monitored
## during one month.
simulate_LR_deerHR <- function(si=2000)
{
    np <- sample(5:10, 1)
    ## the patches:
    cb <- cbind(runif(np,-si,si),
                runif(np,-si,si))
    ## between 5 and 10 patches
    ## simulation
    lipt <- list(ptg=mvtMatrices$listMatBetween[[10]],
                 ptp=mvtMatrices$listMatBetween[[10]])
    lixyt <- list(cb, cb)
    z <- roeDeerMovement(30, lixyt, lipt = lipt,
                         verbose = FALSE)
    return(z)
}

## Year 1
ar1 <- sapply(1:100, function(i) {
    cat(i,"\r")
    s <- simulate_LR_deerHR(si=1414)
    sr <- s[seq(1,nrow(s), length=5000),]
    pc <- mcp(SpatialPoints(as.data.frame(sr)))
    return(pc$area)
})

## Year 5
ar2 <- sapply(1:100, function(i) {
    cat(i,"\r")
    s <- simulate_LR_deerHR(si=2000)
    sr <- s[seq(1,nrow(s), length=5000),]
    pc <- mcp(SpatialPoints(as.data.frame(sr)))
    return(pc$area)
})

LRroeIS2years <- list(ar1,ar2)
names(LRroeIS2years) <- c("year 1", "year 5")
```

The dataset is directly available, and we can show how the home-range size changes between year 1 and year 5:

```r
boxplot(LRroeIS2years, xlab="Year", ylab="Home-range size (ha)")
```

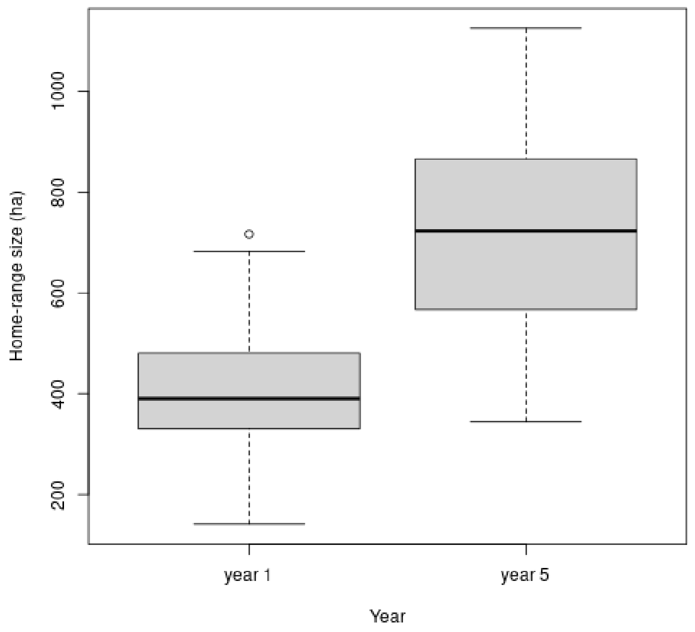

Thus, the simulated mean home-range size changes from about 400 to 700 ha in average, from year 1 to year 5.

We can simulate this increasing home-range size by setting the parameter `LRroeDeerIncreaseDV` to true. We carry out 500 simulations of a camera-trap study. Again, WARNING: THIS CALCULATION TAKES A VERY LONG TIME!!!! (see below):

```
LRDeerDVa <- simulateCTStudy5years(nCT = 100, duration = 30,
                                   niter = 500,
                                   dfPopSizeRedDeer,
                                   listPatches, contourChize,
                                   contourN=northChize,
                                   contourS=southChize,
                                   dff=relationAnimalSigmaMaxd,
                                   habitatMap=habitatMap,
                                   listptg=mvtMatrices$listMatBetween,
                                   ptp=mvtMatrices$MatWithin,
                                   LRroeDeer=TRUE, LRroeDeerIncreaseDV=TRUE, nofi = 0,
                                   backup = TRUE)
```

Again, we did not include the result of these simulations in the package, as the resulting object size was too large to fit in a R package (168 Mb), though it is of course available from the authors upon request. We then used the function `process_all` of the package to estimate population size, trend and population decrease from these data. Note that this code will not work on the readers' computer, as the main object `LRDeerDVa` is not available in the package. This code is provided for information purposes. Note also that the resulting list `LRRoeIncreasingHR` is available as a dataset in the package:

```
LRRoeIncreasingHR <- process_all(LRDeerDVa)
```

We can use the function `present_results_CT` from the package to summarize these results:

```
oldopt <- options(width = 100)
present_results_CT(LRRoeIncreasingHR, dfPopSizeRedDeer)
```

```
##         method Param TrueValue MeanEst     SE  SEEst SE.SEEst  Pcov  PDec PsigD
## 1      REM_raw    N1       300 209.485  41.76 40.032     4.69 0.312
## 2      REM_raw    N5       240 166.591  33.49 32.304    4.026  0.31
## 3      REM_raw   lam   -0.0543  -0.055  0.015  0.014    0.003 0.852     1 0.976
## 4      REM_raw    CR      -0.2  -0.203  0.055  0.051     0.01 0.862     1  0.96
## 5
## 6      REM_det    N1       300 298.666  54.27 57.261    6.522 0.906
## 7      REM_det    N5       240 237.548 43.836 46.176    5.663 0.904
## 8      REM_det   lam   -0.0543  -0.055  0.015  0.014    0.003 0.842     1 0.978
## 9      REM_det    CR      -0.2  -0.203  0.055  0.051     0.01  0.91     1 0.984
## 10
## 11 REM_det_act    N1       300 299.046 54.696 57.366    6.513 0.914
## 12 REM_det_act    N5       240 238.204 43.611 46.301    5.628 0.908
## 13 REM_det_act   lam   -0.0543  -0.055  0.017  0.016    0.003  0.85     1 0.936
## 14 REM_det_act    CR      -0.2  -0.201  0.063  0.058    0.012 0.908 0.998 0.938
## 15
## 16      IS_raw    N1       300  51.386 10.864 10.225    1.362     0
## 17      IS_raw    N5       240  41.682  8.613  8.367    1.117     0
## 18      IS_raw   lam   -0.0543   -0.05  0.023  0.023    0.004 0.892 0.978   0.7
## 19      IS_raw    CR      -0.2  -0.183  0.089  0.089     0.02 0.884 0.964 0.628
## 20
## 21      IS_det    N1       300 260.751 50.388  47.58    7.121 0.744
## 22      IS_det    N5       240 209.799 40.946 38.834    6.046 0.744
## 23      IS_det   lam   -0.0543  -0.052  0.027  0.026    0.006  0.88 0.964 0.648
## 24      IS_det    CR      -0.2  -0.189  0.103  0.101    0.031 0.898 0.952 0.584
## 25
## 26  IS_det_act    N1       300 297.232 60.094 56.095    9.945 0.838
## 27  IS_det_act    N5       240 239.579 47.976 45.909    8.249 0.866
## 28  IS_det_act   lam   -0.0543  -0.051  0.032  0.031    0.008 0.898 0.946 0.532
## 29  IS_det_act    CR      -0.2  -0.185  0.124  0.119    0.041 0.882  0.91 0.508
## 30

options(oldopt)
```

These results are consistent with those obtained under constant home-range scenarios, showing no clear differences in either bias or precision. In particular, the probabilities of detecting the decreasing trend were similar across both settings. This illustrates that, provided the mean movement speed is correctly estimated for the REM, temporal changes in home-range size do not significantly affect these estimation methods.

#### S5.6 Increasing the number of traps and shortening the study period

##### S5.6.1 Increasing the number of traps only

The simulations of the large-range roe-deer revealed that the mean number of camera traps visited by an individual is a critical driver of trend estimation precision. Increasing the home-range size was a convenient way to increase the number of traps visited per animal without altering the nominal sampling effort (number of traps or study duration). In this section, we further explore how the dimensions of sampling effort influence our results. Our large-range simulations were characterized by individuals visiting 7.6 times more traps than in the standard roe deer scenario. To test if this effect is purely additive, we ran again the simulations for the standard roe deer using 760 camera traps instead of 100. This design yields the same density of traps per home range as the large-range scenario, allowing us to determine if increasing technical effort produces results similar to those obtained through increased animal mobility.

Given that this analysis is exploratory and that increasing the number of traps to 760 significantly raises the computational time, we limited this specific scenario to 100 replications, which we considered sufficient to provide a reliable estimate of the general trend and of the magnitude of the precision improvement. The R code for these simulations is the same as for the standard roe deer simulations, just changing the number of traps. As explained before, WARNING: THIS CALCULATION TAKES A VERY LONG TIME!!!! (see below):

```
roeDeerManyTraps <- simulateCTStudy5years(nCT = 780, duration = 30,
                                          niter = 100, dfPopSize,
                                          listPatches, contourChize,
                                          contourN=northChize,
                                          contourS=southChize,
                                          dff=relationAnimalSigmaMaxd,
                                          habitatMap,
                                          listptg=mvtMatrices$listMatBetween,
                                          ptp=mvtMatrices$MatWithin, nofi = 0,
                                          backup = TRUE)
```

Again, we did not include the result of these simulations in the package, as the resulting object size was too large to fit in a R package, though it is of course available from the authors upon request. To maintain a reasonable computational timeframe, we restricted our analysis to the configurations where activity and detectability are accounted for, estimating the population sizes and trends using the REM and IS approaches. Moreover, we reduced the number of bootstrap iterations to 500 (down from 1000). This adjustment was justified by the significantly higher number of encounters and associations in this high-effort scenario (see below), which substantially increases the computational cost per iteration while simultaneously reducing the relative benefit of a higher number of bootstrap samples. Note that this code will not work on the readers' computer, as the main object `roeDeerManyTraps` is not available in the package. This code is provided for information purposes. Note also that the resulting list `res760CT` is available as a dataset in the package:

```
system.time(REM760CT <- processCTStudy(s,method="REM", Nboot=500))
system.time(IS760CT <- processCTStudy(s,method="IS", Nboot=500))
res760CT <- list(REM=REM760CT, IS=IS760CT)
```

We now present the estimation results for the population size during Year 1 and for the trend:

```
re <- do.call(rbind,lapply(res760CT, function(y) {
    popsi <- sapply(y, function(x) x[1,1])
    trend <- sapply(y, function(x) x[6,1])
    sigtrend <- mean(sapply(y, function(x) x[6,4]<0))
    dectrend <- mean(sapply(y, function(x) x[6,1]<0))
    return(c(mean(popsi), sd(popsi), mean(trend), sd(trend), dectrend, sigtrend))
}))
colnames(re) <- c("MeanPop","SEPop", "MeanTrend","SETrend", "PDec", "PsigD")
rownames(re) <- c("REM_da","IS_da")
round(re,3)

##        MeanPop  SEPop MeanTrend SETrend PDec PsigD
## REM_da 301.732 16.103    -0.060   0.021 1.00  0.85
## IS_da  296.281 20.026    -0.054   0.025 0.98  0.77
```

Comparing these results with those of the large-range roe deer simulations, we observe that the precision of the trend estimates is sligthly larger in high-effort scenario (SE $\approx$ 0.02 for the two methods, compared to 0.035 and 0.065 respectively for the REM and IS in the large-range scenario). By increasing the number of camera traps to 760, we effectively raised the mean number of visited traps per individual to 3.48 (SD = 0.17). While this value is nearly identical to that of the large-range roe deer (3.49, SD=0.62), the further gain in precision in the 760-trap scenario likely stems from the massive increase in the total number of encounters and associations. Indeed, the high-effort scenario yielded a much higher volume of data, with an average of 144914 associations (SD = 9955) compared to 17951 (SD = 3667) for the large-range roe deer. Similarly, the mean number of encounters rose to 5164 (SD = 259) compared to 1584 (SD=279) for the large-range roe deer.

Increasing the number of camera traps to 760 also led to a substantial gain in the precision of population size estimates, far exceeding that observed for the large-range roe deer. This difference is also caused by the massive increase in total sampling effort.

##### S5.6.2 Increasing the number of traps while shortening the study period

To maintain a constant total sampling effort while using 760 traps, we reduced the study period duration by 7.6. Specifically, we subsampled the data from the previous scenario, considering only the encounters and associations that occurred during the first 3.95 days instead of the full 30-day period. This approach allows us to simulate a higher trap density across the study area while keeping the total activation time (total trap-days) equivalent to our baseline 100-trap scenario.

Thus, we first selected the encounters/associations that occurred during the first 3.95 days of the study period. The following code will not work on the readers computer, as the main object `roeDeerManyTraps` is not available in the package. It is given for information:

```
## For each iteration
for (i in 1:length(roeDeerManyTraps$encounters)) {
    ## for each year
    for (a in 1:5) {
        z <- roeDeerManyTraps$encounters[[i]]$results[[a]]
        ## for each pair of animal/trap
        for (j in 1:length(z)) {
            ## keep only the encounters occurring during the
            ## first 3.95 days
            x <- z[[j]]
            y <- x[x$beginning < (3.95*3600*24),]
            z[[j]] <- y
        }
        ## store the results
        roeDeerManyTraps$encounters[[i]]$results[[a]] <-
            z[sapply(z,nrow)>0]
    }
}
```

We then use the function `process_all` to estimate the population sizes and trends from these data. Again, this code will not work on the readers' computer, as the object `roeDeerManyTraps` is not available in the package. The resulting object `roe760coef` is available as a dataset in the package:

```
roe760coef <- process_all(roeDeerManyTraps)
```

Note that the function `process_all` assumes that the study period covers 30 days, so that we have to post-process the results to account for the shorter simulated period:

```
## Post-processing
## for each estimation method
for (i in 1:length(roe760coef)) {

    ## for each iteration
    for (j in 1:length(roe760coef[[i]])) {
```

```
        ## Multiply the population size estimates by 7.6 to account
        ## for the fact that the duration was 7.6 times smaller
        x <- roe760coef[[i]][[j]]
        x[1:5,1] <- x[1:5,1]*(30/3.95)
        x[1:5,3] <- x[1:5,3]*(30/3.95)
        x[1:5,4] <- x[1:5,4]*(30/3.95)

        ## Multiply the variance of the population size estimates by
        ## 7.6^2 and updates the standard errors
        va <- x[1:5,2]^2
        vab <- va*((30/3.95)^2)
        x[1:5,2] <- sqrt(vab)

        ## trends and change rate SE are not expected to change
        ## since the popsize estimates are all multiplied by the same value
        ## so that the slope on a log scale will remain the same,
        ## and the ratio between two popsize will remain the same too.

        ## We store the results
        roe760coef[[i]][[j]] <- x
    }
}
```

We can now present the results:

```
oldopt <- options(width = 100)
present_results_CT(roe760coef, dfPopSizeRedDeer)

##          method Param TrueValue    MeanEst         SE     SEEst SE.SEEst Pcov PDec PsigD
## 1       REM_raw    N1       300 101729.111   8554.29  9107.006  885.718    0
## 2       REM_raw    N5       240  79739.894   6960.65   6885.73  633.331    0
## 3       REM_raw   lam   -0.0543     -0.058      0.025     0.026    0.002 0.93 0.99  0.67
## 4       REM_raw    CR      -0.2     -0.211      0.088     0.093    0.012 0.91 0.99  0.66
## 5
## 6       REM_det    N1       300 130229.175 10250.071 11641.913 1153.216    0
## 7       REM_det    N5       240 102103.086  8651.268  8863.083  818.035    0
## 8       REM_det   lam   -0.0543     -0.058      0.025     0.026    0.002 0.93 0.99  0.68
## 9       REM_det    CR      -0.2     -0.211      0.088     0.095    0.013 0.93 0.99  0.63
## 10
## 11 REM_det_act    N1       300 132730.406 11582.017 13048.794 1419.218    0
## 12 REM_det_act    N5       240 103665.088  9583.687  9693.383  968.447    0
## 13 REM_det_act   lam   -0.0543     -0.059      0.028     0.028    0.002 0.92 0.99  0.61
## 14 REM_det_act    CR      -0.2     -0.213      0.099     0.105    0.015 0.94 0.96  0.54
## 15
## 16      IS_raw    N1       300  23649.345  3001.701  2823.486  390.321    0
## 17      IS_raw    N5       240  19155.733  2339.428  2192.686  341.536    0
## 18      IS_raw   lam   -0.0543     -0.049      0.038     0.035    0.003 0.85 0.92  0.43
## 19      IS_raw    CR      -0.2     -0.177      0.143     0.134    0.027 0.87 0.88  0.41
## 20
## 21      IS_det    N1       300 112548.685 15774.562 14275.828  3905.58    0
## 22      IS_det    N5       240  91610.126  15280.89 11402.922 5804.354    0
## 23      IS_det   lam   -0.0543     -0.051      0.043     0.037    0.007 0.85 0.89   0.4
## 24      IS_det    CR      -0.2     -0.171      0.174     0.148    0.057 0.87 0.86  0.37
## 25
## 26  IS_det_act    N1       300  131083.64 21796.328 18453.678 5414.424    0
## 27  IS_det_act    N5       240 105760.523 20912.924 14644.891  9205.39    0
## 28  IS_det_act   lam   -0.0543     -0.053      0.049     0.041    0.007 0.82 0.89  0.33
```

```
## 29  IS_det_act    CR      -0.2     -0.174     0.201     0.166     0.076 0.85 0.83  0.33
## 30

options(oldopt)
```

The precision of the trend estimate is now nearly identical for both the high-trap-density and the large-range scenarios, confirming that when total activation time is held constant, trend precision is primarily driven by the number of traps per home range. However, the high-density design still led to an increase of the precision of population size estimates. Overall, these results confirm that achieving a high number of traps per home range is e prerequisite for reliably estimating population trends.

#### S5.7 Simulations with a wider angle for the detection zone

All the previous simulations were carried out by supposing that the detection zone was a 20 metres circular sector with an angle of 10 degrees (0.175 radians). As noted in the paper, we chose this value based on the simulations by Rowcliffe et al. (2008). However, camera traps are often characterized by a wider angle (e.g., 42 degrees, Howe et al., 2017). After setting the length and angle of the circular sector for their simulations, Rowcliffe et al. (2008) noted that *The conclusions were not sensitive to any of these variables*. We checked that this was indeed the case in this section.

We reproduced the simulations carried out in section S5.1.1, simulating 100 traps over the study area to monitor a roe deer population, without any habitat selection. We used the same code as in this section, just changing the value of the parameter `fieldangle` to an angle of 0.735 radians (42 degrees). We carried out only 500 iterations, but even with 500 iterations, WARNING: THIS CALCULATION TAKES A VERY LONG TIME!!!!

```
simrd5nohs735 <- simulateCTStudy5years(nCT = 100, duration = 30,
                                        niter = 500, dfPopSize,
                                        listPatches, contourChize, contourN=northChize,
                                        contourS=southChize,
                                        dff=relationAnimalSigmaMaxd, habitatMap,
                                        listptg=mvtMatrices$listMatBetween,
                                        ptp=mvtMatrices$MatWithin,
                                        fieldangle=0.735, depth=20,
                                        nofi = 0, backup = TRUE)
```

We did not include the result of these simulations in the package, as the resulting object size was too large to fit in a R package (2.5 Gb), though it is of course available from the authors upon request. We then used the function `process_all` to estimate the population abundance for each year, the variation rate between year 1 and year 5, and the trend $\lambda$ (together with their 90% bootstrap confidence interval). We use the function `process_all` (again, recall that this code will not work since the main object `simrd5nohs735` is not available in the package. This code is provided only for information purposes):

```
roeNoHS735 <- process_all(simrd5nohs735)
```

The resulting object `roeNoHS735` is available as a dataset of the package for the 1000 simulations:

```
str(roeNoHS735,1)

## List of 6
##  $ REM_raw          :List of 500
##  $ REM_detect       :List of 500
##  $ REM_detect_active:List of 500
##  $ IS_raw           :List of 500
```

```
##   $ IS_detect        :List of 500
##   $ IS_detect_active :List of 500
```

We can then build the table summarizing the main results of these simulations. The function `present_results_CT` from the package can be used:

```
oldopt <- options(width = 100) ## to allow a cleaner display of the results

present_results_CT(roeNoHS735, dfPopSize)

##          method Param TrueValue MeanEst     SE  SEEst SE.SEEst  Pcov  PDec PsigD
## 1       REM_raw    N1       300 245.167 42.068 39.617    6.981 0.602
## 2       REM_raw    N5       239  189.67  32.53 32.797     5.68 0.558
## 3       REM_raw   lam   -0.0547  -0.061  0.049  0.048    0.007 0.884 0.894  0.37
## 4       REM_raw    CR   -0.2033  -0.209  0.166  0.165     0.04 0.874 0.892  0.37
## 5
## 6       REM_det    N1       300 305.867  49.15 49.461    8.515 0.892
## 7       REM_det    N5       239 236.566 37.588 40.893    6.881 0.908
## 8       REM_det   lam   -0.0547  -0.061  0.049  0.048    0.007 0.882 0.894 0.374
## 9       REM_det    CR   -0.2033  -0.209  0.166  0.168    0.039  0.91 0.892 0.318
## 10
## 11 REM_det_act    N1       300 304.132 50.146 50.393    8.774 0.896
## 12 REM_det_act    N5       239 235.869 38.112 41.416     6.99 0.912
## 13 REM_det_act   lam   -0.0547   -0.06   0.05  0.049    0.007 0.884 0.888 0.348
## 14 REM_det_act    CR   -0.2033  -0.206  0.173  0.174    0.042  0.92 0.886  0.29
## 15
## 16       IS_raw    N1       300  55.674  10.23  9.745    1.828     0
## 17       IS_raw    N5       239  43.205  8.176  8.121    1.531     0
## 18       IS_raw   lam   -0.0547   -0.06  0.054  0.052    0.007 0.886  0.87  0.34
## 19       IS_raw    CR   -0.2033  -0.203  0.188  0.186     0.05 0.882 0.868  0.35
## 20
## 21       IS_det    N1       300  265.66  45.78 44.305    8.981 0.774
## 22       IS_det    N5       239 208.111 35.138 36.544    6.607 0.758
## 23       IS_det   lam   -0.0547  -0.057  0.051  0.051    0.007   0.9  0.88 0.298
## 24       IS_det    CR   -0.2033  -0.196  0.185  0.185    0.049 0.892  0.86 0.318
## 25
## 26   IS_det_act    N1       300  301.51 54.018 53.033   12.349 0.882
## 27   IS_det_act    N5       239 236.034 42.249 43.054    8.477 0.886
## 28   IS_det_act   lam   -0.0547  -0.057  0.053  0.054    0.007 0.882 0.864 0.286
## 29   IS_det_act    CR   -0.2033  -0.195  0.193  0.197    0.055 0.906 0.858 0.276
## 30

options(oldopt)
```

On could compare this table with the one obtained with camera traps characterized by a detection zone of 0.175 radians (10 degrees). Though this table is already presented in section S5.1.1, we show it again below to facilitate this comparison.

```
oldopt <- options(width = 100) ## to allow a cleaner display of the results

present_results_CT(roeNoHS, dfPopSize)

##          method Param TrueValue MeanEst     SE  SEEst SE.SEEst  Pcov  PDec PsigD
## 1       REM_raw    N1       300  237.31 37.875 38.687    6.215 0.511
## 2       REM_raw    N5       239 181.767 33.792 31.553    6.114 0.444
```

```
## 3      REM_raw   lam  -0.0547  -0.063  0.049  0.048    0.007 0.869 0.906 0.364
## 4      REM_raw    CR  -0.2033   -0.22  0.165  0.165    0.043 0.859 0.906 0.378
## 5
## 6      REM_det    N1      300  303.86  45.68  49.51    7.853 0.923
## 7      REM_det    N5      239 232.533 40.045 40.352    7.572 0.882
## 8      REM_det   lam  -0.0547  -0.063  0.049  0.048    0.007 0.867 0.906 0.367
## 9      REM_det    CR  -0.2033   -0.22  0.165  0.169    0.042   0.9 0.906 0.327
## 10
## 11 REM_det_act    N1      300 303.224 47.154 50.959    8.427 0.928
## 12 REM_det_act    N5      239 232.706 41.018 41.146     7.83 0.882
## 13 REM_det_act   lam  -0.0547  -0.062  0.051   0.05    0.007 0.867 0.889 0.344
## 14 REM_det_act    CR  -0.2033  -0.217  0.172  0.176    0.045 0.904 0.891 0.311
## 15
## 16       IS_raw   N1      300  54.903  9.955 10.084    1.897     0
## 17       IS_raw   N5      239  42.983  8.975    8.3    1.844     0
## 18       IS_raw  lam  -0.0547  -0.058  0.055  0.055    0.008  0.88 0.843 0.289
## 19       IS_raw   CR  -0.2033  -0.198  0.195  0.198    0.057  0.87 0.848 0.279
## 20
## 21       IS_det   N1      300 262.512 47.812 46.688   11.689 0.737
## 22       IS_det   N5      239 205.374 40.169 37.611    9.195 0.682
## 23       IS_det  lam  -0.0547  -0.057  0.056  0.054    0.008 0.884 0.854 0.279
## 24       IS_det   CR  -0.2033  -0.197  0.196  0.197    0.056 0.882 0.841 0.276
## 25
## 26   IS_det_act   N1      300 298.869 57.784 56.794   15.407 0.881
## 27   IS_det_act   N5      239  233.97 49.233 45.015   12.228 0.849
## 28   IS_det_act  lam  -0.0547  -0.057  0.059  0.057    0.009 0.885 0.833 0.272
## 29   IS_det_act   CR  -0.2033  -0.193  0.211  0.214    0.067 0.891 0.834 0.257
## 30

options(oldopt)
```

**The results are not sensitive to the angle of the detection zone.**

#### S5.8 Remark 1: On the bias of trend estimation

In all our simulations, we used equation 1 to estimate $\lambda$. We just replaced $N_t$ by its estimate $\hat{N}_t$, i.e. we estimated $\mu$ with:

$$\hat{\mu} = \frac{\mathrm{Cov}(\log \hat{N}_t, t)}{\mathrm{Var}(t)}$$

However, we can show that $\mathrm{Cov}(\log \hat{N}_t, t)$ is a biased estimate of $\mathrm{Cov}(\log N_t, t)$. Without loss of generality, let us replace the variable $t$ in this equation by a centred variable $t'$ (i.e. with a mean equal to zero), i.e. $t' = t - (T+1)/2$. Then:

$$\begin{aligned}
\mathrm{Cov}(\log \hat{N}_t, t') &= \frac{1}{T}\sum_{t=1}^{T} t'(\log \hat{N}_t - \bar{n}) \\
&= \frac{1}{T}\left\{\sum_{t=1}^{T} t' \log \hat{N}_t\right\} - \frac{1}{T}\bar{n}\left\{\sum_{t=1}^{T} t - (T+1)/2\right\} \\
&= \frac{1}{T}\left\{\sum_{t=1}^{T} t' \log \hat{N}_t\right\} 
\end{aligned} \tag{6}$$

with $\bar{n} = \frac{1}{N}\sum_{t=1}^{T} \log \hat{N}_t$, the mean of $\log \hat{N}_t$ over the study period. Note that the second term in the second row vanishes since:

$$\begin{aligned}\left\{\sum_{t=1}^{T} t - (T+1)/2\right\} &= \left\{\sum_{t=1}^{T} t\right\} - T(T+1)/2 \\ &= T(T+1)/2 - T(T+1)/2 = 0\end{aligned}$$

We use a Taylor expansion of the function:

$$f(\mathbf{x}) = \frac{1}{T}\left\{\sum_{t=1}^{T} t' \log x_t\right\}$$

The gradient vector of this function is equal to:

$$\frac{\partial f(\mathbf{x})}{\partial x_t} = \frac{t'}{T \times x_t}$$

And the Hessian matrix:

$$\frac{\partial f(\mathbf{x})}{\partial x_t \partial x_u} = \begin{cases} 0 & \text{if } t \neq u \\ -\frac{t'}{Tx_t^2} & \text{otherwise} \end{cases}$$

The Taylor approximation around the true value $\mathbf{x} = \mathbf{N} = \{N_t\}$ is:

$$\mathrm{Cov}(\log \hat{N}_t, t') = \frac{1}{T}\left\{\sum_{t=1}^{T} t' \log N_t\right\} + \left\{\sum_{t=1}^{T} \frac{t'(\hat{N}_t - N_t)}{TN_t}\right\} - \left\{\sum_{t=1}^{T} \frac{t'(\hat{N}_t - N_t)^2}{2TN_t^2}\right\}$$

We now define the bias:

$$B = E\left(\mathrm{Cov}(\log \hat{N}_t, t') - \mathrm{Cov}(\log N_t, t')\right)$$

The bias can then be estimated with:

$$\begin{aligned}B &= E\left(\sum_{t=1}^{T} \frac{t'(\hat{N}_t - N_t)}{TN_t}\right) - E\left(\frac{t'(\hat{N}_t - N_t)^2}{2TN_t^2}\right) \\ &= \sum_{t=1}^{T} \frac{t'E(\hat{N}_t - N_t)}{TN_t} - \sum_{t=1}^{T} \frac{t'E\left((\hat{N}_t - N_t)^2\right)}{2TN_t^2} \\ &= \sum_{t=1}^{T} \frac{-t'}{T} \times \frac{\mathrm{Var}(\hat{N}_t)}{2N_t^2} \\ &= \frac{1}{2}\sum_{t=1}^{T} \frac{((T+1)/2) - t}{T} \times \frac{\mathrm{Var}(\hat{N}_t)}{N_t^2}\end{aligned}$$

Note that the first term of the equation on the second row vanishes, as under the hypothesis of zero bias of the estimation of $N_t$ with $\hat{N}_t$, we have $E(\hat{N}_t - N_t) = 0$.

Note that the ratio $\mathrm{Var}(\hat{N}_t)/N_t^2$ in this equation is an approximation of the variance of $\log \hat{N}_t$. Indeed, a first order Taylor approximation of $\log X$ around $E(X)$ shows that:

$$\log X \approx \log E(X) + \frac{(X - E(X))}{E(X)}$$

So that:

$$\mathrm{Var}(\log X) \approx \frac{\mathrm{Var}(X)}{E(X)^2}$$

And therefore:

$$\mathrm{Var}(\log \hat{N}_t) \approx \frac{\mathrm{Var}(\hat{N}_t)}{N_t^2}$$

So that we can interpret equation 7:

- When the variance of $\log \hat{N}_t$ is constant for all $t$, then the ratio $\text{Var}(\hat{N}_t)/N_t^2$ is equal to a constant $k$ that does not depend on $t$, and equation 7 is therefore equal to $(k/T)\sum((T+1)/2)-t)=0$. In other words, when the variance of $\log \hat{N}_t$ is constant for all $t$, the slope estimation is not biased.

- When the variance of $\log \hat{N}_t$ is higher at the beginning of the period, i.e. for small $t$, (i.e. for positive values of $(T+1)/2-t$, the bias $B$ is greater than zero: the slope $\mu = \log \lambda$ is overestimated (we overestimate population increase and underestimate population decreases).

- Conversely, when the variance of $\log \hat{N}_t$ is higher at the end of the study period, i.e. for negative values of $(T+1)/2-t$, the bias is negative: we underestimate $\mu$ and therefore population increase (and we overestimate population decrease).

We can rely on the simulations carried out previously to calculate the variance of $\log \hat{N}_t$ – estimated for example using raw IS for all years, with the two simulations protocols (with and without habitat selection):

```
u <- sapply(roeNoHS$IS_raw, function(x) log(x[1:5,1]))
sd1 <- apply(u,1,var)
sd2 <- apply(sapply(roeHSBiasedSam$IS_raw, function(x) log(x[1:5,1])),1,var)
dv <- data.frame(year=paste0("Year ", 1:5),
                 Variance1=sd1,
                 Variance2=sd2)
names(dv) <- c("Year","Without habitat selection","With habitat selection")
dv

##     Year Without habitat selection With habitat selection
## 1 Year 1                0.03458911             0.03058645
## 2 Year 2                0.04238270             0.02743642
## 3 Year 3                0.03849385             0.02980936
## 4 Year 4                0.03973360             0.02701718
## 5 Year 5                0.04460680             0.02703717
```

The variance of $\log \hat{N}$ is weakly variable from one year to the next in both cases, suggesting a limited bias when a simple unweighted linear regression is used to estimate $\lambda$. Working on a logarithmic scale to estimate this parameter actually makes the variance of $\log \hat{N}$ more stable, and allows this limited bias.
